# Incoherent feedback from coupled amino acids and ribosome pools generates damped oscillations in growing E. coli

**DOI:** 10.1101/2023.10.25.563923

**Authors:** Rossana Droghetti, Philippe Fuchs, Ilaria Iuliani, Valerio Firmano, Giorgio Tallarico, Ludovico Calabrese, Jacopo Grilli, Bianca Sclavi, Luca Ciandrini, Marco Cosentino Lagomarsino

**Affiliations:** IFOM - Istituto Fondazione di Oncologia Molecolare, Milan, Italy; Centre de Biologie Structurale (CBS), Université de Montpellier, CNRS, INSERM, Montpellier, France; Laboratory of Computational and Quantitative Biology (LCQB), Sorbonne Université, Institut de Biologie Paris-Seine, Paris, France; Dept. of Computational Biology, University of Lausanne, Lausanne, Switzerland; Dipartimento di Fisica, Università degli Studi di Milano, Milan, Italy; Quantitative Life Science, The Abdus Salam International Center for Theoretical Physics, Trieste, Italy; Institut Universitaire de France; INFN - Istituto Nazionale Fisica Nucleare sezione di Milano, Milan, Italy

## Abstract

Current theories of bacterial growth physiology demonstrate impressive predictive power but are often phenomenological, lacking mechanistic detail. Incorporating such details would significantly enhance our ability to predict and control bacterial growth under varying environmental conditions. The “Flux Controlled Regulation” (FCR) model serves as a reference framework, linking ribosome allocation to translation efficiency through a steady-state assumption. However, it neglects ppGpp-mediated nutrient sensing and transcriptional regulation of ribosomal operons. Here, we propose a mechanistic model that extends the FCR framework by incorporating three key components: (i) the amino acid pool, (ii) ppGpp sensing of translation elongation rate, and (iii) transcriptional regulation of protein allocation by ppGpp-sensitive promoters. Our model aligns with observed steady-state growth laws and makes testable predictions for unobserved quantities. We show that during environmental changes, the incoherent feedback between sensing and regulation generates oscillatory relaxation dynamics, a behavior that we support by new and existing experimental data.

## I. INTRODUCTION

The regulation of growth is critical for all living cells [1–4]. On general grounds, it can be seen on different levels as a global resource allocation problem whereby enzymes and proteins that mediate the fluxes are produced [5], and the coordination of sensing of nutrients and other environmental cues leading to the regulatory circuits [6, 7]. The resource allocation problem determines the target levels of protein expression that results in the organism’s growth rate (or more in general the fitness) in a given environment and also in possibly fluctuating growth conditions [5, 8, 9]. For example, under carbon-limited growth, there is a trade-off between the expression of ribosomes, which carry out protein biosynthesis, and the metabolic enzymes providing the necessary amino acids and other precursors [5, 10–12]. The theories that originate from this observation work are conceptually powerful and quantitatively predictive both during exponential growth [10] and out-of-steady-state scenarios [13].

On the other hand, the current frameworks miss an explicit description of the regulatory circuits that coordinate the cellular response to perturbations by reading the environmental signals. In bacteria, the circuit that implements this control is based on the small signaling molecule (p)ppGpp (guanosine tetraphosphate or pentaphosphate) [6, 7]. This small molecule can “read” the external environment by sensing changes in the uncharged tRNA caused by changes in amino-acid abundances or other essential nutrients. Under scarcity of these components, uncharged tRNA molecules accumulate in the ribosome, leading to the activation of the ribosome-associated RelA protein, which synthesizes ppGpp by transferring a pyrophosphate group from ATP to GTP or GDP, resulting in the production of ppGpp and AMP or ADP [14, 15]. The less-well-characterized SpoT protein can both degrade and catalyze the synthesis of ppGpp, which provides a mechanism for fine-tuning the cellular response to stress. Recent quantitative measurements of ppGpp levels during nutrient shifts lead to the hypothesis that ppGpp amounts may sense translation elongation speed through the concerted action of RelA and SpoT [16]. The sensing of amino acid levels by ppGpp results in the regulation of ribosomal biosynthesis through its function as a signaling molecule that modulates the activity of RNA polymerase on ribosomal and growth-related promoters. This process involves the DksA protein and a GC-rich “discriminator region” present in the promoter sequences and affects the relative amount of transcripts [7]. As ppGpp levels increase, the production of ribosomal transcripts decreases, enabling bacteria to adjust to nutrient and stress conditions by redirecting resources towards survival and growth [6, 17, 18].

Being able to predict how cells will respond to new perturbations is crucial, and to this end a mechanistic understanding of growth control is essential. It is important to note that the response to a change is specific to the environment or perturbation under investigation [12, 19], while the regulatory mechanism remains the same irrespective of the environment [7, 16], albeit different perturbations can trigger the response of different regulatory circuits. However, obtaining a detailed description of the key circuits controlling resource allocation and growth is challenging as we still lack a comprehensive understanding of all the molecular players. A crucial problem for gaining insight into the underlying sensing and regulation of growth is that during steady-state (i.e., balanced exponential) growth all the relevant molecular players are balanced [4, 20]. In such conditions, even if resource allocation is the result of the action of sensing and regulatory circuits, the mechanistic principles and causal chains governing these links remain hidden. Thus, to understand the regulatory aspects, it is necessary to study the dynamic cellular response to perturbations and focus on the out-of-steady-state behavior. From a modeling perspective, non-steady conditions offer the opportunity to describe jointly growth laws, limiting components, and the role of the external nutrients and cues on cellular growth [9, 13, 21, 22]. Here we focus on perturbations performed by changing the external nutrient source [13, 23]. A number of modeling studies have focused on non-steady conditions, and they can be divided into models that incorporate the growth law theory by using a top-down approach [13, 24, 25], and bottom-up models with a more detailed descriptions of the regulatory mechanisms [9, 17, 21]. Each one of these models makes different modeling assumptions, which we will discuss in more detail below. Recently, Wu and coworkers [16] have studied how the ppGpp regulatory mechanism can sense the elongation rate. However, this study does not include a description of the connection between the environment and the translational speed, which is mediated by the pool of amino acids available for protein synthesis.

In this study, we propose an intermediate approach between a top-down framework and a specific model of the circuits by introducing a comprehensive model that incorporates (I) an explicit description of amino acid sensing, (II) a detailed account of the mechanistic regulation of transcription via ppGpp, and (III) a framework for growth laws and global resource allocation. Notably, our model manifests an emergent property whereby the system’s response to external perturbations exhibits oscillatory behavior, which arises due to the incoherent mutual feedback loop that emerges between amino acid pools and ribosome levels. The key ingredient for this property to arise is the joint description of the amino acids and ribosomes pool in a dynamical framework, which is not addressed in previous frameworks [5, 13].

## II. RESULTS

### A theoretical model to describe response to nutrient changes

To describe the out-of-steady-state dynamics of cell growth, we designed a model framework that, starting from the framework defined in ref. [13] takes into account all the major mechanistic players involved in the response to the internal amino acids concentration, translation rate, nutrient sensing by ppGpp concentration, mRNA dynamics, and protein production. Fig. 1 shows the scheme of the proposed framework, which we describe in the following paragraphs (see Methods and Supplementary Note 1 and 2 for details).

**FIG. 1:**
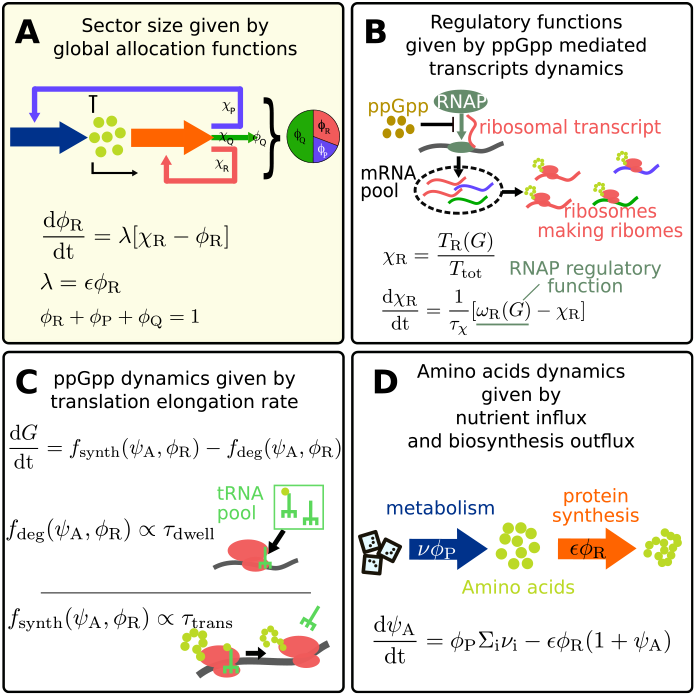
Mechanistic model for bacteria response to external perturbations. Global scheme of the regulatory model. The out-of-steady-state dynamics is governed by four main ingredients: (A) the allocation functions *χ*_i_s, which set the target size for the proteome sectors *ϕ*_i_s (B) the ppGpp-mediated transcript dynamics (total transcript *T* and ribosomal transcripts *T*_R_), which determine the values of the allocation functions by setting the composition of the transcript pool (C) the ppGpp dynamics, which reads the translation elongation rate *E*-determined by the dwelling and translocation time- and regulates the transcript production (D) the production and consumption of amino acids, *ψ*_A_, which control ppGpp production by setting the translation speed. The first module on the dynamics of the sector size is derived from ref. [13]. Each box contains the equation associated to the illustrated mechanism (described in the main text).

In the model, ribosomes are responsible for the synthesis of all proteins, divided into “sectors” representing groups of co-regulated proteins (which usually have similar functions) [13]. The model focuses on three sectors: ribosomal (*ϕ*_R_), constitutive (*ϕ*_P_, assumed to be connected with the flux of amino acids [10] and metabolism in general), and housekeeping (*ϕ*_Q_, assumed to be kept homeostatically constant [10]). Note that the model is is a coarse-grained description of key growth-related cellular processes [10, 12]; for example, the R sector includes ribosomes but also translation-related proteins, and the P sector includes catabolism (e.g. carbon uptake) and anabolism (e.g., amino acids synthesis). Out of steady state, the target size of each sector is regulated by “regulatory functions” [13], denoted as *χ*_R_, *χ*_P_, and *χ*_Q_, representing the fraction of ribosomes that actively translate proteins of the different sectors. From the definition of sectors, *ϕ*_i_ = *M*_i_*/M*_tot_ and the repartition of the total biosynthesis flux, one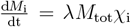, obtains by chain rule the following dynamical equation for the sector size [13]

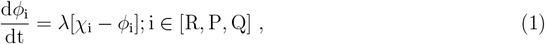

where *λ* is the growth rate (Fig. 1). This equation essentially states that (I) ribosomes translate sector i with allocation function *χ*_i_, and (II) that the equilibrium state of the sector is *ϕ*_i_ = *χ*_i_. In the absence of post-transcriptional control [6, 26], we assume that ribosomes randomly attach to transcripts, initiate protein synthesis, and produce the proteins. Therefore, the composition of the mRNA pool determines the redistribution of the flux among the three sectors. We hence define *χ*_R_ as the ratio of ribosomal transcripts (*T*_R_) to the total number of transcripts (*T*), and similarly for *χ*_P_ and *χ*_Q_.

The ribosomal mRNA pool composition is regulated by the alarmone ppGpp, which controls the partitioning of the RNA polymerase (RNAP) [7]. ppGpp is primarily responsible for the regulation of ribosomal rRNA, which usually represents the limiting step for ribosomal formation [27]. However, previous studies have shown that ribosomal proteins are also under the ppGpp-DksA regulation, in addition to the post-transcriptional control asserted by rRNA concentration [28, 29]. The combined effect of translational and post-translational regulation produces the well-known relation between ppGpp levels and ribosomal mass fraction. In this model, since we do not include explicitly the ribosomal RNA, we describe the ppGpp regulation as a transcriptional effect on just the ribosomal proteins. Consequently, the number of ribosomal transcripts, *T*_R_, depends on the concentration of ppGpp, denoted as *G*, via the partition of the RNA polymerases, denoted by *ω*_R_. We can derive an equation describing the dynamical change of *χ*_R_ after an environmental nutrient shift. This is achieved by integrating our definition of *χ*_R_ — as the ratio of ribosomal-protein transcripts— with the transcripts’ dynamics, assuming they are generated by the available RNAP according to the RNAP partition *ω*_R_ and that they degrade at a constant rate (see Supplementary Note 2 for the detailed equations). The resulting equation can be expressed as

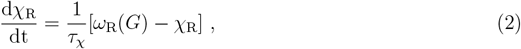

where *τ*_*χ*_ is a time-scale parameter that represents the time needed for the transcript pool to change after the shift. Assuming that the concentration of total transcripts remains constant across the shift, this time scale coincides with the mRNA half-life, which we set to be around 1 min [26] (Fig. 1, Methods). This adaptation time scale is due to the fact that the production of new transcripts is not instantaneous. In our model the partitioning function of the RNA polymerases *ω*_R_ depends solely on the ppGpp pool, and its functional form derives from a fit of the steady-state data (see Supplementary Fig. 1). This assumption is based on the recent study by Balakrishnan and coworkers [26], who found that the mRNA pool composition across conditions is mostly determined by the specific gene on-rates and only depends weakly on other factors such as gene dosage or mRNA degradation. This is especially true for the proteins belonging to the ribosomal sector, whose genes on-rates are governed by ppGpp. An extension of our theory including other dependencies is straightforward. Note that the direct ppGpp regulation of the ribosomal genes introduces also a passive ppGpp regulation of catabolic and anabolic genes, exerted by the genes competition for transcriptional resources. In this context, we can define 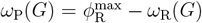. Previous studies [30, 31] show that the ppGpp effect on transcription is more complicated than the simple ribosomal transcription inhibition and report a direct up-regulation of amino acid promoters by the coordinated action of ppGpp and DksA. However, this effect appears to be prominent when cells face amino acid starvation [30], a very different condition from the nutrient upshift studied here.

As found in ref [16], ppGpp levels are directly connected to the translation elongation rate. This quantity reflects the amount of charged tRNAs and other limiting factors that are available for translation. Therefore, to close our model we need to address in a simplified way the dynamics of the amino acids pool. Amino acid levels are determined in our model by the interplay of nutrient uptake, represented by the uptake flux *νϕ*_P_, biosynthesis, represented by the biosynthesis flux *ϵϕ*_R_, and volume growth, which contributes with a dilution term *λψ*_A_ (Fig. 1). Specifically, the model describes the abundance of one compound amino acid species (related to tRNA charging, see below) by the equation

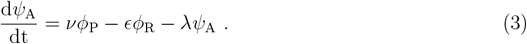

Eq. (3) introduces the normalized amino acid mass *ψ*_A_, which is the ratio of amino acid mass (*A*) to total protein mass (*M*_tot_). The catabolic flux linking nutrients to amino acids is represented by the term *νϕ*_P_, where *ν* denotes the nutrient quality (an average catabolic flux per employed catabolic sector protein), and the biosynthesis flux is represented by *ϵϕ*_R_. The last term accounts for the dilution effect caused by volume growth, which we chose to not neglect, as this term has an impact on the time scales of the relaxation dynamics (see Supplementary Fig. 2).

### Linking amino acid pool, ppGpp and global transcription

A key aspect of our model is the explicit representation of the amino acid pool (which represents a significant departure from ref. [13]). This ingredient plays a vital role in the sensing mechanism of ppGpp, which governs the cellular response to perturbations. Indeed, following a shift in nutrient availability, the first change observed by the cell is in the metabolic flux *νϕ*_P_, due to alterations in nutrient quality (*ν*). Consequently, these changes in nutrient availability lead to variations in the levels of amino acids, which have a direct influence on tRNA charging and the translation rate. In turn, the altered translation rate induces changes in the level of ppGpp (*G*), thereby triggering a transcriptional reconfiguration of the cell’s allocation strategy. Let us further explain the relationship between the translation elongation rate *ϵ* and *ψ*_A_: based on experimental findings (ref. [32]), we express the translation elongation rate *(ϵ*) as the following function of the concentration of charged tRNAs

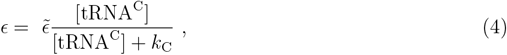

where 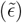 is the theoretical maximum value of the elongation rate and *k*_C_ sets a sensitivity scale. The concentration of charged tRNAs is, in turn, influenced by the pool of available amino acids, which affects the dynamics of tRNA charging. We assume a simple relationship between uncharged tRNAs and cognate amino acids: [tRNA^C^] *∝ ψ*_A_, i.e. that the fast time-scale changes of precursors are instantaneously mirrored by tRNA charging. For a detailed motivation of this assumption please refer to the Supplementary Note 2. Following this assumption we write

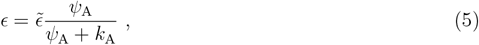

where the scale *k*_A_ is the analogue of *k*_C_ for this pool. This explicit (albeit simplified) description of charged tRNA sensing in our model allows us to make quantitative predictions regarding the size of the amino acid pool and its relationships with other variables (Fig. 2).

**FIG. 2:**
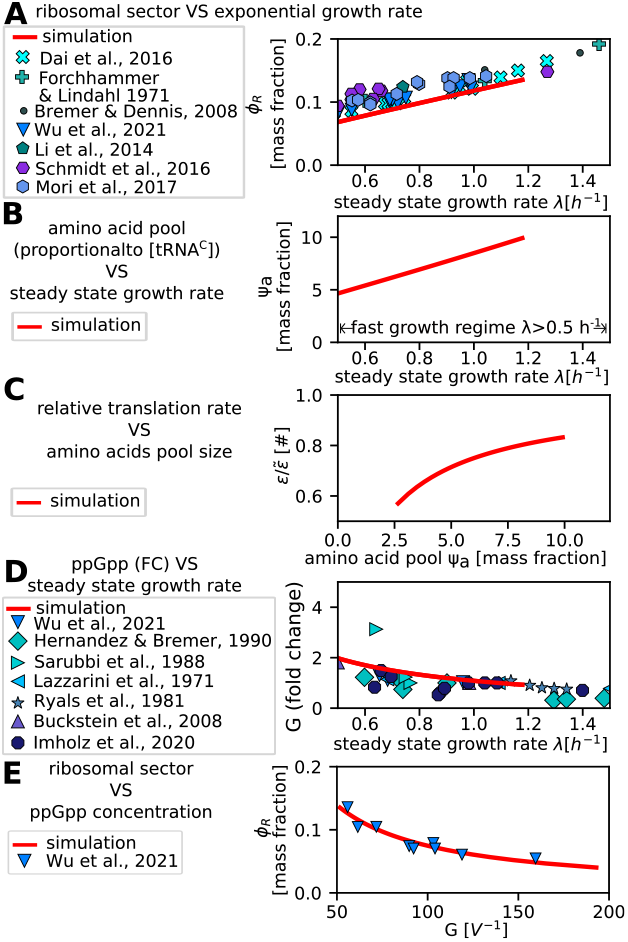
The proposed framework predicts steady-state relationships between the amino acids pool, ppGpp levels and ribosomal allocation, and reproduces the available experimental data. **A:** Model prediction of the dependence of the ribosomal sector fractional size (y-axis) on the exponential growth rate (x-axis). Simulations (red solid line) are compared with experimental data from various studies (ref.s [16, 20, 24, 32, 52–54], blueish points). **B:** The plot shows the simulated model prediction (red solid lines) for the relationship between the steady-state exponential growth rate (*x* axis) and the size (mass fraction *ψ*_A_) of the amino-acid pool (*y* axis). This quantity is proportional to the concentration of charged tRNA ([tRNA^C^]), as explained in the text and in Supplementary Notes 2. **C:** Model prediction (red solid line) for the relationship between the size of the amino-acid pool (*x* axis) and the relative translation rate 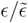 (*y* axis) in steady-state growth. **D:** The model prediction (red solid line) for the relationship between the steady-state exponential growth rate (*x* axis) and the ppGpp concentration (*y* axis) is in line with the experimental data from various studies (ref.s [16, 55–60], blueish points). **E:** The model prediction (red solid line) for the relationship between the ppGpp concentration (*x* axis) and the size of the ribosomal sector (*y* axis) agrees with the available experimental data (blue reverse triangles, data from ref. [16]). For this figure, we restricted ourselves to the fast-growth regime (*λ >* 0.5*h*^*−*1^). For the slow growth regime, where additional phenomena such as degradation and inactive ribosomes also impact physiology, please refer to Supplementary Fig. 3.

Lastly, in order to connect ppGpp levels to the amino-acid pool, we use the model proposed by Wu and coworkers [16], who established that ppGpp level (*G*) is related to charged tRNA levels in a way that ppGpp is effectively a function of translation elongation rate (*ϵ*) through the equation

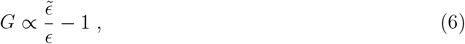

which also means that *G ∝ k*_A_*/ψ*_A_ = *k*_C_*/*[tRNA^C^].

It is important to note that so far *ϵ* solely represents the translation elongation rate. In the presence of stalled ribosomes (e.g., under chloramphenicol treatment [32, 33]) this is distinct from the average ribosomal activity (called *σ* in ref [13] and defined as *λ/ϕ*_R_). The relationship between *G* and tRNA charging is explained in terms of the typical times elongating ribosomes spend in elongation (*τ*_trans_) and waiting for a charged tRNA (*τ*_dwell_), both quantities that are captured by the translation elongation rate *ϵ*, see Fig. 1 and ref. [16]. Note that the data gathered by Wu and coworkers is in contrast with previous models that assumed that ppGpp concentration depends on both ribosomal sector size and amino acid levels [9, 17, 21]. It is also worth mentioning that our model is not the sole framework that describes sensing. The Flux Parity Model (FPM), presented by Chure and Cremer [25], also incorporates a (different) relationship between tRNAs and ppGpp.

### The model reproduces steady-state resource allocation data

Two aspects of our model are noteworthy. Firstly, the ability to reproduce the known steadystate relationships between measured quantities is a crucial requirement, and our model satisfies this criterion. Our model, with respect to the original FCR framework from which is derived, incorporates additional observables, such as ppGpp and amino acid levels. ppGpp levels have been measured and they have been incorporated in our model. Furthermore, the model’s ability to predict the amino acid pool levels is an outcome that can be tested by new experiments, making it a testable prediction.

Fig. 2 shows the steady state results of the simulations of the model, compared with experimental data presented in various studies ref.s [16, 32] TOADD. The simulations in Fig. 2 start from an arbitrary initial condition and collect the steady-state values of the main state variables once the system has reached equilibrium. Panels B and D of the figure show the dependency between the growth rate, the amino-acid pool *ψ*_A_, ppGpp concentration *G*, and are new predictions of our model. Panel C shows the dependence of the translation elongation rate *ϵ* on the amino-acid pool, which derives from our eq. 5, and panel D shows the relationship between ribosomal allocation *ϕ*_R_ and ppGpp levels, fitted from the data (see Supplementary Fig. 1). As we anticipated above Fig. 2 illustrates that the model can accurately reproduce available steady-state data (from ref. [16]) for ppGpp.

In order to avoid unnecessary complications, we restricted this analysis to a “fast-growth” regime, which we defined by the condition *λ >* 0.5*h*^*−*1^. Indeed, it is well known that in slowgrowth conditions, other phenomena such as protein degradation and inactive ribosomes play a significant role in the growth physiology [32, 33]. The model discussed in the main text of this work does not account for these phenomena and therefore refers to the fast-growth regime. We have also studied an extended version of our framework that includes the essential ingredients to describe the slow-growth behavior. For a description of this version of the model and its steady-state predictions please refer to Supplementary Note 3 and Supplementary Fig. 3.

Finally, a well-defined steady-state behavior requires the fixed point to be stable, which we investigated by analyzing the eigenvalues of the linearized dynamics. Specifically, we have studied in detail two versions of the fast-growth model, with instantaneous transcription and with a transcriptional time scale set by the mRNA degradation rate (see Supplementary Fig.s 6 and 7 and Supplementary Note 8 and 9). In both cases, all eigenvalues have a negative real part in the physiologically relevant parameter region, showing that the steady-growth fixed points are always stable. More in general, we also show that in the absence of transcriptional delays, a generic model for ribosome translation regulation will display stable fixed points, as long as the regulatory function is monotonically decreasing in *ϕ*_R_ and increasing in *ψ*_A_ (details and demonstration in Supplementary Note 9).

### Relaxation towards new steady state shows damped oscillations

We next investigated the model behavior during a nutritional upshift. To realize such an upshift, we varied suddenly (in a stepwise fashion) the nutrient quality *ν*, which is the parameter that characterizes the external environment.

Fig. 3A-D display the resulting relaxation pattern, characterized by damped oscillations observed across all the main quantities described by our model. It is worth highlighting that the ribosomal sector size *ϕ*_R_ also exhibits these oscillations, despite being the parameter that changes at a slower pace, given the necessity to dilute the existing proteome composition for any alteration. Fig. 3E visualizes the same oscillations by plotting the progression of the ribosomal sector proteome fraction against the amino acid (charged tRNA) pool and the translation rate *ϵ*. Conversely, ppGpp levels, amino acid (charged tRNA) pool, and translation rate change coherently following a quasi-steady-state relationship (Fig. 3F). Hence, the oscillatory behavior persists even when considering the relationship among the amino acid pool, elongation rate, and ppGpp level observed during a steady state.

**FIG. 3:**
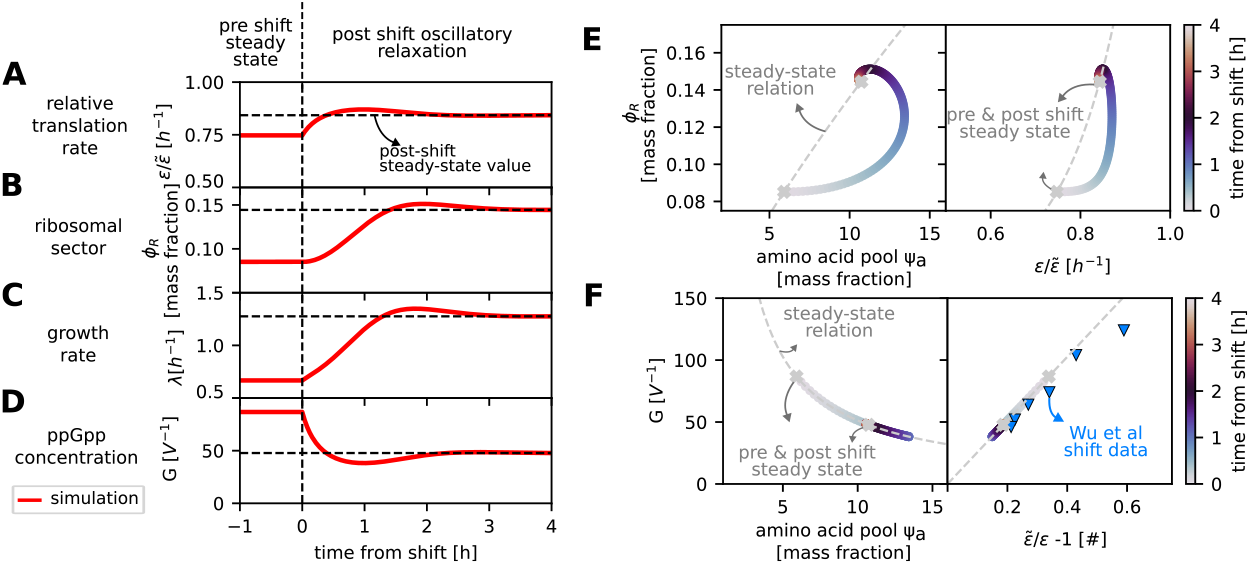
Damped oscillations characterize the relaxation dynamics towards the new steady state. The figure presents simulation predictions of the post-upshift dynamics of various state variables. Panels **A, B, C**, and **D** display the time-dependent behavior of the relative translation elongation rate 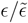, the mass fraction of the ribosomal sector *ϕ*_R_, the growth rate *λ*, and the ppGpp concentration G, respectively. Panel **E** shows the post-shift dynamics of the ribosomal sector *ϕ*_R_ in relation to the amino acids pool *ψ*_A_ and the relative translation elongation rate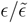, with circles representing simulation results color-coded by time from the shift. The X-shaped symbols denote the pre- and post-shift steady-state values and the dashed line represents the steady-state relationship between the two plotted variables. Similarly, panel **F** shows the post-shift dynamics of the ppGpp concentration *G* versus *ψ*_A_ and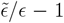, a factor which is found proportional to ppGpp [16]. In this panel, we also show shift data from [16] (blue reverse triangles). These plots show the presence of overshoots due to the damped oscillatory relaxation dynamics predicted by the model.

### Damped oscillations are independent of transcription delays

Following this observation, we asked whether the presence of oscillations stemmed from the fact that in our model, the regulatory functions *χ*_i_ are not a function of *ϵ* derived from steadystate behavior: instead, these functions emerge due to transcript dynamics. Consequently, we questioned whether the existence of oscillations was linked to the transcriptional delay *τ*_*χ*_, which establishes the timescale for adjusting the mRNA pool composition after a change in the ppGpp levels. To study this, we defined a new instantaneous-transcription model, where we set *τ*_*χ*_ = 0 (see Supplemetary Note 4 for more details). Figure 4A-B show that both the instantaneous-transcription model and the complete model (*τ*_*χ*_ *>* 0) exhibit damped oscillations during the transition. This indicates that even if the transcript pool adjusts immediately following ppGpp changes, damped oscillations persist, similar to the non-instantaneous case. Observe that in Figure 4, in order to emphasize the quantitative difference between the behavior of the two models, the value of *τ*_*χ*_ was set to 10min.

However, the important point to realize is that overshoots are expected also for *τ*_*χ*_ = 0, and since empirically *τ*_*χ*_ is small (order one minute [26]) we expect to be close to this situation.

The analysis of the eigenvalues of the two systems confirms this behavior. Indeed in both cases, the eigenvalues are complex with a negative real part in all biologically accessible growth regimes, a property linked to the oscillating relaxation to the new state. This theoretical analysis was carried out spanning all growth regimes, from slow to very fast growth, and is presented in detail in the Supplementary Note 9 and in Supplementary Fig.s 6-9. We find that the presence of a damped oscillatory response is related to the specific set of parameter values used in the model, but as these values change different regimes arise, giving rise to a typical dynamical systems “phase diagram” [34]. Our analysis, reported in Supplementary Note 9, reveals indeed three distinct regimes when the values of the parameters are changed. The first regime is characterized by a shift without oscillations (overdamped), occurring at slow growth rates. The second regime features a shift with damped oscillations, which occurs in mid-to-fast growth. Finally, the third regime is a shift with sustained oscillations. The theoretical possibility of such “oscillatory growth” has been predicted by a previous generic growth model [34]. The threshold between these regimes varies depending on the parameters of the system. Interestingly, if we call *ν*^*∗*^ the nutrient quality after which the oscillations arise, by studying the parametric dependence of *ν*^*∗*^ we find that is not just determined by the details of the resource allocation strategy, but the dynamics of the amino acid pool also plays a role (see Supplementary Note 9). Note that, when the model is defined with the parameter values found for biological systems, the overdamped and sustained oscillatory regimes disappear, and the only accessible regime remaining is the one characterized by damped oscillations.

In our model, the oscillatory behavior arises because of the effective negative feedback loop between the amino acid pool and the ribosomes. This mutual connection is sketched in panel E of Fig. 4: on one hand, ribosomes deplete amino acids due to their consumption for protein synthesis, and on the other hand the ppGpp regulatory circuit enhances ribosome levels in presence of amino acids, since higher amino acids levels lead to higher translation rates, lowering ppGpp production and therefore upregulating the ribosomal sector. On a more mathematical level, this type of feedback is a necessary condition to obtain oscillations, whether they are damped as in our case or sustained (in particular one needs a negative feedback loop with a sufficient delay [35], or a negative autoregulation, see Supplementary Note 10). Our model incudes this feedback as a double connection between amino acid pool and ribosome biogenesis, which is a crucial ingredient to make the damped oscillatory behavior possible.

**FIG. 4:**
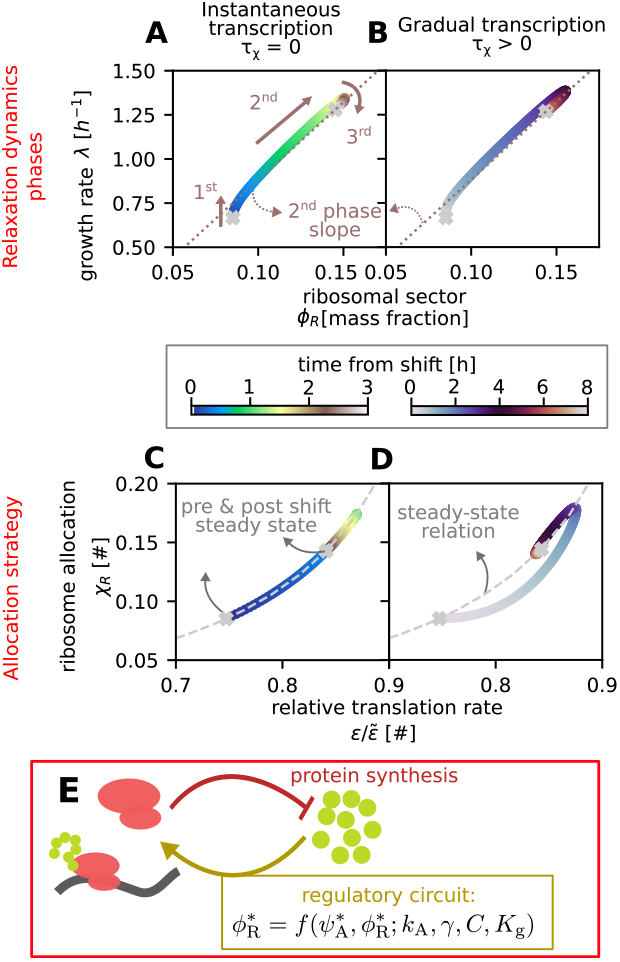
The incoherent feedback between amino acid pool and ribosome production gives rise to damped oscillatory response regardless of delays due to ribosome transcription. Simulations were performed with two different models: with instantaneous transcription (**A, C**) and with gradual mRNA and rRNA pool production (**B, D**). In the instantaneous transcription model, the regulatory functions *χ*_i_ follow ppGpp levels without delay (as in ref. [13]), while they are given by Eq. 2 in the gradual transcription model, where here we set *τ*_*χ*_ = 30min to emphasize the effect of the delay. In all panels, circles indicate simulation results color-coded by the time from the shift, and X-shaped symbols represent the pre- and post-shift steady states. Panels **A** and **B** show the three phases of post-shift adaptation, which are present in both models. The *x* axis displays the ribosomal sector mass fraction, and the *y* axis shows the growth rate. The dashed line in both panels connects (0,0) and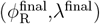, and highlights the second phase (as in Fig. 9 of ref. [13]). Panel **C** and **D** show the ribosome allocation through the shift. The *x* axis displays the relative translation rate, and the *y* axis shows the ribosomal regulatory function *χ*_R_. The plots show that oscillations occur even when the regulatory function instantaneously follows the translation elongation rate along the steadystate relation. Panel **E** provides a sketch of the incoherent feedback loop between the amino acid pool *ψ*_A_ and the ribosomal fraction *ϕ*_R_, which is explicitly described in this study and is responsible for the observed oscillatory behavior.

To interpret the different relaxation time scales involved in a nutrient shift within our model,Fig. 4A-B shows that the process of reaching a new steady state after a change in nutrient quality can be divided into three phases. The first phase is defined by a sudden increase in the translation elongation rate due to the increase in the amino acid pool size *ψ*_A_. This change in the growth rate is detected by the sensory mechanism of the ppGpp, but the synthesis of mRNA and new proteins has not yet changed, therefore, the growth increase of the first phase is driven purely by the change in the translation elongation rate *ϵ*. In the second phase, the synthesis of new proteins starts to adjust to the new protein allocation strategy given by *χ*_R_(*G*). This second phase is slower than the first one because the synthesis of new proteins is not immediate. During this phase, the translation elongation rate remains almost constant while the sector sizes change, and the relaxation proceeds along a straight line in the (*ϕ*_R_, *λ*) plane. In the third and last phase, all the relevant variables, including translational activity, regulatory functions, and protein sectors, oscillate while relaxing around the new steady state. Our study provides a detailed analysis of these three phases and their underlying mechanisms. The study by Erickson and coworkers that introduced the FCR model (ref. [13]) already presented some of the different relaxation phases we identified in our study. However, their model only included the first and second phases and did not account for the oscillatory dynamics observed in our study.

### A ribosome/precursors feedback enables oscillations

To better understand the behavior of our model upon nutrient shifts, we compared it with two existing models (see also the Supplementary Note 7 for further details): the original FCR model [13] and the Flux Parity Model (FPM) proposed by Chure and Cremer [25]. Both models were specifically developed to capture the dynamics of nutritional upshifts. The FCR model operates on a quasi-steady-state assumption linking translational activity *σ* directly to the proteome sector functions *χ*_i_, even during shifts. This approach simplifies the model by avoiding a mechanistic description of the ppGpp regulatory circuits. In order to allow a direct comparison with our model, we have extended the FCR model by incorporating the recently reported relationship between *σ* and ppGpp concentration *G* from ref. [16], which is described by Eq.s S43-S47. This extension enables predictions of ppGpp concentration changes during shifts, allowing direct comparison with our model. Notably, extending the FCR model requires accounting for sequestered ribosomes, as done in ref. [16] for steady-state growth and making assumptions about their behavior during shifts, a step not covered in the original study. This integration, however, does not change the fundamental assumption of the model, which is that protein synthesis is regulated by a direct sensing of the precursor fluxes through the the translational activity, and it follows them adiabatically via a quasisteady-state relation. Further details of this modified FCR model are provided in Supplementary Note 6.

Even considering the extended FCR model, a key difference between it and our framework lies in the treatment of the amino acids pool. The FCR model focuses on catabolic and biosynthesis fluxes to control shift dynamics, without explicitly describing the amino acids pool *ψ*_A_. Instead, the translational activity *σ* is defined as *J*_b_*/M*_R_, relying solely on flux-based sensing of external conditions.

In contrast, the FPM model [25] takes a different approach, using a flux-matching principle to set the biosynthesis rate, ensuring that uptake and biosynthesis fluxes are balanced. This strategy aligns qualitatively with the ppGpp regulatory circuit and includes sensing of charged tRNAs, a quantity indirectly linked to the amino acids pool, although not explicitly described by this model.

Fig. 5 shows that the predictions of our model, the extended FCR model, and the FPM model differ during a nutrient shift. Trivially, the extended FCR model cannot predict the amino tRNA by definition. We can instead compare the charged tRNA prediction by the FPM model and our model, as these are explicitly described in both models (note that in our model we assumed [tRNA^C^] *∝ ψ*_A_). More interestingly only our model predicts the oscillatory relaxation towards the new steady state, because among these three models it is the only one that explicitly describes the feedback relation between amino acids pool and ribosomal allocation sketched in Fig. 4E. To further support this point, in the Supplementary Note 5 we have also analyzed a version of our model with a different ppGpp regulatory circuit (incorporating solely the RelA production term, following refs. [9, 17, 21]) also shows damped oscillations (see Supplementary Note 5 and Supplementary Fig. 4).

**FIG. 5:**
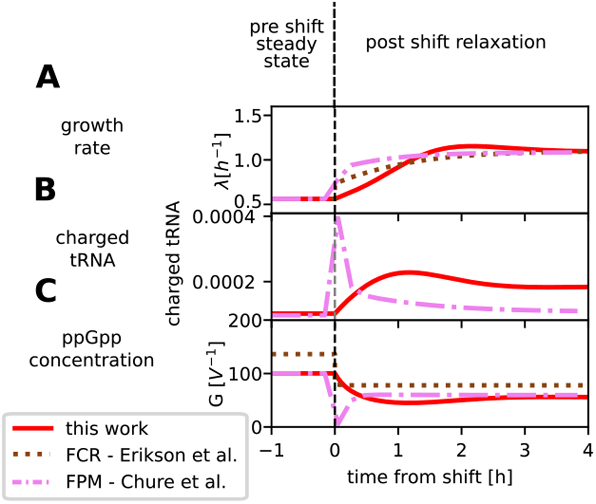
Different theoretical frameworks lead to different predictions for the relaxation dynamics. Panels A-B-C compare simulations of three different models for nutrient shift dynamics: this work (red solid line), the FCR model (ref. [13] - brown dashed line), and the Flux-Parity Model (FPM) (ref. [25] - pink dash-dotted line). Panel **A** shows the dynamics of the instantaneous growth rate. Our model is the only one that predicts an overshoot of the growth rate. Panel **B** shows the size of the amount of charged tRNAs, which is not predicted by the FCR model. For the prediction of our model, given that [tRNA^*c*^] *∝ ψ*_A_ we have normalized *ψ*_A_ to have the same pre-shift value as the FPM prediction. Panel **C** shows the ppGpp concentration *G* across the shift. This can be predicted with all three models, combining the FCR framework with Eq. 6, from ref. [16] (see Supplementary Note 6 for the upgraded FCR model definition). The discrepancies between the steady-state predictions are due to the different values of the parameters used. The figure shows that of all the models tested, just our framework predicts the oscillatory response to the nutrient shift, and this is due to the presence of the incoherent feedback between amino acids levels and the ribosomal sector.

### Damped oscillations are visible experimentally

So far, we have shown that the oscillatory relaxation behavior arises naturally from a theoretical framework that incorporates biological knowledge on nutrient sensing and regulatory response. It is natural to ask whether this predicted behavior has an experimental counterpart.

Considering published data, we observed clear oscillations in data from our reanalysis of nutritional upshift experiments performed in a microfluidic device, as reported in ref. [23] (Fig. 6AB). Our reanalysis involved examining the original data to identify patterns of overshoots and damped oscillations in growth rate and ribosome allocation during nutrient upshifts.

**FIG. 6:**
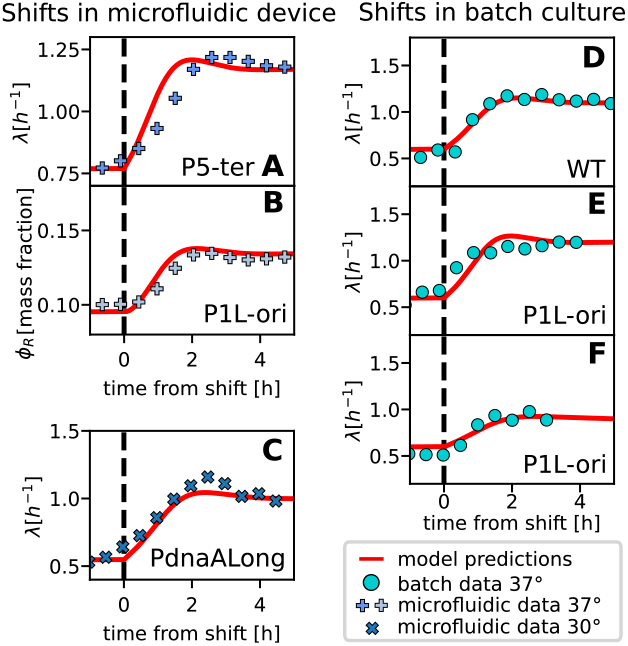
Overshoots compatible with the predictions of our model are observed in in nutrient-shift experiments. The figure shows relaxation to the new steady state for various experiments, both for the growth rate and the ribosomal sector size. In all panels, continuous red lines represent the results of a simulation of the model. **A-B:** Growth rate *λ* and ribosomal sector size *ϕ*_R_ dynamics following an upshift in a microfluidic device at 37^*°*^C, (reanalyzed data from ref. [23]). The points are sliding averages with a window of 75min (see Methods and/or Supplementary Note 11 for details on the definition of *ϕ*_R_). **C:** Growth rate *λ* dynamics following an upshift across the same media as in ref. [23] in a microfluidic mother-machine device at 30^*°*^C. The points are binned averages with a window of 40min. **D-E-F:** Growth rate *λ* dynamics following an upshift across the same media as in ref. [23] in a batch culture at 37^*°*^C, for the wild type (D) and a reporter strain in ref. [23] (E-F). All the panels refer to a biological replicate and show averages among technical replicates. The three panels refer to different experiments and summarize the three typically observed outcomes: clear overshoots, found in 8 biological replicates, shown in Supplementary Fig. 10 (D), lack of visible overshoots at odds with the model, found in 9 replicates shown in Supplementary Fig. 11 (E), and lack of observed overshoots in agreement with the model, found in 4 biological replicates shown in Supplementary Fig. 12 (F).

To test the robustness of this result, and given the lack of coherence on this point looking at other shifts performed in the literature, we also performed new shift experiments in different settings (batch and microfluidics) to investigate the existence of overshoots and damped oscillations during nutrient upshifts.

The new microfluidics experiments were conducted in a similar microfluidic “mother machine” device as in ref. [23] but at a different temperature and with strains that did not carry the ribosomal/constitutive fluorescent reporters. This allowed us to monitor the behavior of the growth rate and ensure that these high-expression GFP reporters were not the cause of the observed damped oscillations. These experiments also tested the robustness of the behavior in a shift involving slower growth conditions. Figure 6C and Supplementary Fig. 17 show the results of these new experiments, for different *ϵ. coli* strains as explained in the methods. In some cases, the observed overshoot is even larger than the one predicted by the model.

For both the reanalyzed and the new microfluidics data, overshoots (and damped oscillations) compatible with the model predictions are apparent in both the growth rate *λ* and the proxied ribosomal sector *ϕ*_R_ (where present). Note that to obtain data on the ribosomal sector, the authors of ref. [23] have used the signal of a GFP expressed from a ribosomal RNA promoter. Before using this signal as a proxy for *ϕ*_R_, we conducted several tests on the signal to see if it was compatible with the sector dynamics behavior (see Supplementary Note 11 and Supplementary Fig. 15 for details).

Additionally, the damped oscillatory behavior observed in mother machine devices is consistent across biological replicates and strains (Fig. 6DE and Supplementary Fig. 14A-B). We also note that a clear growth-rate overshoot on a minute time scale is observed consistently in the microfluidic device experiments (Supplementary Fig. 16). A similar fast overshoot was also previously reported in similar single-cell shifts providing amino acids [36], and connected by direct observation of cell volume and mass to a dilution of the cell. Hence, we surmise that cells entering an upshift effectively experience a hypo-osmotic shock, regardless of whether the experiment was carried out in a microfluidic setting or in batch. Significant biological processes may occur within this short time scale, and consequently, there could be additional physiological adaptations not captured by our model.

The results from experiments in batch culture are more ambiguous, as we found clearly visible overshoots in eight (roughly half) of the experimental replicates (Fig. 6D). Other biological replicates (nine in total) did not show overshoots where the model predicted they would be visible (Fig. 6E) or presented a very small effect in agreement with the model (four bological replicates in total, Fig. 6F). All our batch experimental results are presented in Supplementary Fig. 10, 11 and 12. To understand why model predictions vary across replicates, we need to account for the fact that while the pre-shift and post-shift state were selected to be consistent with steady growth, as the growth rates varied noticeably across replicates, and the model predicts different behavior depending on the growth rate jump.

Despite of the variability across replciates, the overshoots that are visible in these experiments occur consistently across replicates around two hours from the shift, as predicted by the model. Additionally, the batch experiments were also performed with wild-type strains, confirming the idea that growth-rate overshoots upon nutrient upshifts are not due to the presence of fluorescent reporters. Finally, the quantitatively small overshoots predicted by the model and the inferior quality of the batch data also explain why the phenomenon was not widely reported by previous literature. Besides the inferior resolution of these experiments, a possible interpretation of the difficulties encountered in batch-culture experiments could be that in batch population growth can be driven primarily by fast-growing cells, whereas in a mother-machine microfluidic device, a fixed distribution of mother cells is maintained.

To conclude, while we have provided new evidence supporting our model’s predictions, given the limitations of the presented data, our main claims remain limited to the statement that a realistic architecture (based on our current knowledge) linking nutrient sensing to ribosome allocation would generally give rise to oscillations. Biologically, it is also possible that unaccounted elements in the architecture, which are not described by our model, could alter the prediction. Further systematic experiments may clarify the situation.

## III. DISCUSSION

This work presents a dynamical modeling framework that describes the out-of-steady-state growth of *E. coli* cells through a global resource allocation dynamics. The model combines the framework of ref. [13] with a mechanistic description of nutrient sensing and global gene regulation by the ppGpp circuit. This combination of ingredients leads to predict an oscillatory relaxation after a shift in the nutrient conditions, for all shifts that lead to a moderate-to-fast growth rate. While other models predict oscillatory behavior in this context [21], recent work on growth laws in this area did not incorporate this aspect [13, 16]. Crucially, we have shown that removing any transcriptional delay in our model does not ablate the incoherent feedback loop, nor the oscillations. Hence, we conclude that the transcriptional waiting time *per se* should not be regarded as a cause for the oscillations. Our framework indeed connects the presence of oscillations to the feedback interplay between global flux balance and resource allocation and the mechanistic circuits that implement growth control. In other words, we show that the oscillatory behavior is caused by the incoherent feedback between the size of the amino acid pool and the expression of the ribosomal sector.

Importantly, we have shown that damped oscillations of growth rate, ribosome allocation and ribosome proteome fraction have been observed across studies of experimental nutrient shifts in *E. coli*. Interestingly, similar oscillations in response to perturbations have also been observed in yeast [37], where the mechanistic architecture of nutrient sensing and growth regulation is very different [2, 38–40]. This suggests that the global feedback described by our model may be a general feature, or strategy, for growth regulation based on nutrient sensing. However, there are still many unanswered questions regarding these experiments and their relationship with our model. While the mechanism provided by our model provides a possible explanation, the oscillations observed experimentally may be also influenced by other molecular players that are not considered in our study, for example the transient mismatch between volume and mass growth observed after a nutrient shift [36].

Our model builds upon and integrates previous findings in the literature [5, 10, 13, 16], combining established mechanisms into a unified framework. By analyzing these elements collectively, our approach not only consolidates existing knowledge but also generates novel predictions, crucially the intriguing emergence of damped oscillatory behavior—an aspect that has not been explicitly addressed in prior models.

We recognize that several other models have tackled related questions from various perspectives (e.g., [9, 13, 21, 25, 41–43]). Our model is intended to be complementary, emphasizing the interplay between regulatory feedback and resource allocation dynamics to uncover new insights. Specifically, our framework builds on the model proposed in ref. [13], while introducing key innovations. Most notably, we relax the steady-state assumption in defining regulatory functions, allowing for new nontrivial dynamic behavior. Additionally, our model incorporates the latest findings on the dependency of translation rates on ppGpp levels, as reported in ref. [16]. This aspect was not included in previous models that examined the ppGpp regulatory circuit [9, 21, 25].

While our primary focus is the cellular response to nutrient shifts, other studies provide complementary insights into bacterial growth physiology. For example, ref. [42] explores rapid growth rate changes at the onset of nutrient shifts, while ref. [43] investigates limiting factors of bacterial growth. Together, these works enhance our broader understanding of bacterial adaptation and growth dynamics.

Although it is now well-established that ppGpp effectively senses and responds to the translational elongation speed [16], the molecular mechanisms underlying its synthesis and degradation remain largely enigmatic—particularly regarding the role of SpoT. Once thought to function solely as a ppGpp-degrading enzyme, this simplistic view is deemed inconsistent with recent data, which suggests a more complex role for SpoT in stress response and nutrient adaptation [16], further work is still needed to understand its precise role and regulatory fully mechanisms in ppGpp metabolism. We also note that the mechanistic part of our model relies on a series of measurements on ppGpp conducted in *E. coli*, and its predictions may not apply to other bacteria.

In our model, the oscillations arise because the system works as a thermostat, with a built-in feedback system involving a sensor and an effector. The origin of oscillations in our system could be attributed to architectural constraints, evolution, or a combination of both. In order to clarify this point, it will be crucial to study this system from a control-theory perspective, and to discover the optimization goals that it follows over evolution. A previous study [41] taking this approach, has concluded that oscillatory behavior may be due to optimal control towards the adjustment of ribosome synthesis in a switch-like manner. The authors demonstrate that a precursor-only control strategy, which alternates investment between gene expression and metabolism producing an oscillatory time profile of precursor concentration performs much better than a nutrient-only strategy in a dynamical upshift scenario, by avoiding the inefficient transient accumulation of precursors.

A growth regulation system can be seen as a decision-making process that detects the environment. As nutrients in a natural environment tend to fluctuate over different time scales, we can expect that the nutrient quality can vary by fluctuations and net trends. Our model shows that sensing as a low-pass filter in response to time-varying input, preventing the system from reacting to environmental nutrient quality changes above a certain frequency, but also carries a characteristic oscillatory frequency that varies with the parameters. We speculate that these features could facilitate growth control, allowing a cell to spare energy and resources by avoiding reactions to changes that are too short-lived, but also “resonate” with specific frequencies. In our framework, the model variant without delay represents the fastest-reacting version of the network. Biologically, this fast-reacting architecture may be embodied by ribosome sequestration [44], at the cost of the extra proteome sector occupied by inactive ribosomes [32, 33]. We speculate that by studying the long-term fitness trade-offs of a system that can bypass the transcriptional delay and comparing it to one that does not one could address the question of whether *E. coli* hedges its bets on both architectures through regulated ribosome sequestration, in order to set its threshold frequency in a plastic fashion, and optimize its fitness flexibly.

## IV. METHODS

### Model description

We report here the equations needed for the definition and simulation of our mechanistic model. In these notes, we will distinguish the state variables from the parameters by expliciting the time dependence of the first ones. We start with the differential equations that define the dynamical system:

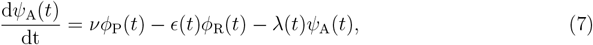

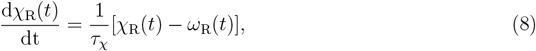

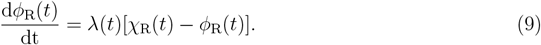

Eq. 7 describes the dynamics of the amino-acid pool, given by the balance of the upcoming nutrient flux (*νϕ*_P_), the outgoing biosynthesis flux (*ϵϕ*_R_), and a dilution term given by volume growth (*λψ*_A_). Eq. 8 is derived from the transcript dynamics, which is explained in detail in the Supplementary Note 2 and 2. Eq. 9 is derived from the sector definition as in ref. [13].

Next, we show the other definitions needed to close the system. We need to define the translation elongation rate, which depends on the amount of charged tRNAs, and therefore on the amino acids level. Eq. 4 connects these quantities:

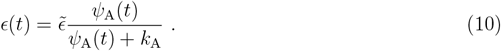

The ppGpp concentration is given by the following empirical relation from ref. [16]

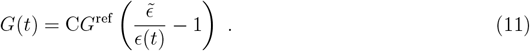

The equation presented in ref. [16] gives the fold change of the ppGpp with respect to a reference condition, therefore, in order to obtain the ppGpp concentration we add the parameter *G*^ref^ to the original equation for the fold change. The value for *G*^ref^ is 55.73 *µM* and is given by ref. [17], see Supplementary Note 1 and 2.

The RNAP allocation on ribosomal genes was assumed to follow the following relationship

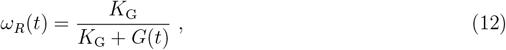

where *K*_G_ is given from a fit of the data presented in ref.s [16, 17], and its value is 8.07 *µM*. The growth rate in our model corresponds to the biosynthesis rate:

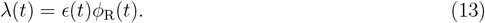

Lastly, the constitutive sector is defined as the remaining part of the proteome, which does not belong to the ribosomal nor to the housekeeping one,

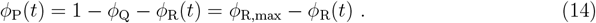

### Nutrient-shift experiments

#### Microfluidics experiments at 30^°^C

##### Strain and growth media

The strain used in this experiment is the wild-type *E. coli* strain BW25113, the parent strain of the Keio collection [45], in which promoter-reporter constructs were inserted in the chromosome as described in ref. [23, 46, 47]. The specific strains used were the ones containing the promoters P5, P5-ter [23, 46] and PdnaALong [46, 47] In upshift experiments, we used two growth media based on the M9 minimal medium as the base and glucose as the carbon source, 0.4% glucose for the slow-medium, while thee fast-medium, in addition, has 0.4% casamino acids. Bacteria were grown overnight in the slow-growth media at 30^*°*^C. Overnight cultures were diluted 500:1 in new growth medium and returned to the incubator for 3 to 4 hours. This is important to guarantee bacteria to be in the exponential phase when injected into the microfluidic device [46].

##### Mother machine experiments

We conducted experiments using a microfluidic “mother machine” device, which consists of 1-*µ*m-wide channels positioned between two larger feeding channels [48]. Bacteria were confined within the microfluidic channels by a narrow opening on one side.

The microfluidic chips were prepared following ref. [46]. In brief, the polydimethylsiloxane (PDMS) devices were fabricated from a mold using standard procedures and bonded to a microscope slide via plasma treatment. Prior to bacterial loading, each chip was passivated by incubating it with 150 *µ*l of a 2% bovine serum albumin (BSA) solution at 30*°*C for 1 hour to minimize bacterial adhesion to the glass or PDMS surfaces. After passivation, the chips were rinsed with freshly filtered medium, and approximately 1 ml of bacterial culture was manually injected into the device. Flow control within the microfluidic setup was achieved using flow sensors integrated into each feeding channel. The Elveflow pressure-driven flow system was used to ensure a continuous and stable flow of fresh growth medium through the microfluidic device at a constant speed. The entire microfluidic setup comprises a pressure-driven flow controller (0 – 2 bar pressure range), two rotary valves (11-port/10-way Mux Distributor), and two flow sensors. The two rotary valves were used to quickly change between growth medium in both the top and bottom channels. In upshift experiments, the valves were programmed to alternate so that the slow-growth medium fed the device first, while the fast-growth medium was delayed. In most cases, a complete upshift experiment lasted between 12 and 20h, with roughly equal time spent in each growth medium. These sensors provided real-time feedback to maintain precise flow rates, ensuring stable and responsive air pressure-driven flow. The system allowed for robust and long-term microfluidic experiments. Temperature regulation was maintained at 30*°*C using a custom-built temperature control system.

##### Image acquisition and data analysis

Imaging was performed using a Nikon Inverted Microscope ECLIPSE Ti-E equipped with a 100X oil immersion objective lens (numerical aperture 1.4) and a Nikon Perfect Focus System to correct for focus drift. An xy motion plate was employed to cycle through predefined regions of interest at specified time intervals. Images were captured using a 16-bit camera at a resolution of 512 x 512 pixels, with each pixel corresponding to 0.1067 *µ*m. The motorized stage and camera were programmed to image up to 40 fields of view, each encompassing approximately eight microchannels, at 3-minute intervals.

This pipeline is the same presented in ref. [46].

#### Microfluidics experiments at 37°C

Original data for this experiment was published in ref [23], please refer to this reference for the details of the experiments. Supplementary figure 14 shows the reanalized data. All the information on the additional analysis of the data are reported in the Supplementary Note 11 and Supplementary Fig. 14-16.

#### Batch experiments at 37 ^°^

##### Strains and growth media

The *E. coli* K12-derived strain BW25113 was used in all experiments, together with the “P1 Long” mutant [23, 49], with an incorporated reporter cassette in the chromosome, containing a Kanamycin resistance gene and the rRNA operon promoter *rrnBP1* followed by a GFP expressing gene (the same as in ref. [23]). Cells were grown in a M9 minimal growth medium complemented with 1% glucose (glu), until the shift to 1% glucose and 1% Casamino Acids (glu+cAA).

##### Culture growth protocol

The strains were first plated from a −80°C glycerol stock to LB-

Agar plates. Then, a preculture was made, where a single colony was inoculated in LB medium. After reached an OD of 0.3, the cells were washed of the LB medium by centrifugation (3 minutes at 8000 g-force) and resuspended in the medium used for the growth experiment (M9+glu). The overnight was prepared by diluting the cells of the preculture such that growth was still in the exponential phase at the start of the experiment.

The growth experiment was performed in the ChiBio chemostat [50], where each culture was placed in a separate reactor and kept in the same conditions at 37^*°*^C. Each reactor is composed of a glass tube filled with 20 *mL* medium and the instruments to measure OD. All other ChiBio parameters (temperature, stirring, gain intensity, etc.) were kept to the default settings presented in ref [50].

##### Growth shift protocol

Growth shifts were performed by adding a 20% solution of Casamino acids to the reactor, such that the medium had 1% cAA concentration. To retain constant growth conditions and to keep OD out of the saturation range (*OD >* 1), periodic dilutions were realized to keep the OD between 0.4 and 0.8. While the ChiBio [50] provides pumps to regulate OD, the pumping rate was not fast enough for the significant volume necessary for the dilution. Instead, a manual dilution was chosen, using a 10 *mL* pipette to dilute the reactor when OD was close to 0.8. One dilution was made before the shift, and another at the moment of the shift. After the shift, dilutions were continued for at least 2 hours of growth.

##### Data Analysis

The ChiBio outputs data files containing the precise OD every minute and the associated time stamps. To synchronize this data with the shift dynamics, the shift time was set as *t* = 0. OD outliers (*e*.*g*. measurements during dilutions) were removed, as well as any data where OD was close to saturation (*OD >* 0.8). To mitigate measurement noise from the OD, for every OD measurement *OD*_*i*_ at time *t*_*i*_, we apply a sliding window average on the logarithm of the OD (to average out deviations from exponential growth). Considering a window size *w*_1_ = 4min, we have

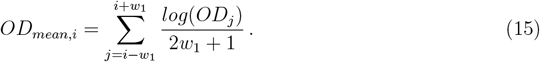

To have a continuous growth curve from start to finish of the experiment, we then consider a new OD (*OD*_*new*_) as the OD with dilutions (*OD*_*mean*_) multiplied by the dilution ratio such that there is no interruption in the growth curve due to dilutions. To measure the actual dilution ratio at every dilution, we calculate the mean growth rate *λ* at the dilution by fitting an exponential growth function on a 20 minute time interval before the dilution. As such, we have

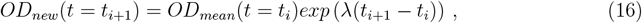

with *t*_*i*_ the last time point before dilution and *t*_*i*+1_ the first time point after the dilution. The dilution ratio is then *OD*_*new*_(*t* = *t*_*i*+1_)*/OD*_*mean*_(*t* = *t*_*i*+1_). From the continuous growth curve, we can measure the instant growth rate *λ*_*i*_ for each time *t*_*i*_. We set

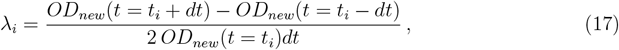

with *dt* = 3 *min*. Further binning of the growth rate aws applied for comparison of different experimental replicates. To obtain the average growth rate shown in Fig. 6 and Supplementary Fig.s 10-12 we compared different technical replicates of the same experiment, excluding realizations according to two criteria: (I) to average only technical replicates that were consistent with each other, we excluded the replicates for which the aligned OD (i.e. the OD curve normalized such that OD(t=0)=0.5) has a normalized L1 distance from the other replicates exceeding 1.5, and (II) to filter for steady-growing populations we excluded all the replicates that did not reach the shift with a steady growth rate, quantifying the steadiness of the growth rate by calculating the coefficient of variation of the growth rate 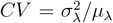in the first hour prior to the shift, where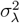is the variance and *µ*_*λ*_ is the average of the measured growth rate across time frames before the shift. If the *CV* of a replicate exceeded 0.05 the replicate was discarded. The curve for the average growth rate across replicates was then calculated for all the experiments for which at least two technical replicates had passed the screening. The results are shown in Supplementary Fig. 10-12, and an example of a technical triplet with a discarded technical replicate is shown in Supplementary Fig. 13.

## DATA AVAILABILITY

The data generated in this study have been deposited in a Mendeley repository at DOI: 10.17632/w294vd3pgh.2, ref [51].

## CODE AVAILABILITY

The code generated in this study have been deposited in a Mendeley repository at DOI: 10.17632/w294vd3pgh.2, ref [51].

## ACKNOWLEDGMENTS

We are very grateful to all the members of the Cosentino Lagomarsino lab for the useful discussions. We would like to thank our experimental collaborators Pietro Cicuta and Aske Petersen for the feedback on the data. We are grateful to Severin J. Schink, Gregory Bokinsky, Frank Bruggeman, Ulrich Gerland, Gabriele Micali, Sander Tans, and Terence Hwa for the feedback on this work.

This work was supported by AIRC - Associazione Italiana per la Ricerca sul Cancro AIRC IG grant no. 23258 (MCL and RD). RD was supported by the AIRC fellowship Italy Pre-Doc 2022 (ID 28176). PF was supported by a grant from the French National Research Agency (REF: ANR-21-CE45-0009). JG was supported by a grant Fondo Italiano per la Scienza (FIS 00002903 CUP J53C23002290001).

## AUTHOR CONTRIBUTIONS

MCL conceived the study. RD performed all the simulations with assistance from L Calabrese. RD, JG, and MCL designed the models. RD took care of the model simulations. RD handled the data analysis with GT, VF, II and PF assistance. RD, PF and II performed the experiments, with L Ciandrini and BS assistance. MCL, JG, and RD wrote the paper.

## References

[1] J. Zhu and C. B. Thompson, “Metabolic regulation of cell growth and proliferation,” Nature Reviews Molecular Cell Biology, vol. 20, pp. 436–450, Apr. 2019.

[2] L. Galdieri, S. Mehrotra, S. Yu, and A. Vancura, “Transcriptional regulation in yeast during diauxic shift and stationary phase,” OMICS: A Journal of Integrative Biology, vol. 14, pp. 629–638, Dec. 2010.

[3] P. C. Marijuan, J. Navarro, and R. del Moral, “On prokaryotic intelligence: Strategies for sensing the environment,” Biosystems, vol. 99, pp. 94–103, Feb. 2010.

[4] M. Scott and T. Hwa, “Bacterial growth laws and their applications,” Current Opinion in Biotechnology, vol. 22, pp. 559–565, Aug. 2011.

[5] M. Scott, S. Klumpp, E. M. Mateescu, and T. Hwa, “Emergence of robust growth laws from optimal regulation of ribosome synthesis,” Molecular Systems Biology, vol. 10, p. 747, aug 2014.

[6] L. U. Magnusson, A. Farewell, and T. Nyström, “ppGpp: a global regulator in Escherichia coli,” Trends in Microbiology, vol. 13, pp. 236–242, may 2005.

[7] K. Potrykus, H. Murphy, N. Philippe, and M. Cashel, “ppGpp is the major source of growth rate control in E. coli,” Environmental Microbiology, vol. 13, pp. 563–575, oct 2010.

[8] B. D. Towbin, Y. Korem, A. Bren, S. Doron, R. Sorek, and U. Alon, “Optimality and sub-optimality in a bacterial growth law,” Nature Communications, vol. 8, Jan. 2017.

[9] E. Bosdriesz, D. Molenaar, B. Teusink, and F. J. Bruggeman, “How fast-growing bacteria robustly tune their ribosome concentration to approximate growth-rate maximization,” The FEBS Journal, vol. 282, pp. 2029–2044, mar 2015.

[10] M. Scott, C. W. Gunderson, E. M. Mateescu, Z. Zhang, and T. Hwa, “Interdependence of cell growth and gene expression: Origins and consequences,” Science, vol. 330, pp. 1099–1102, nov 2010.

[11] R. Hermsen, H. Okano, C. You, N. Werner, and T. Hwa, “A growth-rate composition formula for the growth of e. coli on co-utilized carbon substrates,” Molecular Systems Biology, vol. 11, p. 801, apr 2015.

[12] S. Hui, J. M. Silverman, S. S. Chen, D. W. Erickson, M. Basan, J. Wang, T. Hwa, and J. R. Williamson, “Quantitative proteomic analysis reveals a simple strategy of global resource allocation in bacteria,” Molecular Systems Biology, vol. 11, p. 784, feb 2015.

[13] D. W. Erickson, S. J. Schink, V. Patsalo, J. R. Williamson, U. Gerland, and T. Hwa, “A global resource allocation strategy governs growth transition kinetics of Escherichia coli,” Nature, vol. 551, pp. 119–123, oct 2017.

[14] M. V. Rojiani, H. Jakubowski, and E. Goldman, “Relationship between protein synthesis and concentrations of charged and uncharged tRNATrp in Escherichia coli.,” Proceedings of the National Academy of Sciences, vol. 87, pp. 1511–1515, feb 1990.

[15] E. Goldman and H. Jakubowski, “Uncharged tRNA, protein synthesis, and the bacterial stringent response,” Molecular Microbiology, vol. 4, pp. 2035–2040, ec 1990.

[16] C. Wu, R. Balakrishnan, N. Braniff, M. Mori, G. Manzanarez, Z. Zhang, and T. Hwa, “Cellular perception of growth rate and the mechanistic origin of bacterial growth law,” Proceedings of the National Academy of Sciences, vol. 119, may 2022.

[17] A. G. Marr, “Growth rate of Escherichia coli,” Microbiological Reviews, vol. 55, pp. 316–333, jun 1991.

[18] T. M. Wendrich, G. Blaha, D. N. Wilson, M. A. Marahiel, and K. H. Nierhaus, “Dissection of the mechanism for the stringent factor RelA,” Molecular Cell, vol. 10, pp. 779–788, oct 2002.

[19] M. Mori, Z. Zhang, A. Banaei-Esfahani, J. Lalanne, H. Okano, B. C. Collins, A. Schmidt, O. T. Schubert, D. Lee, G. Li, R. Aebersold, T. Hwa, and C. Ludwig, “From coarse to fine: the absolute Escherichia coli proteome under diverse growth conditions,” Molecular Systems Biology, vol. 17, May 2021.

[20] H. Bremer and P. P. Dennis, “Modulation of chemical composition and other parameters of the cell at different exponential growth rates,” EcoSal Plus, vol. 3, feb 2008.

[21] I. Shachrai, A. Zaslaver, U. Alon, and E. Dekel, “Cost of unneeded proteins in E. coli is reduced after several generations in exponential growth,” Molecular Cell, vol. 38, pp. 758–767, jun 2010.

[22] A. Bren, Y. Hart, E. Dekel, D. Koster, and U. Alon, “The last generation of bacterial growth in limiting nutrient,” BMC Systems Biology, vol. 7, mar 2013.

[23] M. Panlilio, J. Grilli, G. Tallarico, I. Iuliani, B. Sclavi, P. Cicuta, and M. C. Lagomarsino, “Threshold accumulation of a constitutive protein explains E. coli cell-division behavior in nutrient up-shifts,” Proceedings of the National Academy of Sciences, vol. 118, apr 2021.

[24] M. Mori, S. Schink, D. W. Erickson, U. Gerland, and T. Hwa, “Quantifying the benefit of a proteome reserve in fluctuating environments,” Nature Communications, vol. 8, Oct. 2017.

[25] G. Chure and J. Cremer, “An optimal regulation of fluxes dictates microbial growth in and out of steady-state,” eLife, vol. 12, mar 2023.

[26] R. Balakrishnan, M. Mori, I. Segota, Z. Zhang, R. Aebersold, C. Ludwig, and T. Hwa, “Principles of gene regulation quantitatively connect DNA to RNA and proteins in bacteria,” Science, vol. 378, Dec. 2022.

[27] M. Nomura, R. Gourse, and G. Baughman, “Regulation of the synthesis of ribosomes and ribosomal components,” Annual Review of Biochemistry, vol. 53, pp. 75–117, June 1984.

[28] J. J. Lemke, P. Sanchez-Vazquez, H. L. Burgos, G. Hedberg, W. Ross, and R. L. Gourse, “Direct regulation of Escherichia coli ribosomal protein promoters by the transcription factors ppGpp and DksA,” Proceedings of the National Academy of Sciences, vol. 108, pp. 5712–5717, mar 2011.

[29] L. V. Aseev, L. S. Koledinskaya, and I. V. Boni, “Regulation of ribosomal protein operons rplm-rpsi, rpmb-rpmg, and rplu-rpma at the transcriptional and translational levels,” Journal of Bacteriology, vol. 198, pp. 2494–2502, Sept. 2016.

[30] B. J. Paul, M. B. Berkmen, and R. L. Gourse, “Dksa potentiates direct activation of amino acid promoters by ppgpp,” Proceedings of the National Academy of Sciences, vol. 102, pp. 7823–7828, May 2005.

[31] P. Sanchez-Vazquez, C. N. Dewey, N. Kitten, W. Ross, and R. L. Gourse, “Genome-wide effects on Escherichia coli transcription from ppgpp binding to its two sites on rna polymerase,” Proceedings of the National Academy of Sciences, vol. 116, pp. 8310–8319, Apr. 2019.

[32] X. Dai, M. Zhu, M. Warren, R. Balakrishnan, V. Patsalo, H. Okano, J. R. Williamson, K. Fredrick, Y.-P. Wang, and T. Hwa, “Reduction of translating ribosomes enables Escherichia coli to maintain elongation rates during slow growth,” Nature Microbiology, vol. 2, ec 2016.

[33] L. Calabrese, J. Grilli, M. Osella, C. P. Kempes, M. C. Lagomarsino, and L. Ciandrini, “Protein degradation sets the fraction of active ribosomes at vanishing growth,” PLOS Computational Biology, vol. 18, p. e1010059, may 2022.

[34] W.-H. Lin, E. Kussell, L.-S. Young, and C. Jacobs-Wagner, “Origin of exponential growth in non-linear reaction networks,” Proceedings of the National Academy of Sciences, vol. 117, pp. 27795–27804, Oct. 2020.

[35] U. Alon, An Introduction To Systems Biology : Design Principles Of Biological Circuits. CRC Press, 2019.

[36] E. R. Oldewurtel, Y. Kitahara, and S. van Teeffelen, “Robust surface-to-mass coupling and turgor-dependent cell width determine bacterial dry-mass density,” Proceedings of the National Academy of Sciences, vol. 118, Aug. 2021.

[37] A. P. Gasch, P. T. Spellman, C. M. Kao, O. Carmel-Harel, M. B. Eisen, G. Storz, D. Botstein, and P. O. Brown, “Genomic expression programs in the response of yeast cells to environmental changes,” Molecular Biology of the Cell, vol. 11, pp. 4241–4257, ec 2000.

[38] T. Schmelzle and M. N. Hall, “TOR, a central controller of cell growth,” Cell, vol. 103, pp. 253–262, Oct. 2000.

[39] A. G. Hinnebusch, “Translational regulation of gcn4 and the general amino acid control of yeast,” Annual Review of Microbiology, vol. 59, pp. 407–450, Oct. 2005.

[40] Y. J. Joo, J.-H. Kim, U.-B. Kang, M.-H. Yu, and J. Kim, “Gcn4p-mediated transcriptional repression of ribosomal protein genes under amino-acid starvation,” The EMBO Journal, vol. 30, pp. 859–872, ec 2010.

[41] N. Giordano, F. Mairet, J.-L. Gouzé, J. Geiselmann, and H. de Jong, “Dynamical allocation of cellular resources as an optimal control problem: Novel insights into microbial growth strategies,” PLOS Computational Biology, vol. 12, p. e1004802, Mar. 2016.

[42] Y. Korem Kohanim, D. Levi, G. Jona, B. D. Towbin, A. Bren, and U. Alon, “A bacterial growth law out of steady state,” Cell Reports, vol. 23, pp. 2891–2900, June 2018.

[43] N. M. Belliveau, G. Chure, C. L. Hueschen, H. G. Garcia, J. Kondev, D. S. Fisher, J. A. Theriot, and R. Phillips, “Fundamental limits on the rate of bacterial growth and their influence on proteomic composition,” Cell Systems, vol. 12, pp. 924–944.e2, sep 2021.

[44] X. Dai and M. Zhu, “Coupling of ribosome synthesis and translational capacity with cell growth,” Trends in Biochemical Sciences, vol. 45, pp. 681–692, Aug. 2020.

[45] T. Baba, T. Ara, M. Hasegawa, Y. Takai, Y. Okumura, M. Baba, K. A. Datsenko, M. Tomita, B. L. Wanner, and H. Mori, “Construction of Escherichia coli k-12 in-frame, single-gene knockout mutants: the keio collection,” Molecular Systems Biology, vol. 2, Jan. 2006.

[46] I. Iuliani, G. Mbemba, M. C. Lagomarsino, and B. Sclavi, “Direct single-cell observation of a key Escherichia coli cell-cycle oscillator,” Science Advances, vol. 10, July 2024.

[47] C. Saggioro, A. Olliver, and B. Sclavi, “Temperature-dependence of the dnaa–dna interaction and its effect on the autoregulation of dnaa expression,” Biochemical Journal, vol. 449, pp. 333–341, Dec. 2012.

[48] Z. Long, E. Nugent, A. Javer, P. Cicuta, B. Sclavi, M. Cosentino Lagomarsino, and K. D. Dorfman, “Microfluidic chemostat for measuring single cell dynamics in bacteria,” Lab on a Chip, vol. 13, no. 5, p. 947, 2013.

[49] Q. Zhang, E. Brambilla, R. Li, H. Shi, M. Cosentino Lagomarsino, and B. Sclavi, “A Decrease in Transcription Capacity Limits Growth Rate upon Translation Inhibition,” mSystems, vol. 5, pp. 10.1128/msystems.00575-20, Sept. 2020. Publisher: American Society for Microbiology.

[50] H. Steel, R. Habgood, C. L. Kelly, and A. Papachristodoulou, “In situ characterisation and manipulation of biological systems with chi.bio,” PLOS Biology, vol. 18, pp. 1–12, 07 2020.

[51] R. Droghetti and M. Cosentino-Lagomarsino, “Code and data for “tincoherent feedback from cou-pled amino acids and ribosome pools generates damped oscillations in growing bacteria, mendeley repository, doi: 10.17632/w294vd3pgh.2,” 2024.

[52] J. Forchhammer and L. Lindahl, “Growth rate of polypeptide chains as a function of the cell growth rate in a mutant of Escherichia coli 15,” Journal of Molecular Biology, vol. 55, pp. 563–568, Feb. 1971.

[53] A. Schmidt, K. Kochanowski, S. Vedelaar, E. Ahrné, B. Volkmer, L. Callipo, K. Knoops, M. Bauer, R. Aebersold, and M. Heinemann, “The quantitative and condition-dependent Escherichia coli proteome,” Nature Biotechnology, vol. 34, pp. 104–110, jan 2016.

[54] S. H.-J. Li, Z. Li, J. O. Park, C. G. King, J. D. Rabinowitz, N. S. Wingreen, and Z. Gitai, “Escherichia coli translation strategies differ across carbon, nitrogen and phosphorus limitation conditions,” Nature Microbiology, vol. 3, pp. 939–947, July 2018.

[55] R. A. Lazzarini, M. Cashel, and J. Gallant, “On the regulation of guanosine tetraphosphate levels in stringent and relaxed strains of Escherichia coli.,” The Journal of biological chemistry, vol. 246, pp. 4381–4385, July 1971.

[56] J. Ryals, R. Little, and H. Bremer, “Control of rrna and trna syntheses in Escherichia coli by guanosine tetraphosphate,” Journal of Bacteriology, vol. 151, pp. 1261–1268, Sept. 1982.

[57] E. Sarubbi, K. E. Rudd, and M. Cashel, “Basal ppgpp level adjustment shown by new spot mutants affect steady state growth rates and rrna ribosomal promoter regulation in Escherichia coli,” Molecular and General Genetics MGG, vol. 213, pp. 214–222, Aug. 1988.

[58] V. J. Hernandez and H. Bremer, “Guanosine tetraphosphate (ppgpp) dependence of the growth rate control of rrnb p1 promoter activity in Escherichia coli.,” The Journal of biological chemistry, vol. 265, pp. 11605–11614, July 1990.

[59] M. H. Buckstein, J. He, and H. Rubin, “Characterization of nucleotide pools as a function of physiological state in Escherichia coli,” Journal of Bacteriology, vol. 190, pp. 718–726, Jan. 2008.

[60] N. C. E. Imholz, M. J. Noga, N. J. F. van den Broek, and G. Bokinsky, “Calibrating the bacterial growth rate speedometer: A re-evaluation of the relationship between basal ppgpp, growth, and rna synthesis in Escherichia coli,” Frontiers in Microbiology, vol. 11, Sept. 2020.

