## Supplementary notes for "Incoherent feedback from coupled amino acids and ribosome pools generates damped oscillations in growing E. coli"

These Supplementary Notes explain in more detail the model introduced in the main text, illustrate new versions of this framework and the differences in terms of predictions. They also include the analytical calculations of the fixed points and the eigenvalues of the system, they describe the new data gathered for this study and lastly, they describe how we made use of the data reported in ref. [1] to compare with our analytical findings.

### 1. SUPPLEMENTARY NOTE 1: BRIEF DESCRIPTION OF THE MODEL.

To ensure clarity, we begin by providing a synthetic overview of all the components of the model. The following presents a detailed description of the model equations. Our framework starts from the existing "flux-control regulation" (FCR) model [2] and integrates it with an explicit description of three different regulatory modules related to ppGpp dynamics. Fig. 1 of the main text shows the four ingredients of the proposed framework, which we describe in the following paragraphs. Throughout this Supplementary Note, we distinguish the system state variables from the system parameters by making explicit the time dependence of the formers.

*a. Global allocation functions set the proteome sectors sizes.* As in the FCR model, in our model some "global allocation" functions determine the sizes of the proteome sectors. These allocation functions, denoted as $\chi_{\mathrm{i}}(t)$, set the fraction of the total biosynthesis flux that is redirected toward a specific sector, thereby determining the dynamics of the sector sizes ($\phi_{\mathrm{i}}(t)$). Our model focuses on three sectors: ribosomal ($\phi_{\mathrm{R}}(t)$), constitutive ($\phi_{\mathrm{P}}(t)$), and housekeeping ($\phi_{\mathrm{Q}}(t)$), with $\chi_{\mathrm{R}}(t)$, $\chi_{\mathrm{P}}(t)$, and $\chi_{\mathrm{Q}}(t)$ determining the number of ribosomes making ribosomes, constitutive proteins, and housekeeping proteins, respectively. The time course of the sector size is described by the same equation as in the FCR model [2]:

$$\frac{\mathrm{d}\phi_{\mathrm{i}}(t)}{\mathrm{dt}} = \lambda(t)[\chi_{\mathrm{i}}(t) - \phi_{\mathrm{i}}(t)]; \mathrm{i} \in [\mathrm{R}, \mathrm{P}, \mathrm{Q}]. \tag{S1}$$

Here, $\lambda(t)$ represents the growth rate (i.e. the biosynthesis rate), and the term $\chi_{\mathrm{i}}(t) - \phi_{\mathrm{i}}(t)$ specifies the rate of change of the sector size.

*b. ppGpp-mediated transcripts dynamics determine the proteome allocation functions.* Our model proposes that the allocation of the total biosynthesis flux is determined by the composition of the mRNA pool, which is controlled, for the ribosomal sector, by ppGpp-mediated transcript dynamics. Without post-transcriptional control [3, 4], ribosomes randomly attach to transcripts and initiate protein synthesis. Therefore, the mRNA pool composition determines the partition of the flux towards the three proteome sectors. We define $\chi_{\rm R}$ as the ratio of ribosomal to total transcripts ($\frac{T_{\rm R}}{T}$), and $\chi_{\rm P}$ and $\chi_{\rm Q}$ in a similar way. The dynamics of ribosomal transcripts ($T_{\rm R}$) is governed by the ppGpp circuit that interacts with RNA polymerase (RNAP) and this dependence is encoded in the regulatory function

$$\omega_{\rm R}(G(t)) = \frac{K_{\rm G}}{K_{\rm G} + G(t)} \ , \tag{S2}$$

where $\omega_{\rm R}$ represents the fraction of RNAP that transcribes ribosomal genes. The function for $\omega_{\rm R}$ is a fit of experimental data presented in ref.s [5, 6] (Supplementary Fig. 1). The dynamics of ribosomal transcripts is given by

$$\frac{{\rm d}T_{\rm R}(t)}{{\rm dt}} = \beta\omega_{\rm R}(G(t))P - \delta T_{\rm R}(t) \ , \tag{S3}$$

where $\beta$ and $\delta$ are the mRNA production and degradation rates (representing the average behavior of many genes), respectively, and $P$ is the number of available RNAPs. Analogously, total transcripts follow the equation

$$\frac{{\rm d}T(t)}{{\rm dt}} = \beta P - \delta T(t) \ , \tag{S4}$$

The differential equation that governs $\chi_{\rm R}$ derives from its definition and Eqs. (S3) and (S4)

$$\frac{{\rm d}\chi_{\rm R}(t)}{{\rm dt}} = \frac{1}{\tau_\chi}[\omega_{\rm R}(G(t)) - \chi_{\rm R}(t)] \ , \tag{S5}$$

where $\tau_\chi \simeq 1/\delta$ is the time scale separating the (fast) adjustment of the ppGpp level to the new condition and the consequent adjustment of the transcript pool (on the order of minutes, we typically set $\tau_\chi = 1$min) [4].

*c. Translation elongation speed determines ppGpp production and degradation.* A recent study by Wu et al. [6] found a clear relationship between ppGpp level and translation elongation rate ($\epsilon$),

$$G(t) \propto \frac{\tilde{\epsilon}}{\epsilon(t)} - 1, \tag{S6}$$

where $\tilde{\epsilon}$ is the theoretical maximum translation elongation rate. This relationship can be explained by a model where the rate of ppGpp synthesis depends on the time ribosomes spend elongating ($\tau_{\rm trans}$), while the rate of degradation is proportional to the time spent waiting for a charged tRNA ($\tau_{\rm dwell}$) [6]. This finding invalidates previous models [5, 7, 8] that only described ppGpp synthesis by RelA based on loading of uncharged tRNAs, and suggests ppGpp degradation by SpoT plays a more complex role than previously assumed.

Because of the rapid production and degradation rates of ppGpp (on the order of seconds) [7], we assume that ppGpp concentration is always at a quasi-steady state. Moreover, the data from

Wu et al. [6] showed that this relationship between $\epsilon$ and $G$ holds even during out-of-steady-state scenarios. The translation elongation speed $\epsilon$ is related to the fraction of charged tRNA, thus the last module that closes the circle is the dynamics of the free amino acids denoted as $\psi_{\mathrm{A}}$.

*d. Nutrient influx, biosynthesis outflux, and growth govern the amino acid levels and the translation rate.* The mass of amino acids $A$ in the cell is determined by the equilibrium between nutrient influx and outflux due to biosynthesis [9]

$$\frac{\mathrm{d}A(t)}{\mathrm{dt}} = \nu M_{\mathrm{P}}(t) - \epsilon(t) M_{\mathrm{R}}(t) \ , \tag{S7}$$

where $\nu$ is a “nutrient quality” parameter representing a mass-specific catabolic flux [2, 10], and $M_{\mathrm{P}}$ and $M_{\mathrm{R}}$ are the masses of the constitutive and ribosomal proteins, respectively. To obtain an equation with the proteome sectors, we define $\psi_{\mathrm{A}} = A/M_{\mathrm{tot}}$, where $M_{\mathrm{tot}}$ is the mass of total proteins. If we assume a constant dry-mass density [2, 11], we obtain the following equation for $\psi_{\mathrm{A}}$,

$$\frac{\mathrm{d}\psi_{\mathrm{A}}(t)}{\mathrm{dt}} = \nu \phi_{\mathrm{P}}(t) - \epsilon(t)\phi_{\mathrm{R}}(t) - \lambda(t)\psi_{\mathrm{A}}(t) \ , \tag{S8}$$

where the additional term $\lambda\psi_{\mathrm{A}}$ accounts for dilution due to cellular growth. Here, $\phi_{\mathrm{P}}$ and $\phi_{\mathrm{R}}$ denote the fractions of the proteome allocated to the constitutive and ribosomal proteins, respectively. The influx of nutrients and the outflux of amino acids due to biosynthesis are determined by the proteome allocation, which in turn is governed by the dynamics of the ppGpp-mediated transcriptome, as described earlier. The translation elongation rate, $\epsilon$, is also influenced by the amino-acid levels via the availability of charged tRNAs, as discussed earlier.

### 2. SUPPLEMENTARY NOTE 2: MODEL DESCRIPTION - IN DEPTH.

This section presents all the equations required for defining and simulating our model. These equations are also presented in the main text; here we provide a comprehensive description and discuss their validity ranges.

#### A. Dynamics of amino acid pools are set by catabolic flux, protein production and dilution due to volume growth

We start with the equation that describes the dynamics of the amino acids pool valid outside of balanced-growth conditions,

$$\frac{\mathrm{d}\psi_{\mathrm{A}}(t)}{\mathrm{dt}} = \nu \phi_{\mathrm{P}}(t) - \epsilon(t)\phi_{\mathrm{R}}(t) - \lambda(t)\psi_{\mathrm{A}}(t), \tag{S9}$$

where $\psi_A = A/M_{tot}$ is the rescaled amino acids mass as defined in the main text. Although there are twenty different amino acid species, for simplicity we have considered one quantity to describe the amino acid pool. We interpret this as describing the limiting species. The terms on the right-hand side of Eq. (S9) above represent (I) the inflow of amino acids resulting from the catabolic flux $J_A(t) = \nu\phi_P(t)$, (II) the biosynthesis flux $J_b(t) = \epsilon(t)\phi_R(t)$, and (III) a dilution term due to volume growth $-\lambda(t)\psi_A(t)$. The inflow of amino acids is a result of various processes such as nutrient uptake, the conversion of nutrients into central precursors, and the synthesis of specific amino acids. The nature of these processes varies based on the specific nutrients provided to the cells, for instance, whether they consist of solely carbon sources or a nutrient-rich medium containing amino acids. In our model, we have chosen to provide a high-level description of these processes, combining them into a single flux. The nutrient uptake and amino-acid flux is mediated by proteins that do not belong to the ribosomal sector $\phi_R$ [10] nor to the housekeeping sector $\phi_Q$, justifying the assumption that the flux depends on the constitutive/catabolic sector $\phi_P$.

Previous studies have examined the growth-dependent expression of these proteins, particularly those involved in carbon source uptake [12, 13]. This regulation has been incorporated into the FCR framework describing shifts between different carbon sources [2]. While our model derives from the FCR framework, we have chosen for simplicity a less detailed and more generic version of this regulation. As uptake proteome sectors are part of the unregulated constitutive sector $\phi_P$, our model includes only this information. While our model may sacrifice some details of the regulation described in references [12, 13], it is not limited to describing carbon sources alone. Consequently, it avoids certain constraints, such as the maximum growth rate for growth on carbon-only nutrients $\lambda_C = 1.17h^{-1}$ set in ref. [2]. Additionally, to our knowledge, there is no quantitative analysis of substrate uptake in rich media containing amino acids. In any case, this modeling choice is not restrictive, and it is possible to adopt a more detailed description without affecting our main results.

The other main factor setting the amino-acid flux is the nutrient quality $\nu$, which represents the sum of the nutrient qualities of every individual substrate present in the growth medium. Once again, for simplicity, we chose not to describe a specific uptake sector for each substrate component in our model as in ref. [2], but this extension is simple to explore. We can still group all nutrient qualities into a single global parameter interpreting it as follows. Each nutrient quality $\nu_i$ can be expressed as $\nu_i = \nu_i' f_i$, where $\nu_i'$ represents the uptake rate per protein mass [2], and $f_i$ is the fraction of the P sector occupied by the uptake proteins of the $i$-th substrate, such that $\phi_{\mathrm{P}} f_{\mathrm{i}} = \phi_{\mathrm{up,i}}$. Using these definitions, we can express the uptake flux as follows,

$$J_{\mathrm{A}}(t) = \sum_{\mathrm{i}} \nu_{\mathrm{i}} \phi_{\mathrm{i}}(t) = \sum_{\mathrm{i}} \nu_{\mathrm{i}}' \phi_{\mathrm{up,i}}(t) \ . \tag{S10}$$

It should be noted that this simplified description of the catabolic flux has certain implications. With this approach, the only relevant change for a nutrient shift is a change in the value of $\nu$. For instance, these restrictions do not make it possible to describe shifts between carbon sources with very similar nutrient qualities, such as from glucose to lactose. Additionally, when we model an upshift by a stepwise increase in $\nu$, and the catabolic flux $\nu\phi_{\mathrm{P}}$ initially changes due to the change in $\nu$, we implicitly assume that all the proteins responsible for the uptake of the new substrate are already expressed in the pre-shift condition. However, this assumption is not always valid [14]. For a more precise description of the uptake process, we refer to the FCR model [2], which is specifically designed to describe shift data between different carbon sources. Our model can be endowed with the same ingredients if necessary.

Turning to the biosynthesis flux, we choose to use a description similar to the one reported in ref [2]: $J_{\mathrm{b}}(t) = \epsilon(t)\phi_{\mathrm{R}}(t)$, but in our case $\epsilon(t)$ is the actual translation elongation rate instead of the ribosomal activity (which represents an *average* translation rate per ribosome). Note that, in general, the ribosomal activity can be different from the elongation rate, not only because the ribosome can have different speeds but also because the ribosomal activity takes into account ribosomes that can be inactive or sequestered [2, 6, 15]. Supplementary Fig. 3 shows that at fast growth the contributions of the sequestered ribosomes (and degradation) can be neglected, therefore in the fast-growth regimes, the ribosomal activity and the elongation rate are approximately the same. We chose to model the translation elongation rate instead of the ribosomal activity because in our model the former is directly connected to the amino-acid availability, as we describe in the next paragraphs. In a variant of the model applicable to slow-growth data (see below), we have considered an additional factor $f_{bg}$ representing the fraction of active ribosomes as well as protein degradation [16]. This version of the model is described in one of the subsequent sections of these notes.

Finally, the last term of Eq. (S9), the dilution term given by growth, appears in the equation because of the definition of the rescaled mass $\psi_{\mathrm{A}} = M_{\mathrm{A}}/M_{\mathrm{prot}}$. Previous studies have neglected this last term, making the assumption of perfect balancing between the metabolic and biosynthesis [9]. However, neglecting this term leads to inconsistencies in the equations for the concentration of amino acids that do not appear acceptable. Indeed, if we neglect the dilution term in Eq.S9, reduces to

$$\frac{\mathrm{d}\psi_{\mathrm{A}}}{\mathrm{dt}} = \nu\phi_{\mathrm{P}} - \epsilon\phi_{\mathrm{R}} \tag{S11}$$

Therefore the steady-state condition requires $\nu\phi_{\mathrm{P}} = \epsilon\phi_{\mathrm{R}}$. Inserting this condition into Eq. S7 contradicts the exponential accumulation of total amino acids necessary to sustain exponential growth. If we consider the dilution term, we obtain the correct equation for the concentration of amino acids, which includes the dilution term associated with cellular growth. Therefore we kept the dilution term in the definition of our model and in our numerical simulations, which also plays a role in how we describe tRNA charging and elongation [5]. Note that this is not just a mathematical requirement, as neglecting this term has an impact on the relaxation dynamics after a perturbation, which is one of the main phenomena studied in this work, as also pointed out in ref. [17]. Supplementary Fig. 2 shows that if we neglect the dilution term the system needs a much longer time to reach the new equilibrium.

#### B. The elongation rate senses the amino acids pool through charged tRNAs

It is widely recognized that the translation elongation rate is closely tied to the availability of amino acids and other precursors, quantities that play a crucial role in the bacteria's stringent response [2, 7, 15].

In our framework, we account for this precursor dependence by relating the translation elongation rate to the concentration of charged tRNAs, similar to what is presented in ref. [15]. Specifically, we define the elongation rate as follows,

$$\epsilon(t) = \tilde{\epsilon}\frac{[tRNA^{\mathrm{C}}]}{[tRNA^{\mathrm{C}}] + k_{\mathrm{C}}} \; . \tag{S12}$$

In this equation, $\tilde{\epsilon}$ represents a theoretical maximum translation elongation rate, which we estimate empirically as

$$\tilde{\epsilon} = \gamma\frac{\phi_{\mathrm{R}}(\tilde{\epsilon}) - \phi_{\mathrm{R}}^{\mathrm{min}}}{\phi_{\mathrm{R}}(\tilde{\epsilon})} \; , \tag{S13}$$

an empirical expression obtained from the condition $\lambda_{\mathrm{max}} = \tilde{\epsilon}\phi_{\mathrm{R}}(\tilde{\epsilon})$ and the first growth law $\phi_{\mathrm{R}} = \phi_{\mathrm{R}}^{\mathrm{min}} + \lambda/\gamma$. For this derivation, we use $\phi_{\mathrm{R}}(\tilde{\epsilon}) = 1$ in order to get the maximum theoretical elongation rate. This is a requirement of our framework. Indeed, by analyzing the expression of the fixed points of the system (see section 8, Eq. S60) one notices that $\epsilon$ can reach $\tilde{\epsilon}$ if and only if $\phi_{\mathrm{R}}$ reaches unity. Note that this is just a theoretical requirement, as experimentally the ribosomal sector cannot reach unity because of the presence of the other protein sectors.

More specifically, we propose a model based on the following key results from ref. [15]: (I) in steady conditions, the ratio of the concentration of tRNA and rRNA is constant across growth rates, leading to approximately 7 tRNAs per ribosome, and maintained by gene regulation (II) the ratio of elongations factors to ribosomes is constant across growth rates (there are approximately 6 EF per ribosome) and lastly, (III) the percentage of charged tRNA is approximately the 70% of all tRNAs, and it is constant across growth conditions. These results suggest that the main factor that limits translation elongation speed is the diffusion of tRNA (charged or uncharged) toward the ribosomes. Indeed, this hypothesis would explain why the elongation speed increases as the growth rate increases, even if the ratio of tRNA to ribosomes remains constant. In a diffusion-limited scenario, the translation-elongation rate is given by the overall concentration of tRNA, which increases with the growth rate, rather than the amount of tRNA per ribosome, leading to Eq. S12.

In a dynamic situation such as a nutrient shift, tRNA charging will change faster than the total amount of tRNA as the amount of available amino acids has increased. Once the cell reads this signal, it can up-regulate the transcription/translation tRNAs and ribosomes. Therefore, in order to make the above equation S12 compatible with the shift dynamics without describing explicitly tRNA regulation, we use the ansatz that the amount of charged tRNAs is proportional to the amino acids concentration: $[tRNA^{\mathrm{C}}] \propto \psi_{\mathrm{A}}$. By this ansatz obtain the relation

$$\epsilon(t) = \tilde{\epsilon}\frac{\psi_{\mathrm{A}}}{\psi_{\mathrm{A}} + k_{\mathrm{A}}} \ . \tag{S14}$$

This equation captures the fast dependency of the translation elongation rate on the availability of amino acids and provides a link between the concentration of charged tRNAs and the translation activity. The constant $k_{\mathrm{A}}$ has not been directly measured to our knowledge. Therefore, we uses a fit to set the value of $k_{\mathrm{A}}$ that that best reproduced the experimental data shown in Fig 6 of the main text. The fact that all the different experiments (in bulk and mother machine) trends can be reproduced by using a very similar value of $k_{\mathrm{A}}$ (about 2 [mass fraction]) is remarkable, although this value is different from a previous theoretical estimate of the same parameter ($5 \times 10^{-3}$ in ref. [9]).

Finally, we observe that the available data [6] lead to the following empirical relation

$$\epsilon(t) = \tilde{\epsilon}\frac{\phi_{\mathrm{R}}}{\phi_{\mathrm{R}} + \phi_{\mathrm{R}}^{0}} \ , \tag{S15}$$

suggesting a tight mirror relationship between precursor pools, charged tRNA concentration, and ribosomal sector size in a steady condition.

### C. ppGpp senses the translation elongation rate through uncharged tRNAs

In order to model the ppGpp sensing and dynamics, we used the recent results obtained Wu and coworkers [6]. In brief, the authors find that the level of ppGpp is related to the translation elongation rate $\epsilon$ via a simple empirical relation

$$G(t) \propto C\left(\frac{\tilde{\epsilon}}{\epsilon(t)} - 1\right) , \tag{S16}$$

where $C$ is a parameter fitted in ref. [6]. We write that $G$ is proportional to the right-hand term of the above equation because Wu and coauthors measure the fold change of ppGpp with respect to a reference condition, which they chose as the exponential growth on glucose. In order to get the ppGpp concentration one needs to multiply the right-hand term of Eq. S16 with the value of $G$ found in glucose, estimated to be $G^{\text{ref}} = 55.73\mu M$. This estimate is derived from the data gathered by Marr in ref. [5] (see Supplementary Note section V for the details). Differently from the study of Wu and coworkers [6], this study considers the concentration of ppGpp instead of the fold change and allows us to estimate the ppGpp concentration at various growth rates.

Importantly, Wu and coworkers [6]) demonstrate that the relationship between $G$ and $\epsilon$ holds during dynamic regimes, beyond steady states. This suggests that the rapid synthesis and degradation dynamics of ppGpp maintain its pool in a steady-state relationship with the translation elongation rate. Consequently, we consider the regime in which the level of ppGpp $G$ remains at a steady-state with $\epsilon$, and hence it is described by an algebraic relationship rather than an ordinary differential equation (ODE).

It is important to stress that $G$ depends on $\epsilon$ through the ratio $\psi_{\text{A}}/(\psi_{\text{A}}+k_{\text{A}})$. In other words, ppGpp concentration is strictly related to the amount of charged tRNAs, which determines the ribosome waiting time $\tau_{\text{dwell}}$. In the presence of an elongation-inhibiting drug, the ribosomes split into two populations, those that are stuck by the drug do not elongate and are not linked to ppGpp production or degradation, and the others feel an increased pool of charged tRNA, making the elongation rate (conditionally to where elongation is possible) higher [6].

#### D. Regulatory functions emerge from transcripts dynamics

In the FCR framework, regulatory functions are crucial in determining the target size for proteome sector production, ultimately leading to a steady-state growth condition, where $\phi_{\text{i}}(t) = \chi_{\text{i}}(t)\forall t$. Mathematically, these functions represent the fraction of the total biosynthetic flux directed towards the $i$-th sector. However, this definition does not provide insights into the microscopic processes that give rise to the emergence of $\chi_{\text{i}}$. Intuitively, this flux fraction must be set by the fraction of ribosomes that are translating proteins of a given sector.

In the FCR model [2], the authors bypass the problem by using a quasi-steady-state approximation that sets the values of $\chi_{\text{i}}$ based on $\chi_{\text{i}}(t) = \phi_{\text{i}}^*(\sigma(t))$, where $\phi_{\text{i}}^*$ represents the steady-state relationship between $\phi_{\text{i}}$ and translational activity $\sigma$. This approach ensures that even during kinetic regimes following perturbations, the regulatory functions adjust to changes in ribosomal activity in a quasi-steady-state manner (note this formalism defines the translational activity $\sigma$ as $\lambda/\phi_{\text{R}}$, rather than solely the translation elongation rate $\epsilon$ [2], hence $\sigma$ includes a factor from the fraction of active ribosomes). This assumption is justified by the fact that the allocation of ribosomes should reproduce steady-growth states. However, this top-down approach has a drawback in that it has a limited ability to provide insights into the biological mechanisms governing the regulation of these functions and their underlying definition. Our framework aims at overcoming this limitation by distinguishing translational activity from a translation rate linked to the charged tRNA pool and providing a more mechanistic description of the origin of regulatory functions.

To distribute the biosynthetic flux among different proteome sectors, ribosomes must produce proteins in accordance with the ratios determined by the regulatory functions. In our model, ribosomes randomly attach to mRNA transcripts and initiate the synthesis of the corresponding proteins. Therefore, neglecting post-transcriptional regulation [4], the distribution of the biosynthesis flux is solely determined by the composition of the transcript pool.

With this perspective, we can define the regulatory functions as the ratio of the number of transcripts coding for proteins belonging to the $i$-th sector, denoted as $T_{\mathrm{i}}(t)$, to the total number of transcripts, denoted as $T(t)$. This is represented mathematically as

$$\chi_{\mathrm{i}}(t) = \frac{T_{\mathrm{i}}(t)}{T(t)}. \tag{S17}$$

It is important to note that $\Sigma_{\mathrm{i}} T_{\mathrm{i}}(t) = T(t)$, as is the case for the proteome sectors.

To study the dynamics of $\chi_{\mathrm{i}}$, we need to consider the dynamics of transcripts. Transcripts are rapidly degraded and produced by RNA polymerases, denoted as $P$ and assumed to be constant. Thus, the dynamics of transcripts can be described by the following equation,

$$\frac{\mathrm{d}T(t)}{\mathrm{dt}} = \beta P - \delta T(t) \ , \tag{S18}$$

where $\beta$ and $\delta$ are constants representing the mRNA synthesis and degradation rates, respectively. This is analogous to the approach presented in ref. [4], where the authors break down the dynamics of the mRNA pool by studying the production and degradation flux. Eq. S18 represents the dynamics of the total number of transcripts. To study the dynamics of transcripts belonging to the i-th sector, we can replace the total RNAP $P$ with the fraction of RNAP producing transcripts for the i-th sector, denoted as $\omega_{\mathrm{i}}(t)P$. This leads to the following equation

$$\frac{\mathrm{d}T_{\mathrm{i}}(t)}{\mathrm{dt}} = \beta \omega_{\mathrm{i}}(t) P - \delta T_{\mathrm{i}}(t) \ . \tag{S19}$$

By combining Eq. S18 and S19, along with the definition of the regulatory function in Eq. S17, we obtain

$$\frac{\mathrm{d}\chi_{\mathrm{i}}(t)}{\mathrm{dt}} = \frac{1}{\tau_\chi}[\omega_{\mathrm{i}}(t) - \chi_{\mathrm{i}}(t)]. \tag{S20}$$

Here, $\tau_\chi$ is a time constant, which represents the typical time scale for adjusting the composition of the transcripts pool. We set $\tau_\chi = 1/\delta$, based on a few simplifying assumptions. This expression can be obtained by combining Eq. S18 and Eq. S19 and assuming that the total amount of transcripts $T$ is at steady state, $T = \beta P/\delta$. Hence, this expression is valid under the assumption that total transcript and total RNAP concentrations do not vary during the shift, and that transcriptional adaptation occurs faster than exponential growth. This is coherent for *E. coli* where transcript degradation is faster than the growth rate ($\delta \ll \lambda$). In our simulations, we set $\tau_\chi = 1$ minute which is a typical value for transcript half-life in *E. coli* [4]. Recent results show that available RNAP changes across growth conditions [4], and our model can be easily adapted to include this feature, by modifying $\tau_\chi$.

We chose to focus for simplicity on three sectors, namely $\phi_{\mathrm{R}}$,$\phi_{\mathrm{P}}$, and $\phi_{\mathrm{Q}}$, and the constraint $P + R + Q = 1$ applies to both $\phi$ and $\omega$, we only need to specify the ppGpp dependence for the RNAP partition function of ribosomal transcripts, $\omega_{\mathrm{R}}$.

A further aspect of the model we need to address is the allocation of RNA polymerases (RNAPs), specifically in relation to ribosomal transcripts. In *E. coli*, it is known that the stringent response, mediated by ppGpp, regulates the production of ribosomal transcripts by directly influencing the allocation of RNAPs (see ref.s[3, 18] for details). We have left implicit the role of gene dosage in this regulation. Ribosomal protein genes and ribosomal RNAs are found near the replication origin, which affects their copy number across growth conditions [19]. However the recent findings by Balakrishnan and coworkers [4] suggest that the mRNA pool composition is mainly determined by the regulation of the promoter on-rates rather than other factors such as gene dosage and mRNA degradation rates. Therefore, while it is possible to study the role of these other factors by incorporating them in model variants, we have assumed for simplicity that the functions $\omega_{\mathrm{i}}$ just depend on promoter on-rates, which in our theory are just set by ppGpp in the case of the genes belonging to the ribosomal sectors (although one could also add catabolic regulators in a straightforward way).

We describe the allocation of RNAPs to ribosomal genes by the following equation,

$$\omega_R(t) = \frac{K_{\mathrm{G}}}{K_{\mathrm{G}} + G(t)}, \tag{S21}$$

where $K_{\mathrm{G}}$ is determined by fitting the data presented in ref.s [5, 6] and replotted in Supplementary Fig. 1, with a fitted value of $8.07\mu M$.

We highlight the fact that alternative choices are possible for Eq. S21. For example, one can define a model variant where the constraint on ppGpp transcriptional regulation given by a constant Q sector, $\phi_{\rm R} < \phi_{\rm R}^{\rm max}$, is made explicit, by defining

$$\omega_{\rm R} = \phi_{\rm R}^{\rm max} \frac{K_{\rm G}}{K_{\rm G} + G(t)}. \tag{S22}$$

In this model variant, the equations at steady states become

$$\epsilon = \tilde{\epsilon} \frac{\psi_{\rm A}}{\psi_{\rm A} + k_{\rm A}} \ , \tag{S23}$$

$$G = G^{\rm ref} \left( \frac{\tilde{\epsilon}}{\epsilon} - 1 \right) \ , \tag{S24}$$

$$\lambda = \epsilon \phi_R \ , \tag{S25}$$

$$\phi_R = \phi_{max} \frac{K_{\rm G}}{K_{\rm G} + G(t)} \ . \tag{S26}$$

Hence, $G$ affects the ribosomal and RNAP sectors in a more symmetric way. Since we fit the empirical relation $\phi_{\rm R}(G)$ in order to define $\omega_{\rm R}$, our model appears to be sufficiently good for our purposes. Indeed, we verified that none of the main results of this study are affected by this variant. However, the model variant with bounded $\omega_{\rm R}$ leads to more consistent calculations and good agreement with data, and may be preferred in future studies.

The definition we have used for $\chi_{\rm i}$ relies on a set of simplifying assumptions that may not hold in some conditions. Specifically, the following requirements need to be fulfilled:

- For an accurate representation of ppGpp levels in transcript dynamics, both $T$ and $T_{\rm R}$ need to be sufficiently large. Otherwise, the ratio $\frac{T_{\rm R}}{T}$ becomes an approximation of the true allocation strategy.

- The number of ribosomes to be larger than the number of transcripts. This is required because $\frac{T_R}{T}$ is the probability of a ribosome hitting a ribosomal transcript, and for the law of large numbers, this probability is accurately achieved just for a sufficient amount of ribosomes.

- Conversely, the number of ribosomes should not be excessively large, as our model does not account for ribosomal traffic on the transcripts.

In summary, we have $\chi_{\rm R} = \frac{T_{\rm R}}{T}$ only when the following conditions hold: $LT \gg R \gg T$, where $L$ represents the maximum number of ribosomes that can be allocated to a transcript, and $T$ must be sufficiently large to accurately capture ppGpp changes. In other regimes, more general definitions $\chi_{\rm R}$ can be used [20], keeping the rest of the model ingredients similar.

Finally, it is important to emphasize that our framework exclusively focuses on ribosomal proteins, leaving ribosomal RNA implicit. Given the tightly coordinated production of ribosomal proteins and rRNA [21–23], this approximation appears reasonable. However, it is crucial to highlight that in our framework, $\omega_{\mathrm{R}}$ (representing the fraction of RNAP involved in transcribing ribosomal protein transcripts) is computed by considering only the RNA polymerases engaged in the transcription of protein transcripts, excluding those involved in rRNA synthesis. We chose this approach due to the analogy between RNAP sectors and protein sectors, which enables us to equate $\omega_{\mathrm{i}}$ with $\phi_{\mathrm{i}}$ during steady state. Nonetheless, it is worth noting that other studies also incorporate rRNA explicitly [24–26], hence it is in principle feasible to study this model extension, which would make sense in order to study conditions or perturbations where the balance assumed here is or can be lost. Similar considerations hold for representing tRNA biosynthesis in the model [26].

#### E. The dynamics of protein sectors are determined by their regulatory function and the growth rate

The derivation of sector dynamics in our model is straightforward and follows from their definition, precisely as for the FCR model [2]. Here are the main steps of the derivation. Each sector is characterized by the mass ratio of the proteins belonging to the sector to the total protein mass, $\phi_{\mathrm{i}} = M_{\mathrm{i}}/M_{\mathrm{tot}}$. The increment of the total mass is governed by biosynthesis

$$\frac{\mathrm{d}M_{\mathrm{tot}}(t)}{\mathrm{dt}} = \epsilon(t) M_{\mathrm{R}}(t) \ , \tag{S27}$$

while the increment of each sector mass fraction corresponds to a fraction of the total increment, as follows,

$$\frac{\mathrm{d}M_{\mathrm{i}}(t)}{\mathrm{dt}} = \chi_{\mathrm{i}}(t)\epsilon(t) M_{\mathrm{R}}(t). \tag{S28}$$

Combining these equations leads to the differential equation describing the dynamics of each sector

$$\frac{\mathrm{d}\phi_{\mathrm{i}}(t)}{\mathrm{dt}} = \lambda(t)[\chi_{\mathrm{i}}(t) - \phi_{\mathrm{i}}(t)] \ , \tag{S29}$$

where $\lambda$, by definition, is given by $\frac{1}{M_{\mathrm{tot}}}\frac{\mathrm{d}M_{\mathrm{tot}}}{\mathrm{dt}} = \epsilon(t)\phi_{\mathrm{R}}(t)$.

In this study, we focus on three sectors: the ribosomal sector $\phi_{\mathrm{R}}$, the constitutive sector $\phi_{\mathrm{P}}$, and the housekeeping sector $\phi_{\mathrm{Q}}$. As $\phi_{\mathrm{Q}}$ is assumed to be constant, the other two sectors must satisfy the constraint $\phi_{\mathrm{P}}(t) = 1 - \phi_{\mathrm{Q}} - \phi_{\mathrm{R}}(t) = \phi_{\mathrm{R,max}} - \phi_{\mathrm{R}}(t)$ at each time $t$. This constraint also applies to the regulatory functions $\chi_{\mathrm{R}}$ and $\chi_{\mathrm{P}}$, as well as the RNAP partition functions $\omega_{\mathrm{R}}$ and $\omega_{\mathrm{P}}$. This constraint holds for all these quantities simply because of mass conservation.

Thus, we can avoid including $\omega_\mathrm{P}$ and $\chi_\mathrm{P}$ in the simulation and use the relation between $\phi_\mathrm{R}$ and $\phi_\mathrm{P}$ to obtain the dynamics of the latter.

### 3. SUPPLEMENTARY NOTE 3: MODEL VERSION WITH INACTIVE RIBOSOMES AND DEGRADATION.

As mentioned in the main text, the model described so far is not adapted to describe slow-growth behavior due to the significance of protein degradation and inactive ribosomes in cellular physiology [15, 16]. Therefore, to describe this regime, we need to incorporate these two ingredients into the model. To achieve this, we follow ref. [16] and introduce two key elements: the fraction of ribosomes contributing to growth, denoted as $f_\mathrm{bg}$, and the degradation rate, denoted as $\eta$. Both quantities depend on the steady-state growth rate or, equivalently, on the nutrient quality $\nu$.

It is known that the fraction of active ribosomes decreases at slow growth rates and that when the growth rate is zero, $f_\mathrm{bg}$ must also be zero by definition. We first define the fraction of ribosomes contributing to growth as

$$f_\mathrm{bg}(\nu) = \frac{\nu}{\nu + \nu_\mathrm{bg}}. \tag{S30}$$

By this choice, following ref. [16], we propose a Michaelis-Menten dependence on nutrient quality, with $\nu_\mathrm{bg}$ as a parameter fitted to reproduce experimental slow-growth data on $\epsilon$ and $\phi_\mathrm{R}$. It is important to note that currently there is no available data on $f_\mathrm{bg}$, and the above equation represents an empirical approximation for this quantity.

Concerning the protein degradation rate, we have utilized the available data in *E. coli* [16, 27] to fit a hyperbolic function as follows,

$$\eta(\nu) = \frac{\eta_0 + \eta_\infty \nu}{1 + \nu} \;, \tag{S31}$$

where $\eta_0$ and $\eta_\infty$ represent the protein degradation rate values for $\lambda = 0$ and $\lambda = \infty$, respectively.

Once these two mechanisms are defined, we need to integrate them into the system model equations which now become:

$$\frac{\mathrm{d}\psi_\mathrm{A}(t)}{\mathrm{d}t} = \nu\phi_\mathrm{P}(t) - \epsilon(t)\phi_\mathrm{R}(t) f_\mathrm{bg}(\nu)(1 + \psi_\mathrm{A}(t)) + \eta(\nu)\psi_\mathrm{A}(t), \tag{S32}$$

$$\frac{\mathrm{d}\chi_\mathrm{R}(t)}{\mathrm{d}t} = \frac{1}{\tau_\chi}[\omega_\mathrm{R}(t) - \chi_\mathrm{R}(t)], \tag{S33}$$

$$\frac{\mathrm{d}\phi_\mathrm{R}(t)}{\mathrm{d}t} = \epsilon(t)\phi_\mathrm{R}(t) f_\mathrm{bg}(\nu)[\phi_\mathrm{R}(t) - \chi_\mathrm{R}(t)]. \tag{S34}$$

$$\lambda(t) = \epsilon(t)\phi_{\mathrm{R}}(t) f_{\mathrm{bg}}(\nu) - \eta(\nu). \tag{S35}$$

It should be noted that the structure of the equations remains the same, but in Eq. S32, an additional positive term representing protein degradation is included. Furthermore, both Eq. S32 and S34 multiply the ribosome sector by $f_{\mathrm{bg}}$, which is a consequence of the revised definition of the growth rate presented in Eq. S35, incorporating both active ribosomes and the degradation term. The other equations in the system remain unchanged. Note that the definitions provided for $f_{\mathrm{bg}}$ and $\eta$ are suitable just for studying steady-state behavior. During a shift, one would expect these two quantities, along with other relevant variables, to exhibit oscillatory behavior. However, our current definition predicts an immediate shift to the new post-shift equilibrium value. This is a consequence of the dependence on nutrient quality $\nu$, which changes in a stepwise fashion.

Figure 3 shows the predictions of this modified model. Specifically, the plots demonstrate that by incorporating the mechanisms of inactive ribosomes and degradation, the model can accurately reproduce the data even during the slow growth regime. A previous study [16] has already pointed out that degradation is necessary in order to explain the phenomenology of slow growth because the degradation (and the subsequent maintenance synthesis) is required to explain why at zero growth rate one still has measure a non-zero elongation rate [6, 15]. At the light of the current model, we can add that the maintenance activity of the ribosomes is also required to explain the observed ppGpp pool. Indeed, if we assume that the production and degradation of ppGpp is related to ribosomal activity, as claimed in ref. [6], we need to acknowledge that at zero growth rate the fraction of active ribosomes cannot go to zero, which means that the ribosomes cannot be all sequestered. If this were the case, the translation elongation rate $\epsilon$ would go to zero in our model and the ppGpp level, according to Eq. S16, would diverge, which clearly appears as a non-physical solution.

### 4. SUPPLEMENTARY NOTE 4: MODEL VERSION WITH INSTANTANEOUS TRANSCRIPTION.

This section presents a variant of the model that neglects the transcriptional delay due to ppGpp-induced changes in ribosome transcription, which we used to verify if the predicted oscillatory behavior arises solely due to the delay introduced by the rearrangement of the transcripts.

In the absence of transcription delays, the regulatory function $\chi_{\mathrm{R}}$ directly depends on the ppGpp levels, resulting in the following equation:

$$\chi_{\mathrm{R}}(t) = \omega_{\mathrm{R}}(t) = \frac{K_{\mathrm{G}}}{K_{\mathrm{G}} + G(t)} \,. \tag{S36}$$

Under this assumption, the system is reduced to just two ordinary differential equations

$$\frac{\mathrm{d}\psi_{\mathrm{A}}(t)}{\mathrm{dt}} = \nu\phi_{\mathrm{P}}(t) - \epsilon(t)\phi_{\mathrm{R}}(t) - \lambda(t)\psi_{\mathrm{A}}(t), \tag{S37}$$

$$\frac{\mathrm{d}\phi_{\mathrm{R}}(t)}{\mathrm{dt}} = \lambda(t)[\omega_{\mathrm{R}}(t) - \phi_{\mathrm{R}}(t)]. \tag{S38}$$

All the other equations in the model remain unchanged.

### 5. SUPPLEMENTARY NOTE 5: MODEL VERSION WITH PPGPP SYNTHESIS ONLY FROM RELA.

Before the direct measurement of the relationship between ppGpp levels and the translation elongation rate $\epsilon$ (Eq. S16) in ref. [6], the standard approach to describing ppGpp dynamics relied on the elongation-coupled ppGpp synthesis by the enzyme RelA, for which there is solid experimental evidence [28]. Specifically, ppGpp is synthesized by RelA using ATP and GTP as substrates in the presence of uncharged tRNAs. When uncharged tRNAs are loaded onto transcribing ribosomes, the ribosome becomes stalled on the transcript and RelA detects this and attaches to the ribosome, producing (p)ppGpp before detaching. This mechanism links ppGpp production to the state of central precursors, in particular, RelA can sense amino acid starvation (which influences the level of uncharged tRNAs) [3]. In contrast, less is known about how SpoT synthesizes ppGpp and how it senses stressful conditions. It appears that SpoT can be involved in the accumulation of ppGpp under various stressful conditions and nutrient limitations, not necessarily related to amino acid starvation [3, 19]. Therefore, this pathway is typically modeled simply as a degradation term [7, 8].

This information about RelA and the assumption that SpoT works at a constant rate leads to the following equation for ppGpp [5, 7]

$$\frac{\mathrm{d}G}{\mathrm{dt}} = k_{\mathrm{RelA}} f_{\mathrm{c}} \phi_{\mathrm{R}} (1 - P_{\mathrm{C}}) - k_{\mathrm{spoT}} G, \tag{S39}$$

where the first term on the right-hand side describes the synthesis by RelA and the second term represents the degradation by SpoT. We observe that the RelA term depends on both the amount of ribosomes $\phi_{\mathrm{R}}$ and the probability of having a charged tRNA $P_{\mathrm{C}} = \frac{\psi_{\mathrm{A}}}{\psi_{\mathrm{A}} + k_{\mathrm{A}}}$, while $k_{\mathrm{RelA}}$ and $f_{\mathrm{c}}$ are the synthesis constant and the conversion factor between the number of ribosomes and the sector size, respectively (their values can be found in Table 1). Assuming a quasi-steady-state description of ppGpp, the equation reduces to

$$G = \frac{k_{\mathrm{RelA}}}{k_{\mathrm{spoT}}} f_{\mathrm{c}} \phi_{\mathrm{R}} (1 - P_{\mathrm{C}}) \ , \tag{S40}$$

which differs significantly from the equation used here and by Wu and colleagues [6]. This fundamental difference falsifies this model, as shown in Supplementary Fig.4D, and suggests that the role of SpoT cannot be reduced to a simple degradation term (although the exact mechanism of SpoT is not yet known). Moreover, in this model, the relation between the ppGpp concentration and ribosome allocation is given by

$$\chi_{\mathrm{R}} = \frac{K_{\mathrm{G}}}{K_{\mathrm{G}} + G} \tag{S41}$$

following the definition used in ref. [7], where $K_{\mathrm{G}}$ is the same as in eq. 12 of the main text.

While the specific quantitative details may vary, it is worth mentioning that even when incorporating Marr's proposed model into our framework, the oscillatory behavior still emerges. Hence our main results appear to be more generic than these specific aspects of the model. As discussed in the main text, as long as there is a feedback network between amino acids and ribosomes, an oscillatory regime is expected, as shown in Supplementary Fig. 4E-H.

### 6. SUPPLEMENTARY NOTE 6: VARIANT OF THE FCR MODEL INCORPORATING A DESCRIPTION OF PPGPP.

In the main text, we presented the predicted shift behavior based on the FCR model [2]. Since the original version of the FCR model does not incorporate the dynamics of ppGpp, in order to get a prediction for the ppGpp levels within this framework we defined an extended version of the FCR model by integrating the findings from Wu and coauthors [6] to account for the ppGpp description.

Before introducing the equations that define the extended model, it is important to highlight a few key points. In the FCR model, the parameter $\sigma$ represents the ribosomal activity, defined as $\lambda/\phi_{\mathrm{R}}$, which is different from the translation elongation rate $\epsilon$. This distinction becomes significant here because the model for ppGpp, as presented in ref. [6], also introduces sequestered ribosomes, denoted as $\phi_{\mathrm{R,seq}}$, which are ribosomes that are synthesized but do not contribute to growth. Given that the peculiarity of this model is to define the sectors regulatory functions $\chi_{\mathrm{i}}$ as function of $\sigma$, it becomes important to introduce both $\sigma$ and $\epsilon$, which is not strictly necessary in our framework.

Let us proceed by presenting the main equations used in our extended FCR model with a brief explanation, given that most of them are already presented in ref. [2]. We start with the growth rate, which is determined by the nutrient uptake fluxes:

$$\lambda(t) = \mu_1 \phi_{\text{cat1}}(t) + \mu_2 \phi_{\text{cat2}}(t) \ , \tag{S42}$$

where $\phi_{\text{cat}}$ 1 and 2 are the uptake sectors for the first and the second carbon source present in the substrate, and $\mu$ 1 and 2 are their nutrient qualities. From the growth rate we define the ribosomal activity

$$\sigma(t) = \frac{\lambda(t)}{\phi_{\text{R}}(t)} \ . \tag{S43}$$

Next, we want to define the translation elongation rate. To do so we introduce the fraction of sequestered ribosomes

$$f_{\text{seq}}(t) = k_{\text{conv}} \left( \frac{1 - \frac{\sigma(t)}{\gamma}}{\phi_{\text{R}}^{\text{min}}} \right)^2 \ . \tag{S44}$$

With this definition, we define the amount of sequestered ribosomes

$$\phi_{\text{R,seq}}(t) = \phi_{\text{R}}(t) f_{\text{seq}}(t) \ , \tag{S45}$$

and the translation elongation rate, following the definition in ref. [6]

$$\epsilon(t) = \frac{\lambda(t)}{\phi_{\text{R}}(t) - \phi_{\text{R,seq}}(t)}. \tag{S46}$$

The ppGpp dynamics follows the rules defined by Wu and coauthors, as in our model,

$$G(t) = CG^{\text{ref}} \left( \frac{\tilde{\epsilon}}{\epsilon(t)} - 1 \right). \tag{S47}$$

The two regulatory functions are given by steady-state relations [2]

$$\chi_{\text{R}}(t) = \chi_{\text{R}}^*(\sigma(t)), \tag{S48}$$

$$\chi_{\text{cat}}(t) = \chi_{\text{cat}}^*(\sigma(t)), \tag{S49}$$

and, as usual, the dynamics of the sectors are determined by the regulatory functions

$$\frac{d\phi_{\text{R}}(t)}{dt} = \lambda(t)[\chi_{\text{R}}(t) - \phi_{\text{R}}(t)], \tag{S50}$$

$$\frac{d\phi_{\text{cat}}(t)}{dt} = \lambda(t)[\chi_{\text{cat}}(t) - \phi_{\text{cat}}(t)]. \tag{S51}$$

Most of the equations presented here are directly taken from the FCR [2], with the addition of the ppGpp equation from Wu et al. [6]. However, we have introduced two new equations that describe the behavior of sequestered ribosomes during a shift, namely Eq.s S44 and S45. We have defined this extension in order to make testable predictions for ppGpp during shifts. While the authors of ref. [6] do not explicitly describe how sequestered ribosomes behave during a shift and only state that the sequestering process is independent of protein synthesis, we propose an ansatz to maintain the steady-state relationship between ribosomal activity $\sigma$ and translation elongation rate $\epsilon$ as described in ref. [6],

$$\sigma(t) = \epsilon(t)\frac{\phi_{\mathrm{R}}(t) - \phi_{\mathrm{R,seq}}(t)}{\phi_{\mathrm{R}}(t)}. \tag{S52}$$

During a shift, $\sigma$ changes faster than $\phi_{\mathrm{R}}$, and in order to maintain the correspondence between $\sigma$ and $\epsilon$, it is necessary for the sequestered ribosomes to change as fast as $\sigma$ does. To account for this, we model $f_{\mathrm{seq}}$ as a dynamic function of $\sigma$. Let us explain how we describe the sequestered ribosomes. According to ref. [6], the sequestered ribosomes constitute a fraction of the total number of ribosomes at steady-state. Using the equations $\phi_{\mathrm{R}} = \left(\frac{A\eta_{\mathrm{R/P},\phi}}{\eta_{\mathrm{N,R/P}}}\right)/G$ and $\phi_{\mathrm{R,seq}} = \frac{B\eta_{\mathrm{R/P},\phi}}{\eta_{\mathrm{N,R/P}}}G$, where $G$ is the ppGpp level the rest are constants fitted from the growth law (see ref. [6] for details), we can write:

$$\phi_{\mathrm{R,seq}} = \frac{k_{\mathrm{conv}}}{\phi_{\mathrm{R}}}, \tag{S53}$$

where $k_{\mathrm{conv}} = \left(\frac{\eta_{\mathrm{R/P},\phi}}{\eta_{\mathrm{N,R/P}}}\right)^2 AB$. We want to express the sequestered ribosomes as $\phi_{\mathrm{R,seq}} = \phi_{\mathrm{R}} f_{\mathrm{seq}}$. By substituting the above equation, we obtain

$$f_{\mathrm{seq}}(\sigma) = \frac{k_{\mathrm{conv}}}{\phi_{\mathrm{R}}^2(\sigma)} = k_{\mathrm{conv}}\left(\frac{1-\frac{\sigma}{\gamma}}{\phi_{\mathrm{R}}^{\min}}\right)^2, \tag{S54}$$

where we have explicitly included the dependence on ribosomal activity $\sigma$ by using the first growth law [2].

We have defined this updated version of the FCR framework in order to compare its prediction with the ones of our model. The prediction and the dynamics of the updated FCR model are presented in Supplementary Fig. 5, which shows that this model reproduces the observed dynamics, for both up and downshifts, as well as making predictions for additional observables. However, due to the steady-state assumptions intrinsic in the FCR model, it does not produce any damped oscillation in response to a nutrient change.

### 7. SUPPLEMENTARY NOTE 7: COMPARISON WITH OTHER MODELS.

This section compares our framework with two other models describing nutritional upshifts available in the literature, the Flux-Controlled Regulation (FCR) model [2], from which our framework derives, and the Flux Parity Model (FPM) introduced by Chure and coauthors in ref. [17].

As anticipated in the main text, one important aspect by which our framework deviates from the FCR model is that the FCR model assumes that the steady-growth relationship between the ribosomal activity $\sigma$ and the regulatory functions $\chi_{\rm i}$ is also valid in non-steady conditions and during shifts. This assumption allows the authors to connect these two quantities directly, avoiding the description of the ppGpp mechanistic circuits that implement the synthesis control. By contrast, our model includes explicitly the relation between the translation elongation rate and the ppGpp concentration $G$, and the transcriptional regulation leading to $\chi_{\rm i}$, which we use to make predictions about the ppGpp concentration and transcriptional reprogramming during the shift.

Another difference between the FCR model and our model is the description of the amino-acid pool. In the FCR model, the main quantities that control the shift dynamics are the catabolic and biosynthesis fluxes, which are related to the accumulation of amino acids. However, the FCR framework does not explicitly account for the amino acids in the description, and the external condition is described by the fluxes only. In our model, the translation elongation rate is directly connected with the amino acid level, which is justified by the tRNA sensing model.

In contrast, the FPM model follows an optimization principle for defining the control of biosynthesis, always maximizing the growth rate. This model presents the sensing of the charged tRNAs, a quantity that can be related to the amino acid pool in our model. An important aspect of our model is that by explicitly accounting for the sensing and regulatory modules, it does not assume any optimization principle [29].

Fig. 5 of the main text compares the predictions of the three models for a nutrient upshift, showing that, due to their different assumptions, they differ. The FCR model does not predict the amino acid pool and the FPM needs to be integrated with our hypothesis on the charged tRNA pool to do so. Furthermore, the FCR and FPM models do not predict the oscillatory relaxation toward the new steady state found by our model, as they do not explicitly describe the dynamics of the amino acid pool and its feedback interaction with the ribosomal proteins.

Supplementary Table 1: Parameters table

| Parameter | Value | Units | Reference |
|---|---|---|---|
| $\phi_{\mathrm{R}}^{\max}$ | 0.333 | dimensionless [mass fraction] | ref. [2] |
| $\phi_{\mathrm{R}}^{\min}$ | 0.049 | dimensionless [mass fraction] | ref. [2] |
| $\gamma$ | 11.02 | rate [$h^{-1}$] | ref. [2] |
| $\tilde{\epsilon}$ | 10.48 | rate [$h^{-1}$] | this work, from Eq. S13 |
| $k_{\mathrm{A}}$ | 2 | dimensionless [mass fraction] | fit of the experimental data shown in Fig. 6 |
| $\tau_{\chi}$ | 0.964 | time [$min$] | measured in ref [4] |
| $K_{\mathrm{G}}$ | 8.07476 | concentration [$\mu$ M] | fit of ref. [5] data (reported here in Supplementary Fig. 1) |
| $G^{\mathrm{ref}}$ | 55.73 | concentration [$\mu$ M] | data from ref. [5] (reported here in Supplementary Fig. 1) |
| $A$ | 1.85E-05 | concentration [mass fraction*$\mu$ M] | ref. [6] |
| $B$ | 2.11E-06 | inverse of concentration [mass fraction/$\mu M^{-1}$] | ref. [6] |
| $C$ | 4.6 | dimensionless [fold change] | ref. [6] |
| $\eta_{\mathrm{N,R/P}}$ | 6.8E-05 | dimensionless [mass fraction] | ref. [6] |
| $\eta_{\mathrm{R/P},\phi}$ | 0.46 | dimensionless | ref. [2] |
| $\eta_0$ | 0.05 | rate [$h^-1$] | fit of data reported in ref. [16] (original data from ref. [27]) |
| $\eta_\infty$ | 0.009 | rate [$h^-1$] | fit of data reported in ref. [16] (original data from ref. [27]) |
| $\nu_{\mathrm{bg}}$ | 1 | rate [$h^{-1}$] | this work, fit of data reported here in Fig 3 |
| $k_1$ | 1 | rate [$s^{-1}$] | ref. [7] |
| $k_2$ | 0.035 | rate [$s^{-1}$] | ref. [7] |
| $f_{\mathrm{c}}$ | 80 | $\mu M$ | ref. [9] (box 1) |
| $K_{\mathrm{G}}$ | 8.07476 | concentration [$\mu$ M] | fit of ref.s [5, 6] data, see Supplementary Fig. 1 |

### 8. SUPPLEMENTARY NOTE 8: CALCULATION OF FIXED POINTS.

In order to gain a deeper mathematical understanding of the system, it is desirable to obtain expressions for the fixed points of the dynamical system as functions of the parameters. It is particularly informative to examine these expressions in terms of the nutrient quality parameter, denoted as $\nu$, as it encapsulates the state of the environment and governs nutritional shifts when altered.

Since our system consists of three coupled differential equations, obtaining a closed analytical expression for the steady state becomes requires solving a cubic equation. Unfortunately, solving this cubic equation analytically yields lengthy and non-transparent solutions involving various combinations of the parameters.

Hence, obtaining a simple expression of the fixed points as solely a function of the parameters is a complicated task. An easier option is to compute the dependence on the growth rate $\lambda$, which can still be useful. At steady-state our system reads:

$$\epsilon = \tilde{\epsilon}\frac{\psi_\mathrm{A}}{\psi_\mathrm{A} + k_\mathrm{A}}, \tag{S55}$$

$$g = G_\mathrm{ref}C\left(\frac{\tilde{\epsilon}}{\epsilon} - 1\right), \tag{S56}$$

$$\lambda = \epsilon\phi_\mathrm{R}, \tag{S57}$$

$$\phi_\mathrm{R} = \frac{K_\mathrm{G}}{K_\mathrm{G} + G}. \tag{S58}$$

We can solve this system for $\phi_\mathrm{R}$:

$$\phi_\mathrm{R}(\lambda) = \frac{\lambda}{\tilde{\epsilon}}\frac{1 - K_\mathrm{G}/(G_\mathrm{ref}C)}{2}\left(1 + \sqrt{1 + \frac{4K_\mathrm{G}/(G_\mathrm{ref}C)}{\left(1 - K_\mathrm{G}/(G_\mathrm{ref}C)\right)^2}\frac{\tilde{\epsilon}}{\lambda}}\right), \tag{S59}$$

and use this expression to obtain the other fixed points as a function of $\lambda$:

$$\epsilon(\lambda) = \tilde{\epsilon}\frac{\phi_\mathrm{R}}{\phi_\mathrm{R} + \frac{K_\mathrm{G}}{G_\mathrm{ref}C}(1 - \phi_\mathrm{R})} = \frac{\lambda}{\phi_\mathrm{R}} = \frac{2}{1 - K_\mathrm{G}/(G_\mathrm{ref}C)}\frac{\tilde{\epsilon}}{1 + \sqrt{1 + \frac{4K_\mathrm{G}/(G_\mathrm{ref}C)}{(1 - K_\mathrm{G}/(G_\mathrm{ref}C))^2}\frac{\tilde{\epsilon}}{\lambda}}}, \tag{S60}$$

$$G(\lambda) = G_\mathrm{ref}C\left(\frac{\tilde{\epsilon}}{\epsilon} - 1\right) = G_\mathrm{ref}C\left(\phi_R\frac{\tilde{\epsilon}}{\lambda} - 1\right) = \frac{G_\mathrm{ref}C - K_\mathrm{G}}{2}\left(1 + \sqrt{1 + \frac{4K_\mathrm{G}/(G_\mathrm{ref}C)}{\left(1 - K_\mathrm{G}/(G_\mathrm{ref}C)\right)^2}\frac{\tilde{\epsilon}}{\lambda}}\right) - G_\mathrm{ref}C, \tag{S61}$$

$$\psi_\mathrm{A} = k_\mathrm{A}\frac{1}{\frac{\tilde{\epsilon}}{\epsilon} - 1} = k_\mathrm{A}\frac{1}{\frac{1 - K_\mathrm{G}/(G_\mathrm{ref}C)}{2}\left(1 + \sqrt{1 + \frac{4K_\mathrm{G}/(G_\mathrm{ref}C)}{(1 - K_\mathrm{G}/(G_\mathrm{ref}C))^2}\frac{\tilde{\epsilon}}{\lambda}}\right) - 1}. \tag{S62}$$

Let us comment briefly on these results. The expression for $\phi_{\rm R}(\lambda)$, Eq. S59, is the so-called "first growth law", and is usually modeled as a linear relation of the growth rate $\lambda$. Our solution highlights that the dependence, in our model, is not exactly linear, but there is a correction term given by the features of the ppGpp regulatory circuit.

The expressions of the translation elongation rate $\epsilon$ and the one of the amino acids pool $\psi_{\rm A}$ are also interesting. Eq. S60 shows that $\epsilon$ cannot reach the value $\tilde{\epsilon}$ unless the size of the ribosomal sector $\phi_{\rm R}$ reaches 1. This is empirically impossible given that the value of the maximum extension of $\phi_{\rm R}$ is about 0.55 (see 1 and ref.s [2, 10]). Therefore, in this model, the translational speed is always lower than $\tilde{\epsilon}$, which may seem counterintuitive especially if one just looks at Eq. S14. This fact has consequences on the amino-acid pool, visible in Eq. S62. By having $\epsilon < \tilde{\epsilon}$ regardless of the nutrient quality, this equation shows that the amino-acid pool size never diverges, even in those environments characterized by very rich nutrients. This implies that (within our model) the rate of cellular growth is always limited by the amount of available amino acids, and not just by the allocation strategy of cellular resources.

### 9. SUPPLEMENTARY NOTE 9: MATHEMATICAL STUDY OF THE APPEARANCE OF THE OSCILLATOR RELAXATION RESPONSE.

As described in the main text, in order to obtain oscillatory response (even in the case of damped oscillations), a negative feedback loop with a substantial delay is a necessary requirement [30]. In our system, such a feedback loop exists between the amino acids pool and the ribosomal mass fraction. However, the presence of feedback per se is not sufficient to guarantee oscillatory behavior, as in general the behavior depends on the values of the parameters. This section presents an analysis of our model's relaxation behavior, examining both systematically varying these parameters (close to biologically realistic regimes) to fully characterize the model's capabilities and limitations.

We start with the eigenvalues analysis of the system operating with the parameters values reported in biological literature (reported in Supplementary Table 1). As reported in the main text, for this set of parameters values we found just one relaxation behavior, which is the one showing damped oscillations. This finding is confirmed by the numerical study of the system eigenvalues: (I) all three eigenvalues show a negative real part, which ensures the convergence of the system to a new steady state after a perturbation, and (II) two eigenvalues present a non-zero imaginary part, which is responsible for the emergence of the oscillatory behavior [30]. The real and imaginary parts of the eigenvalues are plotted in Supplementary Fig. 6. The plot of the imaginary part shows that this goes to zero as also the nutrient quality reaches zero, implying that, for the kind of shift studied here, oscillations are always predicted.

We found that this is not true in general, and that with different sets of parameters, additional relaxation regimes arise. In the following, we present a more theoretical study of the system and of the 2×2 system variant, in order to get more insight into these theoretical predictions of the proposed framework.

*General requirement for stability.* We start performing stability analysis, linearizing the system around the fixed points and compute the eigenvalues of the dynamical system, which encode information on how the system relaxes upon weak perturbations of the steady state. For systems described by just two coupled ODEs (2×2 systems), the determinant and the trace of the Jacobian matrix correspond to the product and the sum of the eigenvalues, hence can be used to fully determine if the relaxation is oscillatory without computing directly the eigenvalues. Indeed in 2×2 systems (i.e. the system with instantaneous transcription described earlier), the determinant and trace of the Jacobian read

$$\det(J) = \frac{\partial \dot{\phi}_{\mathrm{R}}}{\partial \phi_{\mathrm{R}}} \frac{\partial \dot{\psi}_{\mathrm{A}}}{\partial \psi_{\mathrm{A}}} - \frac{\partial \dot{\phi}_{\mathrm{R}}}{\partial \psi_{\mathrm{A}}} \frac{\partial \dot{\psi}_{\mathrm{A}}}{\partial \phi_{\mathrm{R}}} \tag{S63}$$

$$\mathrm{Tr}(J) = \frac{\partial \dot{\phi}_{\mathrm{R}}}{\partial \phi_{\mathrm{R}}} + \frac{\partial \dot{\psi}_{\mathrm{A}}}{\partial \psi_{\mathrm{A}}} \tag{S64}$$

where the dot represents the time derivative of the quantity. Specifically, 2×2 system presents complex eigenvalues if and only if

$$4 \det(J) > \mathrm{Tr}(J)^2, \tag{S65}$$

if, in addition to this, the following relationship holds

$$\mathrm{Tr}(J) < 0 \wedge \det(J) > 0, \tag{S66}$$

then the fixed points are stable, therefore the oscillations are damped toward the new equilibrium point.

*Stability conditions of a generic regulation.* Before studying under which conditions Eq.s S65 and S66 hold for our system, we can study whether a system with an unspecified regulatory circuit presents stable fixed points. We can define a 2×2 system for translation regulation with a generic regulatory circuit as follows

$$\frac{\mathrm{d}\psi_{\mathrm{A}}}{\mathrm{dt}} = \nu \phi_{\mathrm{R}}^{\max} - \tilde{\epsilon} \frac{\psi_{\mathrm{A}}}{k_{\mathrm{A}} + \psi_{\mathrm{A}}} \phi_{\mathrm{R}} (1 + \psi_{\mathrm{A}}), \tag{S67}$$

$$\frac{\mathrm{d}\phi_{\mathrm{R}}}{\mathrm{dt}} = \tilde{\epsilon} \frac{\psi_{\mathrm{A}}}{k_{\mathrm{A}} + \psi_{\mathrm{A}}} \phi_{\mathrm{R}} \left( f(\psi_{\mathrm{A}}, \phi_{\mathrm{R}}) - \phi_{\mathrm{R}} \right) \tag{S68}$$

where the first equation corresponds to Eq. S9, because it is not directly tied to the regulation, while the second equation contains a generic function $f$, which can in principle depend on both $\phi_{\mathrm{R}}$ and $\psi_{\mathrm{A}}$, which sets the target size of the ribosomal sector. The partial derivatives of this generic system are

$$\frac{\partial\dot{\psi}_{\mathrm{A}}}{\partial\psi_{\mathrm{A}}} = -\phi_{\mathrm{R}}\tilde{\epsilon}\frac{\psi_{\mathrm{A}}}{k_{\mathrm{A}}+\psi_{\mathrm{A}}}\left(1+\frac{k_{\mathrm{A}}}{\psi_{\mathrm{A}}}\frac{1+\psi_{\mathrm{A}}}{k_{\mathrm{A}}+\psi_{\mathrm{A}}}\right), \tag{S69}$$

$$\frac{\partial\dot{\psi}_{\mathrm{A}}}{\partial\phi_{\mathrm{R}}} = -\nu-\tilde{\epsilon}\frac{\psi_{\mathrm{A}}}{k_{\mathrm{A}}+\psi_{\mathrm{A}}}(1+\psi_{\mathrm{A}}), \tag{S70}$$

$$\frac{\partial\dot{\phi}_{\mathrm{R}}}{\partial\phi_{\mathrm{R}}} = \phi_{\mathrm{R}}\tilde{\epsilon}\frac{\psi_{\mathrm{A}}}{k_{\mathrm{A}}+\psi_{\mathrm{A}}}\left(\frac{\partial f}{\partial\phi_{\mathrm{R}}}-1\right), \tag{S71}$$

$$\frac{\partial\dot{\phi}_{\mathrm{R}}}{\partial\psi_{\mathrm{A}}} = \phi_{\mathrm{R}}\tilde{\epsilon}\frac{\psi_{\mathrm{A}}}{k_{\mathrm{A}}+\psi_{\mathrm{A}}}\frac{\partial f}{\partial\psi_{\mathrm{A}}}. \tag{S72}$$

Note that, given that the above expressions need to be evaluated in a fixed point, some of them are already simplified (for example the terms $f-\phi_{\mathrm{R}}$ are set to zero). We will now study the positivity of the determinant and the negativity of the trace, starting by noticing that both $\frac{\partial\dot{\psi}_{\mathrm{A}}}{\partial\psi_{\mathrm{A}}}$ and $\frac{\partial\dot{\psi}_{\mathrm{A}}}{\partial\phi_{\mathrm{R}}}$ are negative regardless of the nature of $f$.

The condition on the trace implies

$$\frac{\partial\dot{\phi}_{\mathrm{R}}}{\partial\phi_{\mathrm{R}}} < -\frac{\partial\dot{\psi}_{\mathrm{A}}}{\partial\psi_{\mathrm{A}}} \tag{S73}$$

which is the most permissive condition. This condition is surely satisfied if $\frac{\partial f}{\partial\phi_{\mathrm{R}}} < 0$, which implies a regulation that is monotonically decreasing in $\phi_{\mathrm{R}}$. Moreover, in this case, there is also an homeostatic mechanism towards the target size of the sector. Indeed, if $\phi_{\mathrm{R}}$ is slightly lower than the target, the regulation will push for $\phi_{\mathrm{R}}$ to increase, and vice versa in the opposite case.

The condition on the determinant is more complicated because it involves more factors. We can rearrange the terms of the determinant and obtain

$$\frac{\partial\dot{\psi}_{\mathrm{A}}/\partial\psi_{\mathrm{A}}}{\partial\dot{\psi}_{\mathrm{A}}/\partial\phi_{\mathrm{R}}}\left(\frac{\partial f}{\partial\phi_{\mathrm{R}}}-1\right) < \frac{\partial f}{\partial\psi_{\mathrm{A}}}, \tag{S74}$$

which defines the least restrictive condition on $f$ in order to have a positive determinant. If we assume that $\frac{\partial f}{\partial\phi_{\mathrm{R}}} < 0$ (given by the condition on the trace), then the left-hand term of Eq. S74 becomes negative, therefore by requiring $\frac{\partial f}{\partial\psi_{\mathrm{A}}} > 0$ the above condition is satisfied. Also this latter condition has a physical interpretation, indeed, in this case, if the amino acids build up then the circuit will increase the target size of $\phi_{\mathrm{R}}$, and the increased biosynthesis will push back the amino acids pool size. We will not study the presence of oscillations in this generic system, as this requires many assumptions on $f$, making this study less generic that is intended to be.

*Stability conditions in the case with instantaneous transcription.* It is possible to gain more specific insight when applying the above requirements to our system or variants of it. In particular, this section considers the model variant with instantaneous transcription, which corresponds to a 2×2 system. For illustrative purposes we set the parameter $k_{\mathrm{A}} = 5\ 10^{-3}$ previously suggested in the literature [9]. This is different from the one consistently given by our fits of experimental data, but gives more distinct relaxation regimes, which are easier to visualize. A discussion of the stability and relaxation study for the complete system with biological parameters is present in the main text, and Supplementary Fig. 6 shows the eigenvalues of the system. Of particular interest is the study of the parametric dependence of the onset of the damped oscillatory response, which is the nutrient quality after which shifts present an oscillatory relaxation behavior instead of an overdamped one.

We start by pointing out that in this 2×2 model the fixed points are always stable, as the partial derivatives of the regulatory function fulfill the conditions found above for the generic system. In addition, we can insert the definition of the trace and the determinant in Eq. S65, and we find the following requirement for the presence of oscillations,

$$\left(\frac{\partial \dot{\psi}_{\mathrm{A}}}{\partial \psi_{\mathrm{A}}} - \frac{\partial \dot{\phi}_{\mathrm{R}}}{\partial \phi_{\mathrm{R}}}\right)^2 < 4\frac{\partial \dot{\psi}_{\mathrm{A}}}{\partial \phi_{\mathrm{R}}}\frac{\partial \dot{\phi}_{\mathrm{R}}}{\partial \psi_{\mathrm{A}}}. \tag{S75}$$

The left-hand side term of Eq. (S75) contains two characteristic time scales: $\frac{\partial \dot{\psi}_{\mathrm{A}}}{\partial \psi_{\mathrm{A}}}$ and $\frac{\partial \dot{\phi}_{\mathrm{R}}}{\partial \phi_{\mathrm{R}}}$, which are, respectively, the time scale of the reaction of $\psi_{\mathrm{A}}$ in response to changes in $\phi_{\mathrm{R}}$ and the reaction time scale of $\phi_{\mathrm{R}}$ in response to changes in $\psi_{\mathrm{A}}$ [30]. Similarly, the right-hand side term of Eq. (S75) can be interpreted as an interaction strength between the two quantities. For a system where the time scales of the feedback interactions match, the left-hand term will always be zero and therefore the system will always show damped oscillations during the relaxation to the new steady state.

The specific values of the system parameters determine whether or not inequality S75 holds, setting a boundary between damped oscillations and exponential relaxation in the phase diagram of the system. We can calculate the partial derivatives of the system:

$$\frac{\partial \dot{\psi}_{\mathrm{A}}}{\partial \psi_{\mathrm{A}}} = -\phi_{\mathrm{R}}\tilde{\epsilon}\frac{\psi_{\mathrm{A}}}{k_{\mathrm{A}} + \psi_{\mathrm{A}}}\left(1 + \frac{k_{\mathrm{A}}}{\psi_{\mathrm{A}}}\frac{1+\psi_{\mathrm{A}}}{k_{\mathrm{A}} + \psi_{\mathrm{A}}}\right), \tag{S76}$$

$$\frac{\partial \dot{\psi}_{\mathrm{A}}}{\partial \phi_{\mathrm{R}}} = -\nu - \tilde{\epsilon}\frac{\psi_{\mathrm{A}}}{k_{\mathrm{A}} + \psi_{\mathrm{A}}}(1 + \psi_{\mathrm{A}}), \tag{S77}$$

$$\frac{\partial \dot{\phi}_{\mathrm{R}}}{\partial \phi_{\mathrm{R}}} = -\phi_{\mathrm{R}}\tilde{\epsilon}\frac{\psi_{\mathrm{A}}}{k_{\mathrm{A}} + \psi_{\mathrm{A}}}, \tag{S78}$$

$$\frac{\partial \dot{\phi}_{\mathrm{R}}}{\partial \psi_{\mathrm{A}}} = \phi_{\mathrm{R}} \tilde{\epsilon} \frac{\psi_{\mathrm{A}}}{k_{\mathrm{A}} + \psi_{\mathrm{A}}} \frac{\frac{K_{\mathrm{G}}/(CG^{\mathrm{ref}})}{k_{\mathrm{A}}}}{(\frac{K_{\mathrm{G}}/(CG^{\mathrm{ref}})}{k_{\mathrm{A}}} \psi_{\mathrm{A}} + 1)^2}. \tag{S79}$$

Note that $\frac{\partial \dot{\phi}_{\mathrm{R}}}{\partial \psi_{\mathrm{A}}}$ depends solely on a single parameter, namely $\frac{K_{\mathrm{G}}/(CG^{\mathrm{ref}})}{k_{\mathrm{A}}}$, which represents the ratio of two scales. The first, denoted as $k_{\mathrm{A}}$, represents the concentration of amino acids at which the elongation rate $\epsilon$ attains half of its maximum value. The second, $K_{\mathrm{G}}/(CG^{\mathrm{ref}})$, corresponds to the ratio of the Michaelis-Menten constant of $\chi_{\mathrm{R}}(G)$, $K_{\mathrm{G}}$, governing the extent of ppGpp required for altering resource allocation, and the sensitivity of ppGpp concerning variations in the translation elongation rate $\epsilon$: $CG^{\mathrm{ref}}$. We refer to this factor as $\tilde{G} = K_{\mathrm{G}}/(CG^{\mathrm{ref}})$. This observation implies that the ribosomal resource allocation strategy is solely contingent on the relationship between these two scales. However, as we will demonstrate in this section, the overall system dynamics are more intricate.

By substituting the partial derivatives into Eq. S75, we derive the subsequent condition for $\nu$ to initiate oscillations,

$$\frac{\tilde{\epsilon} k_{\mathrm{A}} \tilde{G}}{4\phi_{\mathrm{R}}^{\max}} < \nu \frac{(\psi_{\mathrm{A}}(\nu) + k_{\mathrm{A}})^3}{\psi_{\mathrm{A}}(\nu)(1 + \psi_{\mathrm{A}}(\nu))^2}. \tag{S80}$$

Since Eq. S80 still incorporates a state variable (the amino acids pool $\psi_{\mathrm{A}}$), it is not possible to derive a phase diagram solely based on a single dimensionless parameter combination. Instead, we adopt the steady-state function fit $\psi_{\mathrm{A}}^*(\nu)$ to obtain the trends shown by Supplementary Fig. 8.

To acquire the function $\psi_{\mathrm{A}}^*(\nu)$, we chose to simulate the system's evolution for varying nutrient qualities ($\nu$), permitting the system to attain equilibrium. Subsequently, we collected the essential data, such as the equilibrium point coordinates $(\nu, \psi_{\mathrm{A}}, \phi_{\mathrm{R}}, \lambda)$, or the relationship between $\psi_{\mathrm{A}}$ and $\phi_{\mathrm{R}}$ concerning $\nu$. This data was then subjected to a polynomial fitting process, simplifying analysis and interpretation

*Dimensional analysis helps to characterize the phase diagram of damped oscillations.* Although we could only analyze Eq. S80 numerically, we can isolate several key trends by dimensional analysis, leveraging Eq. S80 to better characterize the phase diagram. We initiated this by plotting the right-hand side of Eq. S80 divided by $\nu$, as a function of $\nu$ itself. Figure 8A indicates that the RHS term can be approximated as linear in $\nu$, implying that it can be expressed as $\frac{(\psi_{\mathrm{A}}(\nu)+k_{\mathrm{A}})^3}{\psi_{\mathrm{A}}(\nu)(1+\psi_{\mathrm{A}}(\nu))^2} = f_{\mathrm{const}}(\tilde{G}, \tilde{\epsilon}, k_{\mathrm{A}})\nu$, where $f_{\mathrm{const}}$ represents a constant function of the system parameters.

To further break down the function $f_{\mathrm{const}}$, we analyzed the dimensions of the terms in

$$\frac{\tilde{\epsilon} k_{\mathrm{A}} \tilde{G}}{4\phi_{\mathrm{R}}^{\max}} < \nu^2 f_{\mathrm{const}}(\tilde{G}, \tilde{\epsilon}, k_{\mathrm{A}}), \tag{S81}$$

which is Eq.S80 with the given RHS ansatz. We note that the LHS dimension is $h^{-1}$ due to $\tilde{\epsilon}$, while the RHS dimension is $h^{-2}$ because of $\nu^2$. This implies that the RHS function must have the dimension of $h$, specifically $f_{\mathrm{const}}(\tilde{G}, \tilde{\epsilon}, k_{\mathrm{A}}) = h_{\mathrm{const}}(\tilde{G}, k_{\mathrm{A}})/\tilde{\epsilon}$.

Substituting this relation into the previous equation, we derive

$$\left(\frac{k_{\mathrm{A}}\tilde{G}}{4\phi_{\mathrm{R}}^{\max}}\right)/h_{\mathrm{const}}(\tilde{G}, k_{\mathrm{A}}) < \frac{\nu^2}{\tilde{\epsilon}^2}, \tag{S82}$$

signifying that $\nu^*/\tilde{\epsilon}$ must be a constant. This prediction is verified by panel B of Supplementary Fig. 8, illustrating that $\nu^*/\tilde{\epsilon}$ remains well approximated by a constant while altering the value of $\tilde{\epsilon}$.

Unfortunately, breaking down the dependence on the other parameters is not as straightforward. Nonetheless, we can use numerical simulations to show how $\nu^*/\tilde{\epsilon}$ changes with alterations in the parameter values (Supplementary Fig. 8C-D-E). In particular, we can analyze how $h_{\mathrm{const}}$ changes when the other parameters involved in Eq. S82 change, by extracting and plotting the function

$$h_{\mathrm{const}}(\tilde{G}, k_{\mathrm{A}}) = \left(\frac{k_{\mathrm{A}}\tilde{G}}{4\phi_{\mathrm{R}}^{\max}}\right)\frac{\tilde{\epsilon}^2}{(\nu^*)^2}, \tag{S83}$$

as a function of the parameters.

*Amino acid dynamics (and not only ppGpp regulation) contribute to the damped oscillations threshold.* Lastly, we aim to ascertain whether the effective parameter discovered earlier appears in $h_{\mathrm{const}}(\tilde{G}, k_{\mathrm{A}})$. To this aim, we examined the trend of $\nu^*/\tilde{\epsilon}$ in a scenario where $K_{\mathrm{G}}$ and $C$ change while maintaining $\tilde{G} = K_{\mathrm{G}}/(CG^{\mathrm{ref}})$ constant. Supplementary Fig. 8C demonstrates that the parameter $\tilde{G}$ indeed functions as an effective parameter, as $\nu^*/\tilde{\epsilon}$ remains constant even if $\tilde{G}$ changes.

In contrast, Supplementary Fig. 8D shows that even though the parameter combination $\tilde{G}/k_{\mathrm{A}}$ appears in the ribosomal regulatory function $\chi_{\mathrm{R}}$, it is insufficient to classify it as an effective parameter for the entire system. Indeed in our system, $k_{\mathrm{A}}$ also appears in the equation governing amino acid dynamics, but not in combination with $\tilde{G}$. Consequently, amino acid dynamics play a pivotal role in the emergence of oscillatory behavior, as this characteristic is not solely reliant on biosynthesis regulation.

*Different results for the model including transcriptional delays.* This Supplementary section has so far focused on the study of the reduced 2×2 dynamical system, because of the presence of simple criteria that ensure the stability of the fixed points and the nature of the relaxation. It is important to highlight that some of the considerations above do not apply to the complete model (a 3×3 dynamical system), which includes the transcriptional timescale for mRNA production, and hence should be more realistic.

For this complete model, there are no simple criteria that describe stability and relaxation, therefore, as mentioned in the main text, we performed a direct study of the eigenvalues of the system, shown in Supplementary Fig. 7, together with the eigenvalues of the 2×2 system with instantaneous transcription. For this analysis, we set again $k_{\mathrm{A}} = 5\ 10^{-3}$, as above. This analysis shows that our model exhibits a variety of behaviors depending on the parameters describing the nutritional upshift. From the analysis of the eigenvalues of the system, we can identify three regimes: (I) a shift without oscillations at slow growth rates, (II) a shift with damped oscillations in the fast growth regime, and (III) a shift with sustained oscillations (Hopf bifurcation) at growth rates far above the maximum achievable one experimentally [31]. The presence of sustained oscillations occurs when the real part of the complex eigenvalues becomes positive. These oscillations are shown in Supplementary Fig. 9. Note that all these three growth regimes are compatible with a system that can achieve balanced growth during steady-state [31]. As for the 2×2 system, the exact threshold between these regimes depends on the parameters of the system.

### 10. SUPPLEMENTARY NOTE 10: THE PROPOSED MODEL IS ANALOGOUS TO A NEGATIVE AUTOREGULATION.

In this section, we argue that the emergent feedback from the global regulation in our model can be paralleled to a negative autoregulation. A signature of the negative autoregulation is that the production rate of a quantity $x$ depends on the quantity itself, and it decreases as $x$ increases.

We will argue that this holds for $\phi_{\mathrm{R}}$, by focusing on the feedback loop between the amino acid pool $\psi_{\mathrm{A}}$ and the ribosomal sector size $\phi_{\mathrm{R}}$, described by Eq. S9 and S29. The steady-state expression of $\psi_{\mathrm{A}}$ reads

$$\psi_{\mathrm{A}}^{*} = \frac{\nu}{\lambda}(\phi_{\mathrm{R}}^{\max} - \phi_{\mathrm{R}}) - \frac{\epsilon}{\lambda}\phi_{\mathrm{R}} , \tag{S84}$$

which is a monotonic decreasing function of $\phi_{\mathrm{R}}$. We can insert this expression in the equation for $\chi_{\mathrm{R}}$, which is the production term of $\phi_{\mathrm{R}}$ (Eq. S29). We obtain the following expression

$$\chi_{\mathrm{R}} = K_{\mathrm{G}}\left(K_{\mathrm{G}} + G^{\mathrm{ref}}C\left(\frac{\psi_{\mathrm{A}} + k_{\mathrm{A}}}{\psi_{\mathrm{A}}} - 1\right)\right)^{-1} , \tag{S85}$$

which shows that the regulatory function $\chi_{\mathrm{R}}$ is monotonically increasing in the amino acid concentration $\psi_{\mathrm{A}}$. This follows from the fact that as the amino acids builds up, ppGpp production decreases, and the target for ribosome transcription increases. In turn, $\psi_{\mathrm{A}}^{*}$ decreases with increasing $\phi_{\mathrm{R}}$, and this makes the regulatory function a decreasing function of $\phi_{\mathrm{R}}$. For small circuits, negative autoregulation feedback loops were argued to provide quicker response to perturbation, and speculated to be selected based on this trait [32].

### 11. SUPPLEMENTARY NOTE 11: ANALYSIS OF EXPERIMENTAL DATA FROM PANLILIO ET AL. 2021.

In this work, we utilize the nutritional shift data reported in ref. [1]. The authors of that study investigate the dynamics of the shift at the single-cell level using a mother-machine microfluidic device. We leverage their data to compare the model predictions with experimental results. Figure 5 of the main text presents the outcomes of this comparative analysis.

#### A. Experimental apparatus and processed data

The microfluidic chip consists of parallel channels where the cells grow, with nutrients flowing from both the top and bottom of each channel. The data acquisition process used a microscope to capture images of approximately 10 channels per field of view simultaneously, with a time interval of 5 minutes between each capture. The acquired images were then processed using tracking software, which analyzed the cell growth on a frame-by-frame basis. The software recognized the cell's area and width and computed its volume using a sphero-cylindrical approximation. Panlilio and colleagues employed *E. coli* strains with fluorescent tags applied to the ribosomal (P1 and P1long) or constitutive promoters (P5). These tags were inserted near the replication origin ("ori" strains) or near the terminus ("ter" strains) of the genetic material. The constitutive P5 promoter is derived from the bacteriophage T5 [33]. On the other hand, the growth rate-dependent *E. coli* ribosomal promoter rrnB P1 is one of the two promoters responsible for initiating the transcription of the rRNA operon [34, 35]. The native P1 promoter contains several distinctive features, including a GC-rich discriminator region that renders it sensitive to the alarmone ppGpp [36, 37]. It also possesses an upstream sequence (UP element) that significantly enhances promoter activity [38]. Furthermore, the P1 promoter contains three Fis-binding sites located even further upstream, which further activate the promoter under nutrient-rich conditions [39–41]. P1long refers to the full-length rrnB P1 promoter, while P1 denotes the truncated version lacking the Fis-binding sites but still retaining the discriminator and UP element. It is important to note that the P5 promoter lacks the discriminator region, UP element, and Fis-binding sites. For a more detailed description of the promoters refer to ref.s [1, 42]. By using these fluorescently labeled strains, the authors obtained information on the fluorescence intensity, which is indicative of protein expression levels, throughout the entire experiment.

The performed shift was an upshift from minimal medium (M9) + glucose to M9 + glucose + casaminoacids [1]. The relevant processed data for our analysis consists of the time series data for each cell, including volume and fluorescence intensity were downloaded from the Mendeley Data repository associated to the study (DOI: 10.17632/5d4yhyjn8j.1). These data report on volume, cell-division events and fluorescence intensity of the tracked cell lineages.

### B. Computation of population data, growth rate, and sector size

We focused exclusively on the population-average time series of the growth rate and sector size (derived from the fluorescent reporters) across the shift. This section describes how we derived these average quantities from the single-cell data.

*a. Population averages.* We transformed the individual cell time series into population-average time series by calculating the average signal across all cells present at each experimental time interval using a sliding window average. The step size was 4 minutes, and the window size was 75 minutes, resulting in the coarse-grained population time series shown in Fig. 6 of the main text. In contrast, Supplementary Fig. 14 displays a less coarse-grained population time series, with a step size of 4 minutes and a window size of 20 minutes.

*b. Growth rate.* To compute the growth rate, we used the single-cell time series of the volume. We applied a Savitzky-Golay filter to the data and then calculated the growth rate as the central derivative of the volume with respect to time, divided by the volume

$$\lambda(t_{\mathrm{i}}) = \frac{1}{V(t)} \frac{V(t_{\mathrm{i+1}}) - V(t_{\mathrm{i-1}})}{t_{\mathrm{i+1}} - t_{\mathrm{i-1}}}. \tag{S86}$$

This was done for each cell, and the resulting growth rates were averaged across all cells as described above to obtain the population time series.

*c. Proteome sector size.* To extract information on the proteome sector size from the fluorescence signal, we first normalized the fluorescence intensity by the cell volume. This ratio is expected to be proportional to the protein concentration. To determine the conversion constant between the fluorescence signal and the proteome sector size, we used different procedures for the ribosomal and constitutive protein signals.

For the ribosomal signal, we used the growth law at steady state, along with the data on the growth rate, to find the conversion factor between the GFP signal normalized by the volume and the ribosomal sector size. This conversion factor was denoted as $k_{\mathrm{conv,R}}$ and was defined such that it satisfies

$$k_{\mathrm{conv,R}} \left( \frac{F_{\mathrm{p1}}}{V} \right)_{\mathrm{s1}} = \phi \mathrm{R}^{\mathrm{min}} + \frac{\lambda_{\mathrm{s1}}}{\gamma}, \tag{S87}$$

where the subscript $s1$ denotes the pre-shift steady state. The ribosomal time series was then obtained as the product of the conversion factor and the fluorescence signal normalized by the cell volume, $\phi_{\mathrm{R}}(t) = k_{\mathrm{conv,R}} \frac{F_{\mathrm{p1}}(t)}{V(t)}$.

For the constitutive protein signal, we utilized the constraint that the sum of the ribosomal and constitutive protein sector sizes is constant $\phi_{\mathrm{R}} + \phi_{\mathrm{P}} = 0.55$. We defined the conversion factor $k_{\mathrm{conv,P}}$ such that it satisfies

$$\phi_{\mathrm{R,s1}} + k_{\mathrm{conv,P}} \left( \frac{F_{\mathrm{p5}}}{V} \right)_{\mathrm{s1}} = 0.55. \tag{S88}$$

The constitutive protein time series was then obtained as the product of the conversion factor and the fluorescence signal normalized by the cell volume, $\phi \mathrm{P}(t) = k_{\mathrm{conv,P}} \frac{F_{\mathrm{p5}}(t)}{V(t)}$.

### C. Consistency checks on the fluorescence proxies

The approach used to compute the proteome sector size in this study is unique in the literature, as most previous studies have relied on macromolecular quantification or proteomic assays to assess the proteome [2, 10, 12, 43]. To ensure that using the fluorescence signal as a proxy for the proteome is valid, we conducted several consistency checks on the data.

Firstly, we examined whether the conversion factor $k_{\mathrm{conv,R}}$ computed using the pre-shift steady-state data remained consistent with the post-shift data. If different conversion factors were obtained for the two steady states within a time series, it would indicate a potential issue with either the steady states or the experiment itself. Panel A of Supplementary Fig. 15 shows that the conversion factor computed for the first steady state in the pair $(\lambda^{\mathrm{p5-ter}}, \phi_{\mathrm{R}}^{\mathrm{p1long-ori}})$ correctly places the point of the second steady state on the trend of the first growth law from literature data, as expected. However, the other pairs, where $\phi_{\mathrm{R}}^{\mathrm{p1long-ori}}$ is coupled with the growth rate from other strains (p1-ori, p1long-ori, and p1long-ter), did not meet this requirement. It is important to note that we excluded the fluorescence signal of the p1-ori and p1long-ter strains from this test for two main reasons: (I) the promoter P1long is the true ribosomal promoter as it conserves all the discriminatory regions and the "ori" position ensures that phenomena like gene dosage and supercoiling are taken into account, (II) they do not exhibit a post-shift steady state (shown in Supplementary Fig. 14B).

The second check we performed focused on the constitutive sector. We examined whether the constraint $\phi_{\mathrm{R}} + \phi_{\mathrm{P}} = 0.55$ held throughout the entire shift, despite the fact that the two sector fractions were derived from independent fluorescent reporters in different experiments. Panel B of Supplementary Fig. 15 demonstrates that this constraint holds during the entire shift, even if it is enforced on just the steady-state pre-shift data.

#### D. A cell-density imbalance alters the shift dynamics

The high temporal resolution of the data collected by Panlilio and colleagues exhibits a recently studied biophysical phenomenon occurring during amino acid upshifts, which goes beyond our model. Specifically, in the first minutes following the shift, the volume and mass of a cell become temporarily uncoupled. This feature appears in similar shifts that were studied recently monitoring synchronously both the volume and the refractive index of individual cells [44]. In the Panlilio et al. data, upon close examination of the growth rate data presented in Supplementary Fig. 14 and Supplementary Fig. 16, it is apparent that all the population-average time series exhibit a rapid overshoot in volume growth rate that is not matched by the protein dynamics. This discrepancy can be attributed to the osmotic imbalance that occurs immediately after the shift, as noted by Oldewurtel and colleagues [44].

This volume-mass growth imbalance has the potential to impact the shift dynamics by introducing a perturbation in the protein density. It is important to note that our model, as well as most shift models available in the literature, cannot account for this phenomenon, as a crucial assumption of most available models is that the protein density remains constant and that the sector sizes reflect the intracellular concentrations of the different proteins.

### SUPPLEMENTARY FIGURES

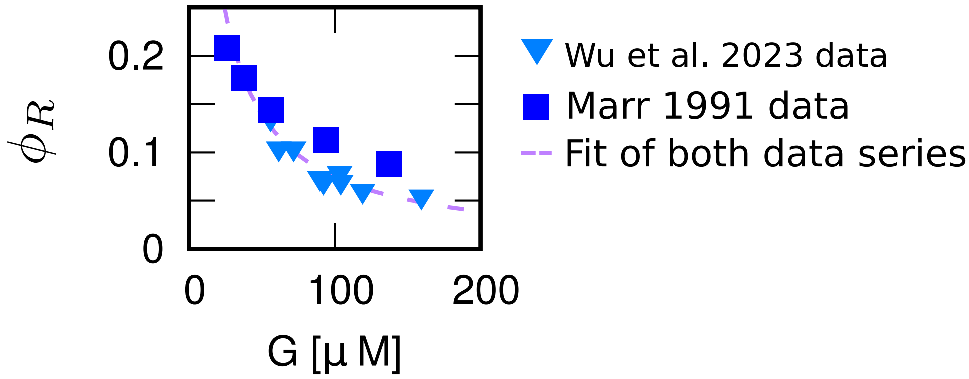


Supplementary Figure 1: **Fit of the dependency RNAP regulatory function $\omega_R$ from ppGpp concentration from experimental data.** The plot shows steady-state data from ref. [5, 6], the $y$ axis represents the ribosomal sector size, while the $x$ axis reports the ppGpp concentration. To obtain the plotted data, we have taken the data of ppGpp concentration versus the growth rate reported in ref. [5] and transformed the growth rate in the ribosomal sector size using the first growth law [2]. These data are shown as blue squares. Additionally, we have taken the data in ref [6] (blue reverse triangles), which reports the fold change of ppGpp levels versus the ribosomal sector size, and converted the fold change into absolute concentration using the data from ref. [5]. For the conversion, we used the value of ppGpp corresponding to a growth rate of approximately 1 1/h, which should be close to the growth on glucose that has been taken as a reference in ref. [6]: $G^{\mathrm{ref}} = 55.73\mu M$. The dashed line is a fit of both series of data.

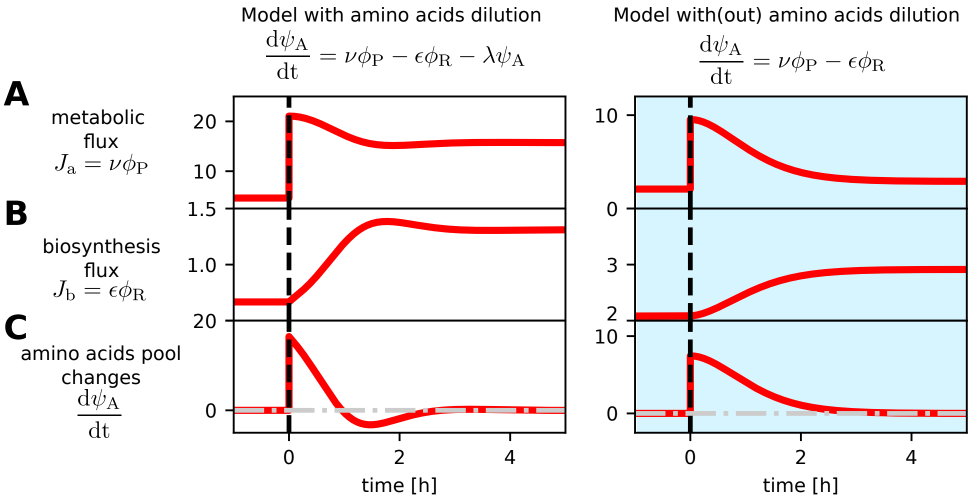


Supplementary Figure 2: **Neglecting the dilution of the amino acid pool affects the predicted adaptation.** The plots compare the dynamics of a model that considers amino acid dilution due to cell growth (main text, right panels) and a model that neglects it (left panels). In all panels, the $x$ axis represents the time from the shift (note that the span of the axis is different in the right and left panels). **A:** metabolic flux dynamics following an upshift. The metabolic flux, which supplies the amino acids pool, is given by the nutrient quality $\nu$ times the size of the constitutive sector $\phi_P$. We assume $\nu$ changes instantaneously over the shift, and so does the metabolic flux. **B:** biosynthesis flux dynamics following an upshift. This flux represents the protein production and is given by the translation elongation rate $\epsilon$ times the ribosomal sector size $\phi_R$. **C:** changes in the amino acids pool following an upshift. This plot shows the dynamics of the derivative of the size of the amino acids pool, which is also the difference between the two fluxes and, when considered, the dilution term (see equations at the top of the panels). The figure shows that both models reach equilibrium after being perturbed, contrary to what is shown in ref [17]. This happens because in our model the feedback between amino acid pool and ribosomes ensures that the disparity between the fluxes is adjusted by tuning the proteome composition, and this is true also for the model without dilution. However, if the dilution is neglected the system reaction to the perturbation changes.

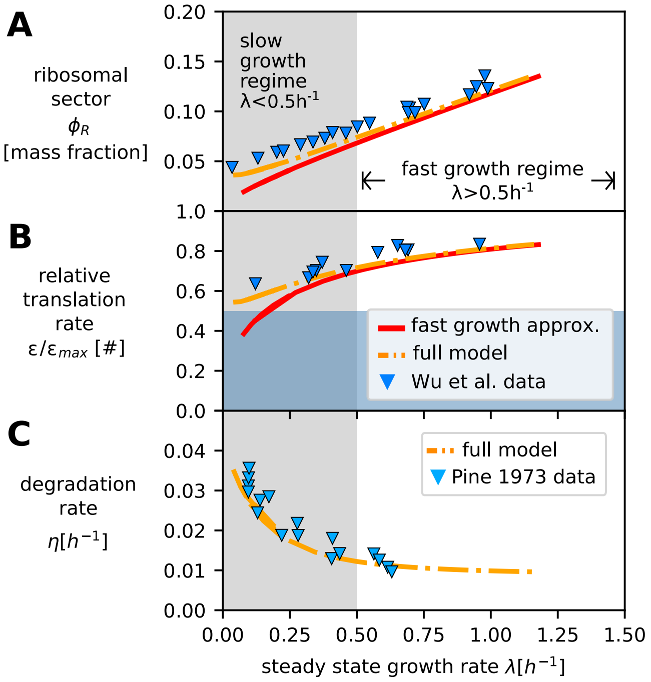


Supplementary Figure 3: **Adding degradation and inactive ribosomes is sufficient to quantitatively describe the slow growth regime.** The figure compares two versions of the proposed framework: the fast-growth approximation (red solid line) proposed in the main text and a full model that considers degradation and inactive ribosomes (orange dashed line). **A:** results for both models, along with experimental data from [6]. The $x$-axis indicates the steady-state exponential growth rate (shared with panels B-C), while the $y$-axis represents the size of the ribosomal sector. **B:** growth rate shown on the $x$-axis, with the relative translation rate shown on the y-axis. **C:** degradation rate vs growth rate, for the full model (orange dashed line) and for the data reported in [16, 27]. This panel shows an input of the full model, whereby degradation depends on growth rate in a non-linear fashion. The results obtained from the fast-growth model predict that $\phi_\mathrm{R} \to 0$ and $\epsilon \to 0$ as $\lambda \to 0$, which is in contrast with experimental evidence. However, both $\phi_\mathrm{R}$ and $\epsilon$ show a non-zero offset at zero growth rate when the full model is considered. This is explained by the fact that at slow growth rates ($\lambda < 0.5h^{-1}$), the phenomena of protein degradation and inactive ribosomes become relevant for growth physiology, as described in [16].

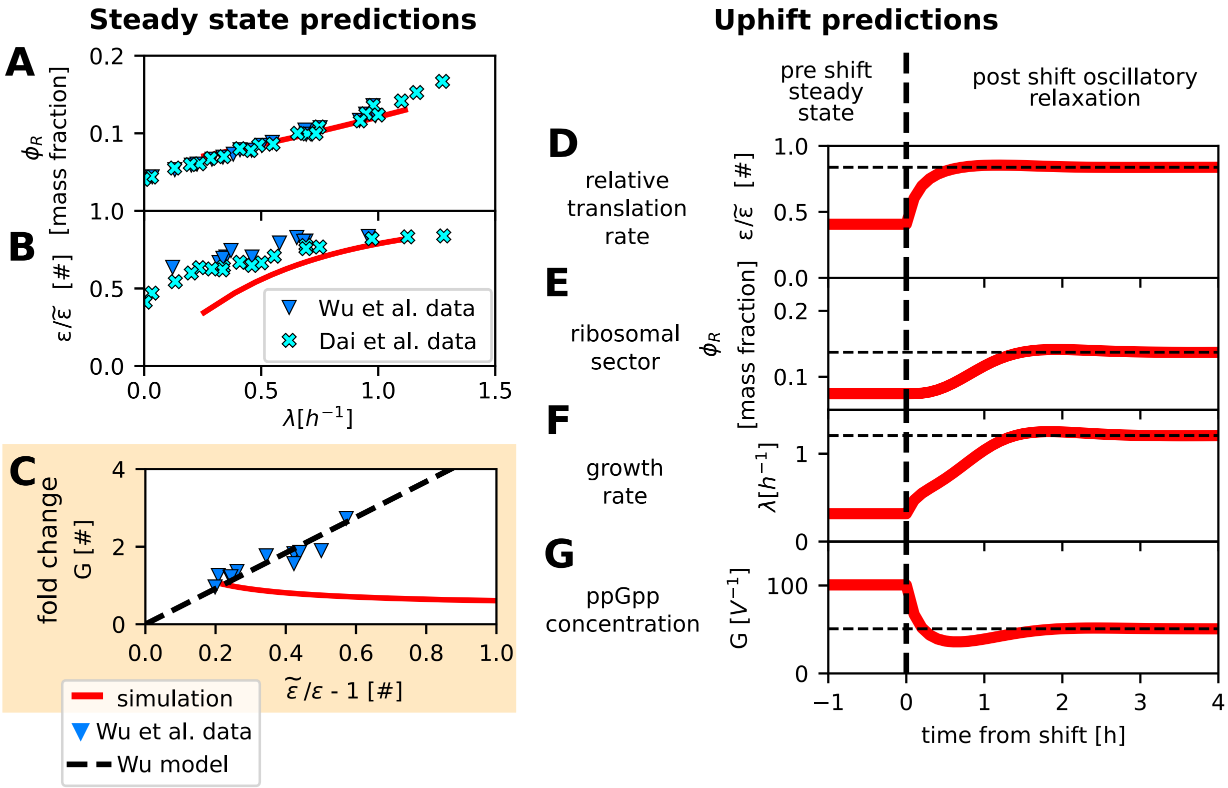


Supplementary Figure 4: **The model version that includes just the action of RelA still presents the oscillatory dynamics.** Panels **A-C** show the steady state prediction of this version of the model (red solid line) and experimental data from ref.s [15] (cyan crosses) and [6] (blue triangles), in particular, is plotted the ribosomal sector size (**A**) and the relative translation elongation rate (**B**) versus the growth rate, and the ppGpp concentration versus the inverse of the relative translation rate (**C**). These panels show that overall this model is performing well in the fast growth regime, except for the last prediction on ppGpp, which is the one that falsifies it. Panels **D-G** show the model predictions following an upshift (**D:** relative translation rate, **E:** ribosomal sector, **F:** growth rate, **G:** ppGpp concentration). These plots show that even if the relation between ppGpp and $\epsilon$ has changed the model still predicts an oscillatory relaxation towards the new steady state.

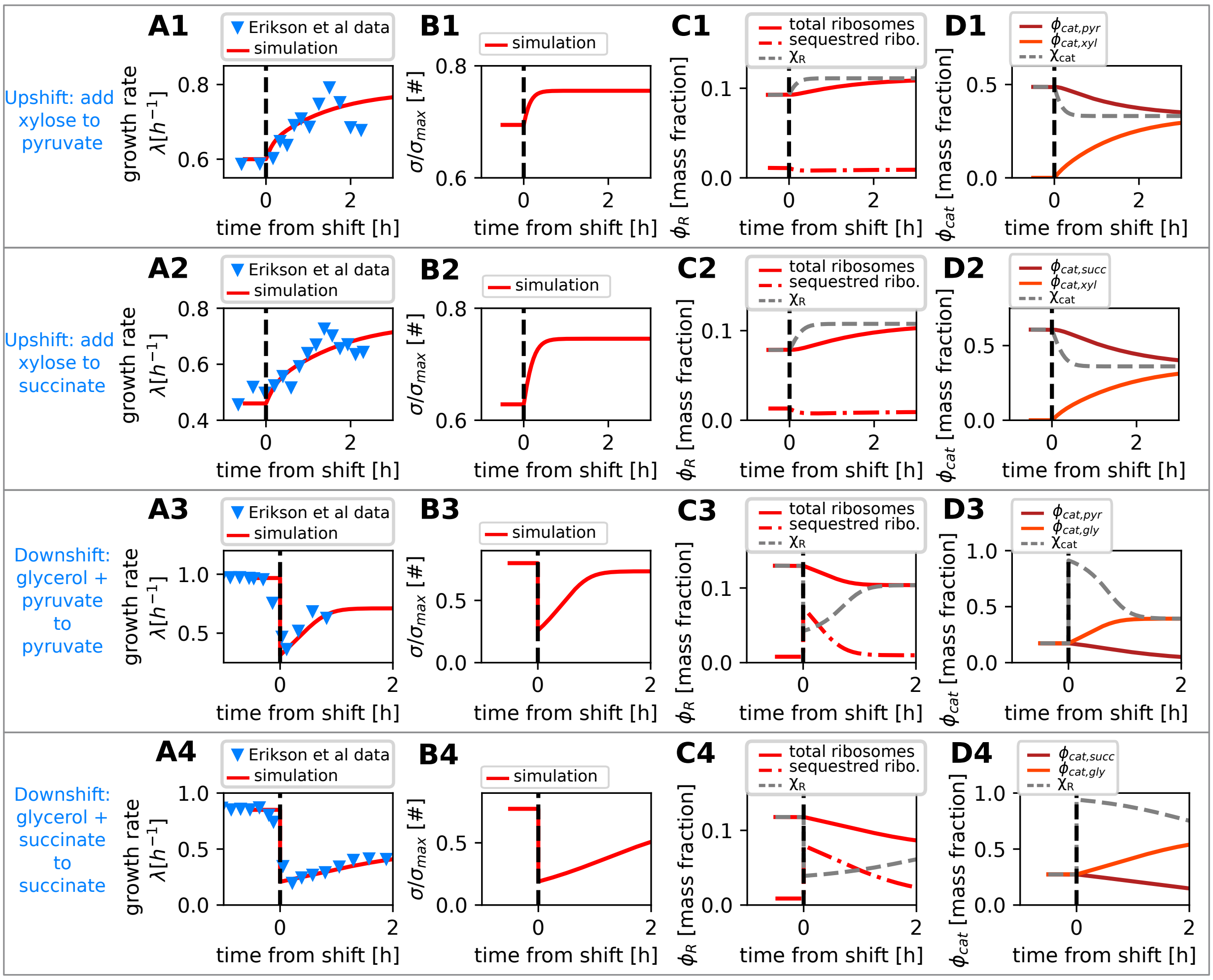


Supplementary Figure 5: **Performance of the FCR model variant with ppGpp dynamics.** The plot shows the dynamics of the principal quantities involved during nutrient shifts predicted by the FCR model equipped with the description of the ppGpp dynamics introduced in this work, and it compares with shift data collected in ref. [2]. The first two rows show the upshift dynamics, while the two bottom ones are the downshift one. All plots display the time from the shift on the x-axis. The first column (**A1-4**) shows the comparison between the predicted growth rate (red solid line) and the one observed in ref. [2] (blue reverse triangles). The second column (**B1-4**) shows the dynamics of the relative ribosomal activity $\sigma/\sigma_{\max}$, which rapidly increases during upshifts and decreases and recovers during downshifts. The third column (**C1-4**) shows the dynamics of the ribosomal sector and its regulatory function $\chi_{\rm R}$ (grey fine dashed line). In particular, we show the dynamics of the total ribosomes (red solid line) and of the sequestered ones (red dashed line), which are necessary in order to include the ppGpp dynamics, as described in these notes. The last column (**D1-4**) shows the dynamics of the catabolic sector $\phi_{\rm cat}$ (solid lines) and of its regulatory function $\chi_{\rm cat}$ (grey fine dashed line). This model describes a different uptake sector for each substrate present in the medium, therefore in this case we included two different sectors, one of which gets activated or diluted after an up or downshift, respectively.

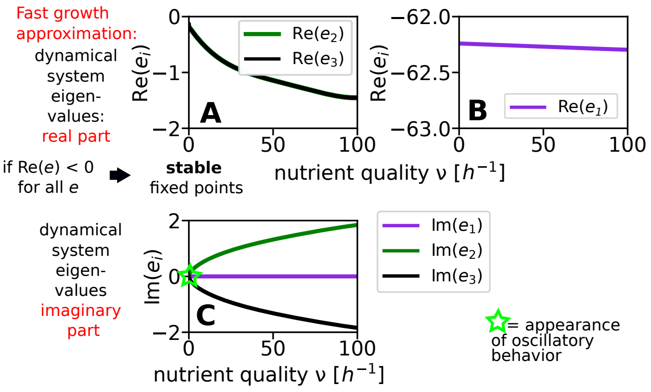


Supplementary Figure 6: **The eigenvalues from linear-stability analysis predict the presence of stable fixed points and oscillatory relaxation towards the new steady state.** In these simulations $k_A$ was set to 2 [mass fraction], the value that best reproduces the experimental data analyzed in Fig. 6. **A-B:** real part of the eigenvalues plotted as a function of the nutrient quality $\nu$. The real part of the eigenvalues is always negative in the relevant range of parameters, indicating stable fixed points. Panel **C** shows the imaginary parts of the eigenvalues of the models as a function of nutrient quality $\nu$. Panel C shows that for the imaginary part is always non-zero for every value of $\nu$, which implies the presence of damped oscillatory relaxation in every nutrient shift, regardless of the initial and final value of the growth rate.

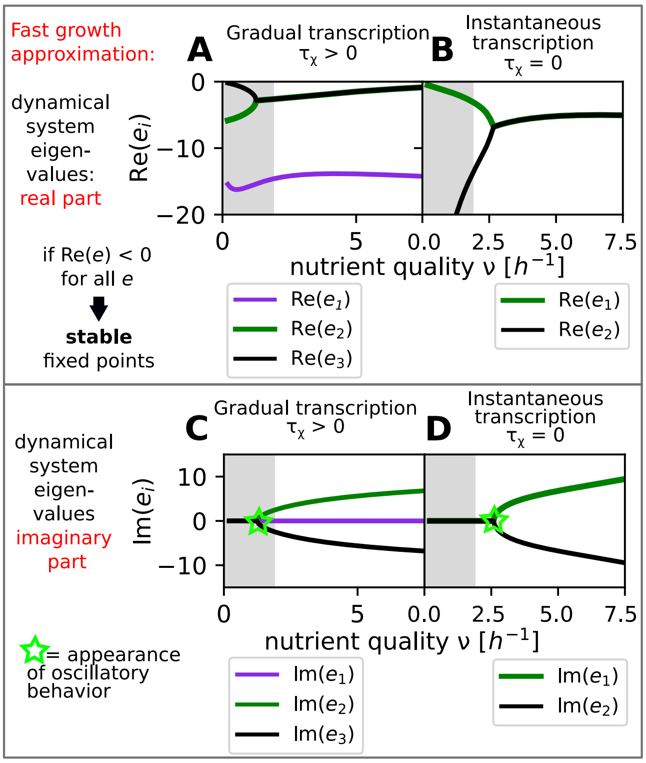


Supplementary Figure 7: **The eigenvalues from linear-stability analysis predict the presence three different relaxation regimes for the model outside the biological range of parameters.** This figure shows results for two variants of the model where $k_A$ was set to 5 $10^{-3}$, the one where transcriptional reprogramming from ppGpp has a timescale given by the degradation of the transcripts and a variant where transcription is instantaneous. **A:** real part of the eigenvalues plotted as a function of the nutrient quality $\nu$. The real part of the eigenvalues is always negative in the relevant range of parameters, indicating stable fixed points. **B:** the real part of the eigenvalues of the model variant with instantaneous transcription remains negative. Panels **C** and **D** show the imaginary parts of the eigenvalues of the instantaneous and gradual transcription models as a function of nutrient quality $\nu$. Above a threshold value of $\nu$, the eigenvalues become complex, leading to oscillatory relaxation. In all panels, the grey areas represent the slow-growth regime, defined as $\lambda < 0.5h^{-1}$, the dashed grey line is an empirical maximum growth rate $\lambda \simeq 2h^{-1}$.

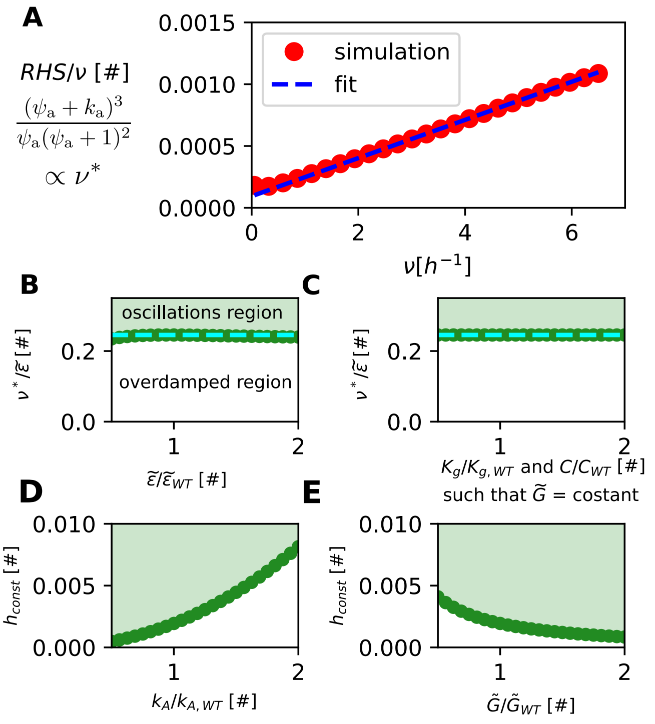


Supplementary Figure 8: **The onset of oscillatory behavior depends on key parameters such as maximum elongation speed and sensitivity of ppGpp to elongation rate.** These panels show the study of the threshold nutrient quality $\nu^*$ value triggering damped oscillations (a value that depends on the model parameters). **A:** the x-axis represents the nutrient quality $\nu$, while the y-axis shows the right-hand side of eq S80 (the definition id near the vertical axis). The red circles are the result of the numerical simulations of the RHS, while the blue dashed line is a linear fit of these points (for $\nu > 2$). This plot shows that for large $\nu$ the RHS can be approximated as a linear function of $\nu$. In all the subsequent panels we have studied the parameter dependence of $\nu^*$ in order to understand better the phase diagram of the system. The green points correspond to $\nu^*/\tilde{\epsilon}$ given the parameter values, the green shaded area indicates the parameter region where the model predicts oscillatory behavior, while the white area corresponds to overdamped relaxation. The explored range of all parameters spans from 0.5 to 2 times the value of the parameter estimated for the wild type, reported in table 1. **B:** trend of $\nu^*/\tilde{\epsilon}$ versus $\tilde{\epsilon}$. The panel shows that $\nu^*/\tilde{\epsilon}$ is almost constant in $\tilde{\epsilon}$, as predicted by the dimensionality study. **C:** $\nu^*/\tilde{\epsilon}$ is constant if $K_{\mathrm{G}}$ and $C$ are changed in such a way that $K_{\mathrm{G}}/(CG^{\mathrm{ref}}$ remains constant. This demonstrates that three parameters $K_{\mathrm{G}}$, $C$ and $G^{\mathrm{ref}}$ appears always in combination in Eq. S80. **D:** trend of $h_{\mathrm{const}}$ (Eq. S82) versus $k_{\mathrm{A}}$, the parameter that controls the sensitivity of the translation elongation rate to amino acid levels (Eq. S14). **E:** trend of $h_{\mathrm{const}}$ (Eq. S82) versus $\tilde{G}$.

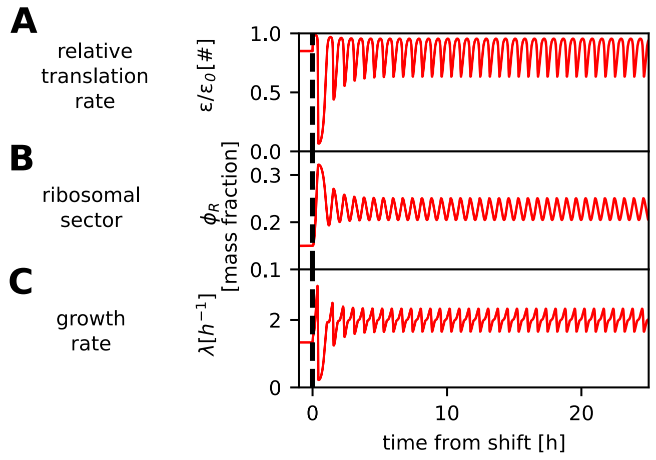


Supplementary Figure 9: **Presence of Hopf bifurcation in non-biological regimes.** All the plots are obtained by simulating an upshift in a regime where growth exceeds the maximum achievable growth rate. **A:** dynamics of the relative translation elongation rate $\epsilon/\tilde{\epsilon}$ across the shift. **B:** dynamics of the ribosomal sector size $\phi_R$ across the shift. **C:** dynamics of the instantaneous growth rate $\lambda$ across the shift. The plots show that in this regime the oscillations are not damped, and the system shows sustained oscillations around the new equilibrium. Mathematically, this happens because the real part of the complex eigenvalues becomes positive if the nutrient quality exceeds a certain threshold, as can be seen in Supplementary Fig. 7A. This phenomenon is called Hopf bifurcation in the theory of dynamical systems.

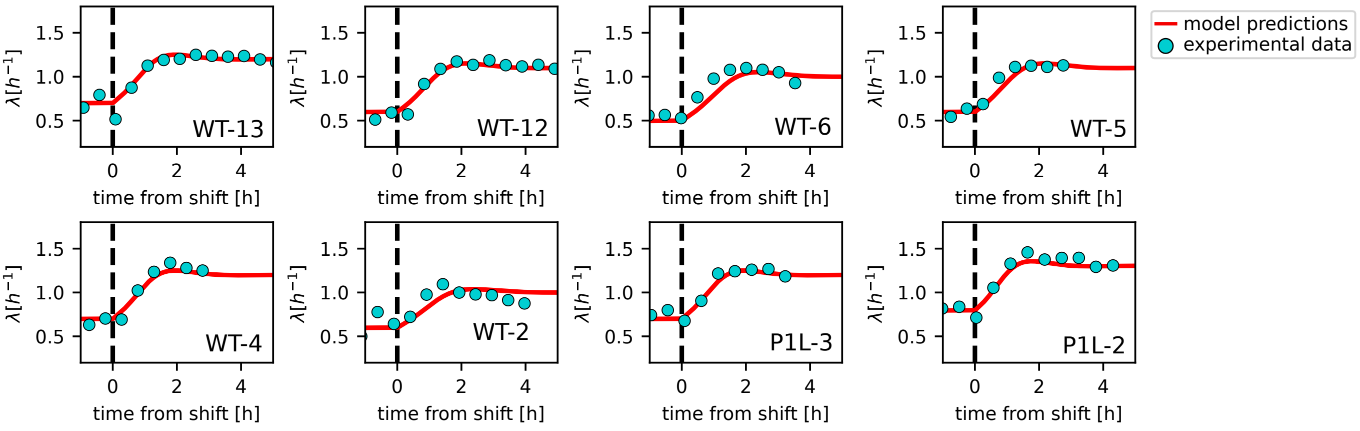


Supplementary Figure 10: **Batch nutrient upshifts showing overshoots that are compatible with the model predictions.** In all panels, the solid red line represents the model prediction (from simulations), with the initial and final growth rates adjusted to align with the experimental data. The blue circles indicate the average growth rates from the technical replicates of each experiment that successfully passed the filtering for the two criteria illustrated in the Methods section: stationary growth and coherence between technical replicates. The label in the bottom right corner specifies the strain and the index of the biological replicate. This figure includes the plots from all experiments that exhibited overshoots consistently with the theoretical predictions.

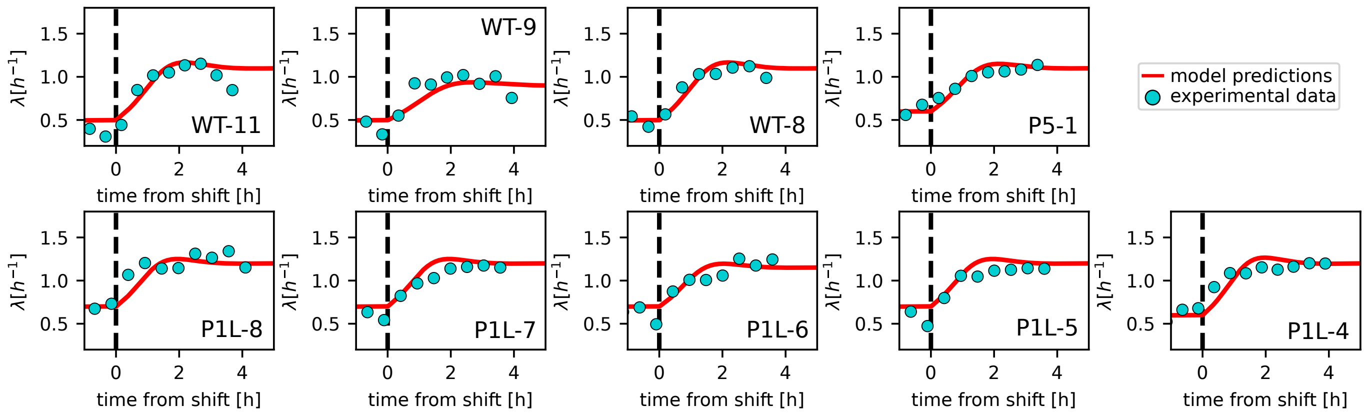


Supplementary Figure 11: **Batch nutrient upshift experiments that do not show visible overshoots, contrary to the model predictions.** In all panels, the solid red line represents the model prediction (from simulations), with the initial and final growth rates adjusted to align with the experimental data. The blue circles indicate the average growth rates from the technical replicates of each experiment that successfully passed the filtering for the two criteria illustrated in the Methods section: stationary growth and coherence between technical replicates. The label in the bottom right corner specifies the strain and the index of the biological replicate. This figure includes plots from all experiments where the data did not show clear overshoot oscillatory relaxation, contrary to model predictions.

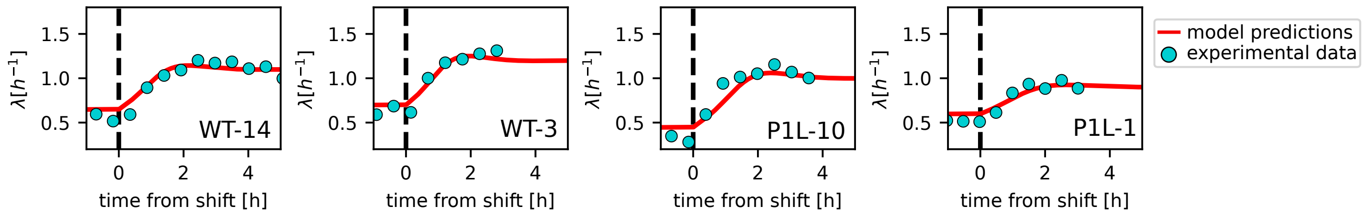


Supplementary Figure 12: **Batch nutrient upshift that do not show overshoots, but agree with model predictions.** In all panels, the solid red line represents the model prediction (from simulations), with the initial and final growth rates adjusted to align with the experimental data. The blue circles indicate the average growth rates from the technical replicates of each experiment that successfully passed the filtering for the two criteria illustrated in the Methods section: stationary growth and coherence between technical replicates. The label in the bottom right corner specifies the strain and the index of the biological replicate. This figure includes plots from all experiments that did not show a clear overshoot, but were consistent with the theoretical predictions of a very small overshoot. The model predicts different overshoot behavior depending on initial and final growth rates, and these vary across replicates (despite the use of the same medium and the stationarity of the growth).

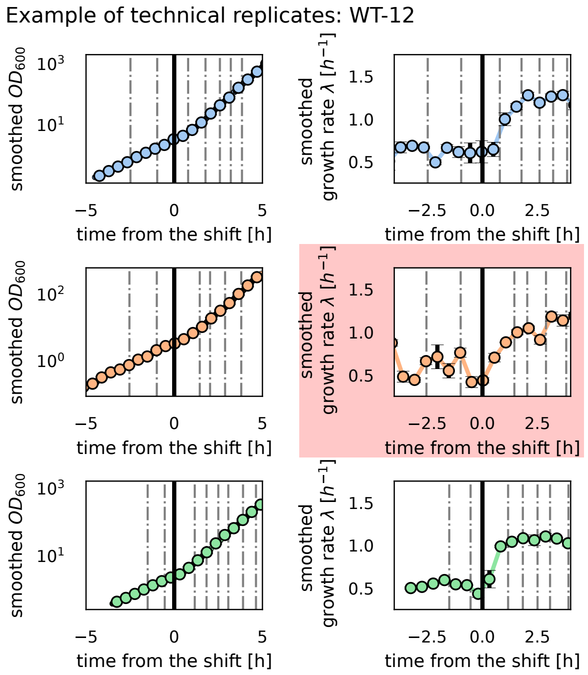


Supplementary Figure 13: **Technical replicates for wild-type biological replicate number 12, shown as an example.** The vertical dashed line corresponds to dilution points (see Methods), while the vertical solid line corresponds to the time of the shift. The left panels display the optical density ($OD_{600}$) measurements from three technical replicates, with data smoothed and patched as described in the Methods section. Circles represent 30-minute binned averages of the original time series. The right panels show the corresponding growth rates ($\lambda$) calculated from the smoothed-patched $OD_{600}$ values, also binned at 30-minute intervals. Error bars correspond to the standard deviation calculated on the bin. In calculating the average growth rate presented in Supplementary Fig. 10, we applied the selection criteria detailed in the Methods section. For this specific case, the second technical replicate (shown in red) was excluded from analysis due to unstable growth rate before the shift.

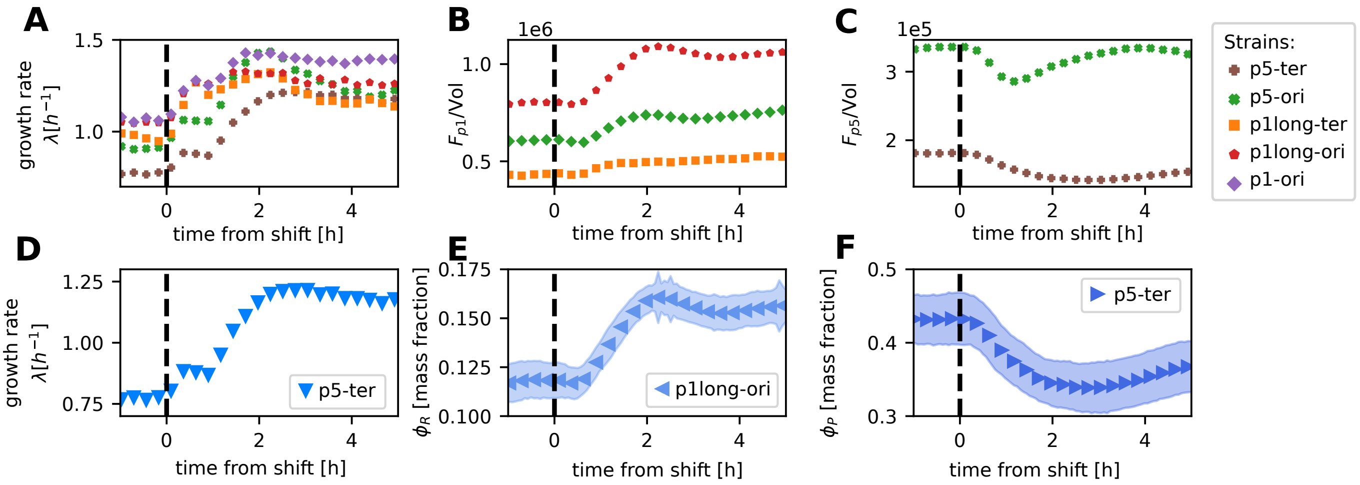


Supplementary Figure 14: **Time series of the shift dynamics for the different strains utilized in Panlilio et al. 2021** [1]. **A-C:** data from all the five studied strains, averaged across experimental replicates. **D-F:** time series used in this work. The first row shows the shift time series of the growth rate (**A**), the fluorescence signal associated with the ribosomal promoter (p1 and p1long) divided by the volume (**B**) and the fluorescence over volume for the constitutive promoter p5 (**C**). The second row shows the same data, but just for the selected strains. Panels **E** and **F** also show the standard deviation of the population average. The chosen growth rate is from p5-ter because we find it consistent with the first growth law. The fluorescent signal of the ribosomal promoter is p1long-ori because is the only one that displays a post-shift steady state. The fluorescent signal of the constitutive promoter is p5-ter because is consistent with the constraint $\phi_P + \phi_R = 0.55$. Despite the differences in growth dynamics across replicates and strains, panels A and B show that all the experiments show a coherent overshoot in growth rate and ribosomal sector (proxied by fluorescence of the P1 promoter) around 2h, displaying a remarkable consistency.

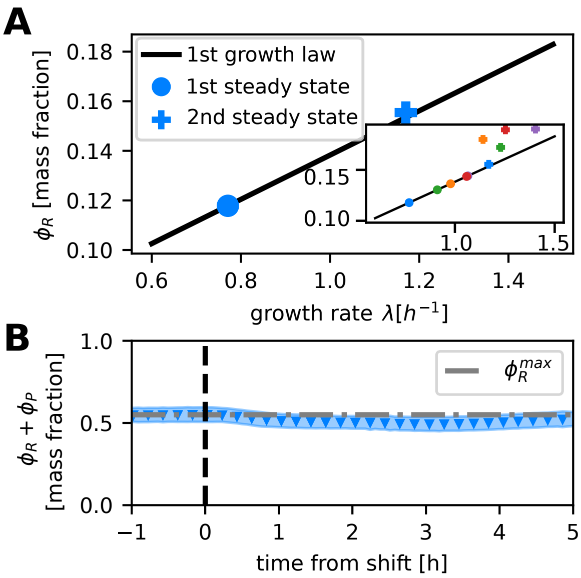


Supplementary Figure 15: **The fluorescent reporters from Panlilio et al. consistently obey the proteome sector relationships. A:** steady states for the pair of time series $(\lambda^{\mathrm{p5-ter}}, \phi_{\mathrm{R}}^{\mathrm{p1long-ori}})$. Once fixed the conversion factor $k_{\mathrm{conv,R}}$ using the data from the pre-shift steady state (circle) the post-shift steady state (plus sign) automatically obeys the first growth law (black line [2]), as expected. The inset shows the performance of the other pairs, obtained by pairing up $\phi_{\mathrm{R}}^{\mathrm{p1long-ori}}$ with growth rate from the other remaining strains The symbols are blue for $(\lambda^{\mathrm{p5-ter}}, \phi_{\mathrm{R}}^{\mathrm{p1long-ori}})$, and other colors correspond to the other pairs. **B:** once fixed the conversion factor for $\phi_{\mathrm{P}}$ using the pre-shift conditions, the constraint $\phi_{\mathrm{R}} + \phi_{\mathrm{P}} = 0.55$ is satisfied throughout the shift. The results shown in these two panels suggest that the fluorescent reporters can be a robust proxy for the ribosomal and constitutive sector size. Original data from ref. [1].

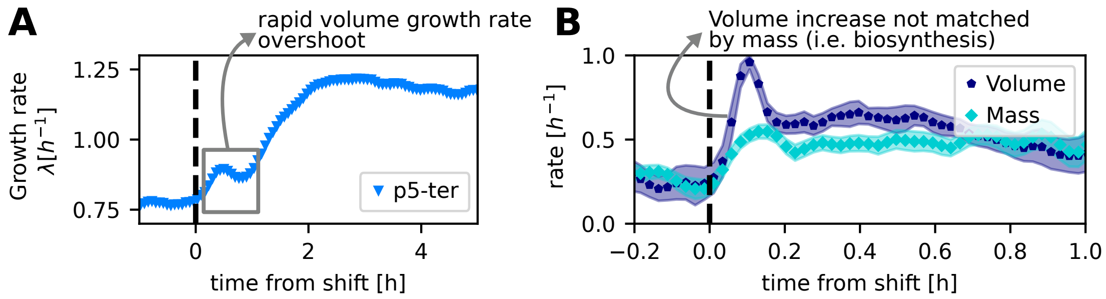


Supplementary Figure 16: **The rapid growth rate overshoot in the data from Panlilio et al. may be linked with a cell-density unbalance. A:** shift dynamics of the volume growth rate of the strain p5-ter [1], The grey box highlights the fast rapid overshoot of this quantity. **B:** Replotted data from ref. [44] form a similar amino-acid upshift (from minimal medium+mannose to minimal medium+glucose+casaminoacids) on agar pads, monitoring accumulation rates of single-cell volume and mass following a shift. Panel B shows that following a shift, the volume and the mass display very different dynamics and the cells become diluted. Indeed, cell volume increases 15 % faster than mass in the first 50 min [44]. This dilution, possibly of osmotic origin, could be the reason why the rapid overshoot in the volume growth rate is not matched by an overshoot in protein production, as monitored by the fluorescent reporters.

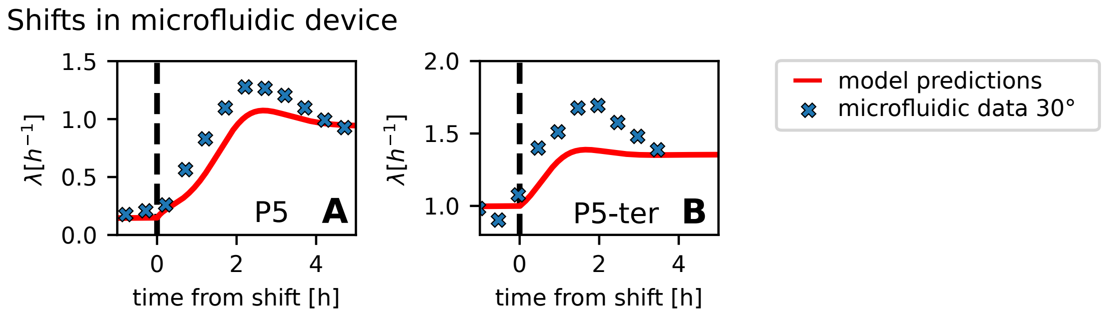


Supplementary Figure 17: **Additional overshoots observed in microfluidic experiments. A-B:** The figure shows the relaxation of the growth rate to the new steady-state for additional microfluidic experiments performed at 30°C. In both panels, continuous red lines represent the results of a simulation of the model. The points are binned averages with a window of 40min. The label in the bottom right corner of the plot indicates the strain used (see methods and ref.[1, 45]). In this case, the overshoot is much more pronounced than the one predicted by the model in these conditions.

### SUPPLEMENTARY REFERENCES

---


[1] M. Panlilio, J. Grilli, G. Tallarico, I. Iuliani, B. Sclavi, P. Cicuta, and M. C. Lagomarsino, Threshold accumulation of a constitutive protein explains E. coli cell-division behavior in nutrient upshifts, Proceedings of the National Academy of Sciences **118**, 10.1073/pnas.2016391118 (2021).

[2] D. W. Erickson, S. J. Schink, V. Patsalo, J. R. Williamson, U. Gerland, and T. Hwa, A global resource allocation strategy governs growth transition kinetics of Escherichia coli, Nature **551**, 119 (2017).

[3] L. U. Magnusson, A. Farewell, and T. Nyström, ppGpp: a global regulator in Escherichia coli, Trends in Microbiology **13**, 236 (2005).

[4] R. Balakrishnan, M. Mori, I. Segota, Z. Zhang, R. Aebersold, C. Ludwig, and T. Hwa, Principles of gene regulation quantitatively connect DNA to RNA and proteins in bacteria, Science **378**, 10.1126/science.abk2066 (2022).

[5] A. G. Marr, Growth rate of Escherichia coli, Microbiological Reviews **55**, 316 (1991).

[6] C. Wu, R. Balakrishnan, N. Braniff, M. Mori, G. Manzanarez, Z. Zhang, and T. Hwa, Cellular perception of growth rate and the mechanistic origin of bacterial growth law, Proceedings of the National Academy of Sciences **119**, 10.1073/pnas.2201585119 (2022).

[7] I. Shachrai, A. Zaslaver, U. Alon, and E. Dekel, Cost of unneeded proteins in E. coli is reduced after several generations in exponential growth, Molecular Cell **38**, 758 (2010).

[8] E. Bosdriesz, D. Molenaar, B. Teusink, and F. J. Bruggeman, How fast-growing bacteria robustly tune their ribosome concentration to approximate growth-rate maximization, The FEBS Journal **282**, 2029 (2015).

[9] M. Scott, S. Klumpp, E. M. Mateescu, and T. Hwa, Emergence of robust growth laws from optimal regulation of ribosome synthesis, Molecular Systems Biology **10**, 747 (2014).

[10] M. Scott, C. W. Gunderson, E. M. Mateescu, Z. Zhang, and T. Hwa, Interdependence of cell growth and gene expression: Origins and consequences, Science **330**, 1099 (2010).

[11] H. Bremer and P. P. Dennis, Modulation of chemical composition and other parameters of the cell at different exponential growth rates, EcoSal Plus **3**, 10.1128/ecosal.5.2.3 (2008).

[12] C. You, H. Okano, S. Hui, Z. Zhang, M. Kim, C. W. Gunderson, Y.-P. Wang, P. Lenz, D. Yan, and T. Hwa, Coordination of bacterial proteome with metabolism by cyclic AMP signalling, Nature **500**, 301 (2013).

[13] R. Hermsen, H. Okano, C. You, N. Werner, and T. Hwa, A growth-rate composition formula for

the growth of e. coli on co-utilized carbon substrates, Molecular Systems Biology **11**, 801 (2015).
[14] N. M. Belliveau, G. Chure, C. L. Hueschen, H. G. Garcia, J. Kondev, D. S. Fisher, J. A. Theriot, and R. Phillips, Fundamental limits on the rate of bacterial growth and their influence on proteomic composition, Cell Systems **12**, 924 (2021).
[15] X. Dai, M. Zhu, M. Warren, R. Balakrishnan, V. Patsalo, H. Okano, J. R. Williamson, K. Fredrick, Y.-P. Wang, and T. Hwa, Reduction of translating ribosomes enables Escherichia coli to maintain elongation rates during slow growth, Nature Microbiology **2**, 10.1038/nmicrobiol.2016.231 (2016).
[16] L. Calabrese, J. Grilli, M. Osella, C. P. Kempes, M. C. Lagomarsino, and L. Ciandrini, Protein degradation sets the fraction of active ribosomes at vanishing growth, PLOS Computational Biology **18**, e1010059 (2022).
[17] G. Chure and J. Cremer, An optimal regulation of fluxes dictates microbial growth in and out of steady-state, eLife **12**, 10.7554/elife.84878 (2023).
[18] K. Potrykus, H. Murphy, N. Philippe, and M. Cashel, ppGpp is the major source of growth rate control in E. coli, Environmental Microbiology **13**, 563 (2010).
[19] H. Bremer and P. Dennis, Feedback control of ribosome function in Escherichia coli, Biochimie **90**, 493 (2008).
[20] L. Calabrese, L. Ciandrini, and M. C. Lagomarsino, How total mRNA influences cell growth, biorxiv 10.1101/2023.03.17.533181 (2023).
[21] M. Nomura, J. L. Yates, D. Dean, and L. E. Post, Feedback regulation of ribosomal protein gene expression in Escherichia coli: structural homology of ribosomal RNA and ribosomal protein MRNA., Proceedings of the National Academy of Sciences **77**, 7084 (1980).
[22] B. J. Paul, W. Ross, T. Gaal, and R. L. Gourse, rRNA transcription in Escherichia coli, Annual Review of Genetics **38**, 749 (2004).
[23] J. J. Lemke, P. Sanchez-Vazquez, H. L. Burgos, G. Hedberg, W. Ross, and R. L. Gourse, Direct regulation of Escherichia coli ribosomal protein promoters by the transcription factors ppGpp and DksA, Proceedings of the National Academy of Sciences **108**, 5712 (2011).
[24] P. P. Dennis, M. Ehrenberg, and H. Bremer, Control of rRNA synthesis in Escherichia coli: a systems biology approach, Microbiology and Molecular Biology Reviews **68**, 639 (2004).
[25] S. Kostinski and S. Reuveni, Ribosome composition maximizes cellular growth rates in e. coli, Physical Review Letters **125**, 028103 (2020).
[26] A. Roy, D. Goberman, and R. Pugatch, A unifying autocatalytic network-based framework for bacterial growth laws, Proceedings of the National Academy of Sciences **118**, 10.1073/pnas.2107829118 (2021).

[27] M. J. Pine, Regulation of intracellular proteolysis in Escherichia coli, Journal of Bacteriology **115**, 107 (1973).

[28] T. M. Wendrich, G. Blaha, D. N. Wilson, M. A. Marahiel, and K. H. Nierhaus, Dissection of the mechanism for the stringent factor RelA, Molecular Cell **10**, 779 (2002).

[29] M. Scott and T. Hwa, Shaping bacterial gene expression by physiological and proteome allocation constraints, Nature Reviews Microbiology **21**, 327 (2022).

[30] U. Alon, *An Introduction To Systems Biology : Design Principles Of Biological Circuits* (CRC Press, 2019).

[31] W.-H. Lin, E. Kussell, L.-S. Young, and C. Jacobs-Wagner, Origin of exponential growth in nonlinear reaction networks, Proceedings of the National Academy of Sciences **117**, 27795 (2020).

[32] N. Rosenfeld, M. B. Elowitz, and U. Alon, Negative autoregulation speeds the response times of transcription networks, Journal of Molecular Biology **323**, 785 (2002).

[33] J. Wang, Y. Jiang, M. Vincent, Y. Sun, H. Yu, J. Wang, Q. Bao, H. Kong, and S. Hu, Complete genome sequence of bacteriophage t5, Virology **332**, 45 (2005).

[34] M. Maeda, T. Shimada, and A. Ishihama, Strength and regulation of seven rRNA promoters in Escherichia coli, PLOS ONE **10**, e0144697 (2015).

[35] X. Zhang and H. Bremer, Control of the Escherichia coli rrnB p1 promoter strength by ppGpp, Journal of Biological Chemistry **270**, 11181 (1995).

[36] B. Gummesson, M. Lovmar, and T. Nyström, A proximal promoter element required for positive transcriptional control by guanosine tetraphosphate and DksA protein during the stringent response, Journal of Biological Chemistry **288**, 21055 (2013).

[37] Q. Zhang, E. Brambilla, R. Li, H. Shi, M. C. Lagomarsino, and B. Sclavi, A decrease in transcription capacity limits growth rate upon translation inhibition, mSystems **5**, 10.1128/msystems.00575-20 (2020).

[38] L. Rao, W. Ross, J. Appleman, T. Gaal, S. Leirmo, P. J. Schlax, M. Record, and R. L. Gourse, Factor independent activation of rrnB p1, Journal of Molecular Biology **235**, 1421 (1994).

[39] P. Hengen, Information analysis of fis binding sites, Nucleic Acids Research **25**, 4994 (1997).

[40] W. Ross, J. F. Thompson, J. T. Newlands, and R. L. Gourse, E.coli fis protein activates ribosomal RNA transcription in vitro and in vivo., The EMBO Journal **9**, 3733 (1990).

[41] C. A. Ball, R. Osuna, K. C. Ferguson, and R. C. Johnson, Dramatic changes in fis levels upon nutrient upshift in Escherichia coli, Journal of Bacteriology **174**, 8043 (1992).

[42] M. M. C. A. Panlilio, *Non-equilibrium growth laws: growth and gene expression dynamics in Escherichia coli under a nutritional upshift*, Ph.D. thesis, Department of Physics, University of

Cambridge (September 2020).

[43] S. Hui, J. M. Silverman, S. S. Chen, D. W. Erickson, M. Basan, J. Wang, T. Hwa, and J. R. Williamson, Quantitative proteomic analysis reveals a simple strategy of global resource allocation in bacteria, Molecular Systems Biology **11**, 784 (2015).

[44] E. R. Oldewurtel, Y. Kitahara, and S. van Teeffelen, Robust surface-to-mass coupling and turgor-dependent cell width determine bacterial dry-mass density, Proceedings of the National Academy of Sciences **118**, 10.1073/pnas.2021416118 (2021).

[45] I. Iuliani, G. Mbemba, M. C. Lagomarsino, and B. Sclavi, Direct single-cell observation of a key Escherichia coli cell-cycle oscillator, Science Advances **10**, 10.1126/sciadv.ado5398 (2024).
